# An AI-based and coding-free integration for forest Leaf Area Index calculation

**DOI:** 10.1101/2025.07.24.666563

**Authors:** Tao Ma, Minxue Tang, Fangni Liu, Akwasi Duah-Gyamfi, Stephen Adu-Bredu, Mark Arcebal K. Naive, Imma Oliveras Menor, Sam Moore, Zhiyuan Zhang, Sophie Fauset, William Hagan Brown, Jill Ashley, Scott Davidson, Forzia Ibrahim, Emmanuel Amponsah Manu, Shalom D. Addo-Danso, Yadvinder Malhi, Cecilia A. L. Dahlsjo, Huanyuan Zhang-Zheng

**Affiliations:** College of Resources and Environmental Sciences, China Agricultural University, Beijing, China; Environmental Change Institute, School of Geography and the Environment, University of Oxford, Oxford, United Kingdom; Georgina Mace Centre for the Living Planet, Department of Life Sciences, Imperial College London, Silwood Park Campus, Buckhurst Road, Ascot, United Kingdom; Forestry Research Institute of Ghana, Council for Scientific and Industrial Research, Kumasi, Ghana; Center for Integrative Conservation and Yunnan Key Laboratory for Conservation of Tropical Rainforests & Asian Elephants, Xishuangbanna Tropical Botanical Garden, Chinese Academy of Sciences, Mengla, Yunnan 666303, China; University of Chinese Academy of Sciences, Beijing 100049, China; AMAP – botAnique et Modélisation de l’Architecture des Plantes et des Végétations, Université de Montpellier, CIRAD, CNRS, INRAE, IRD, Montpellier CEDEX 5, France; College of Resources and Environment, Fujian Agriculture and Forestry University, Fuzhou, Fujian 350002, China; School of Geography, Earth and Environmental Sciences, University of Plymouth, Plymouth, UK; Département des sciences biologiques, Université du Québec à Montréal, Montréal, Canada; Leverhulme Centre for Nature Recovery, University of Oxford, Oxford, United Kingdom

**Keywords:** Leaf Area Index (LAI), digital hemispherical photography (DHP), machine-learning segmentation, ilastik, automated workflow, seasonal dynamics

## Abstract

Seasonal and spatial variations in leaf area index (LAI) are challenging to detect in tropical forests due to dynamic lighting conditions and the subtle differences in the variation. Many existing LAI software tools offer one-click processing of all images through auto-threshold segmentation (e.g., HemispheR, HemiPy and Hemisfer), but they produce results with large discrepancies. Some software (e.g. CAN-EYE) requires manual tuning of each image, making large-scale analysis impractical.
We analysed 19,000 images from four tropical forest subtypes and found that using coding-free AI software to process hemispherical images can significantly improve the consistency of leaf-sky segmentation, thereby enhancing LAI outcomes.
The results show that replacing the auto-threshold with AI substantially reduced inter-software disagreement and delineated correct seasonal and spatial patterns. CAN-EYE was able to identify seasonal patterns but produced less accurate results than the CAN-EYE-AI integrated approach due to subjective user bias.
The high consistency achieved through AI integration enables reliable cross-site and cross-operator comparisons. As users can customise the AI model according to local images and combine the AI model with other LAI software, our integrated, affordable, and coding-free method offers wide applicability and high consistency of LAI measurements, facilitating the advancement of tropical forest monitoring and research.

**Data/Code for peer review statement:** One of the key features of this method is ‘coding-free’. The method is explained in Protocolv20251118.docx. We have uploaded R codes for drawing figures in a zip pack.

These codes and the protocol will be deposited in the Zenodo (or figshare) database under accession link **<u>[TBC]</u>**. Since Zenodo allows authors to archive updated versions after publication, we may update the protocol by uploading a revised version to Zenodo. Please check the Zenodo archive for any new versions. In the protocol, we note that users can use **Image_conversion_20220407.m** and **lets_change_values.R** instead of the ‘Renormalise’ function of ilastik to modify values in the classification output images. These codes are not essential for users following our protocol, but could be useful for integrating ilastik with other LAI software not covered in this paper. Additionally, the protocol mentions that **Gather_LAI_fapar_from_caneye.R** can be used to consolidate output Excel files, eliminating the need to manually open each file. Field measurements of LAI and GCC are available on request.

## Introduction

The Leaf Area Index (LAI) is widely used in climate change models, global carbon cycle assessments, and ecosystem restoration studies (Hazarika et al., 2005; Sumida et al., 2018; Veryard et al., 2023). In the context of global climate change, accurately capturing spatiotemporal variations in LAI (Jiang et al., 2017) has become essential to research in vegetation ecology and environmental science (Houborg et al., 2016; Ma et al., 2023). In tropical forests, LAI exhibits subtle, but important, spatial and temporal variations (de Wasseige et al., 2003; Malhado et al., 2009) with values differing among forest subtypes, ranging from wet evergreen to semi-deciduous and dry forests (Middinti et al., 2017). While seasonality in wet evergreen tropical forests may not be visually apparent, these minor seasonal variations (phenology) play a critical role in tropical forest functioning (Wu et al., 2017, 2016). Accurately detecting subtle fluctuations in LAI has long posed a significant challenge (Li et al., 2024; Yan et al., 2019; Zou et al., 2023).

Currently, most *in situ* LAI is measured by DSLR digital hemispherical photography (DHP), which uses fisheye lenses to capture canopy hemispheres, which are segmented into leaf and sky pixels to derive gap fraction for LAI estimation (Yang et al., 2023). To determine temporal or spatial LAI patterns, segmentation must be both accurate and consistent, yet it has long been recognised as the main methodological bottleneck (Jonckheere et al., 2004). Some tools (e.g., CAN-EYE) rely on manual segmentation, which is time-consuming and introduces subjective error, undermining cross-ecosystem comparability (Fang et al., 2014; Jonckheere et al., 2005a). Threshold choice can vary strongly among operators, generating substantial subjectivity that compromises cross-site studies and even multi-year monitoring at the same site when personnel change (Jonckheere et al., 2005a; Pueschel et al., 2012).

Some software tools allow ‘one-click’ batch processing by using automatic threshold detection. Based on 300+ hemispherical photos of Belgian temperate forests, researchers comprehensively assessed 35 thresholding algorithms (Jonckheere et al., 2005a). Compared with manual thresholding, these approaches significantly reduced subjectivity and were widely regarded as a major breakthrough at the time. However, the study also noted that these methods face challenges in dense canopies or under uneven illumination, challenges that are particularly common in tropical forests. Moreover, they also reported significant variation in segmentation outcomes among auto-threshold methods, which contributes to inter-software discrepancy in LAI. Such inconsistencies have been frequently reported in the literature (de Wasseige et al., 2003; Jarčuška et al., 2010; Jonckheere et al., 2005a; Liu et al., 2021; Pastorella and Paletto, 2013; Woodgate et al., 2015), affecting the reliability of LAI measurements. Auto-threshold techniques remain widely used in open-source LAI packages such as HemispheR and HemiPy, which have shown reliable or promising performance in temperate forests, but their applicability to tropical forests with denser canopies and highly heterogeneous light conditions remains underexplored (Chianucci and Macek, 2023). Given the recent leap in the development of AI image processing, we hypothesise that integrating LAI packages with AI image processing tools could potentially improve their LAI outcome. Our exploration does not seek to implement the latest and most powerful AI package, but rather utilises a free, no-coding-required AI image processing software (ilastik) (Berg et al., 2019) which is accessible to a wide range of users.

In this paper, we use ilastik to replace manual or auto-thresholding in the leaf-sky segmentation step of several LAI software tools(HemispheR, HemiPy, Hemisfer, and CAN-EYE), while keeping the LAI calculation based on gap fraction unchanged. Particularly, the aims are to investigate (1) whether AI improves CAN-EYE’s results, as the AI model eliminates subjectivity in manual image segmentation, and (2) whether replacing auto-thresholding with AI improves LAI outcomes in HemispheR, HemiPy, and Hemisfer. The testing dataset includes 19,044 hemispherical photos from four tropical forest types in Ghana, Africa. Since the magnitude of tropical forest LAI could not be ascertained without destruction, we focused on the seasonal and spatial variation of LAI, which was validated by independent measurements and literature knowledge. Additionally, we provide a step-by-step guide on how to integrate the AI software with the four LAI software tools.

## Technical workflow

### Overview

Our workflow consists of two major steps, which separate image segmentation from LAI computation (Figure 1): an ilastik-based segmentation step and a software-specific calculation step. In the first step, users train a supervised ilastik classifier on labelled canopy/sky pixels and apply the model to batch-segment hemispherical photographs. In the second step, the resulting binary images are imported into existing LAI tools (e.g., CAN-EYE, HemispheR, HemiPy and Hemisfer), where LAI is computed using each software’s native routines.

**Figure 1.**
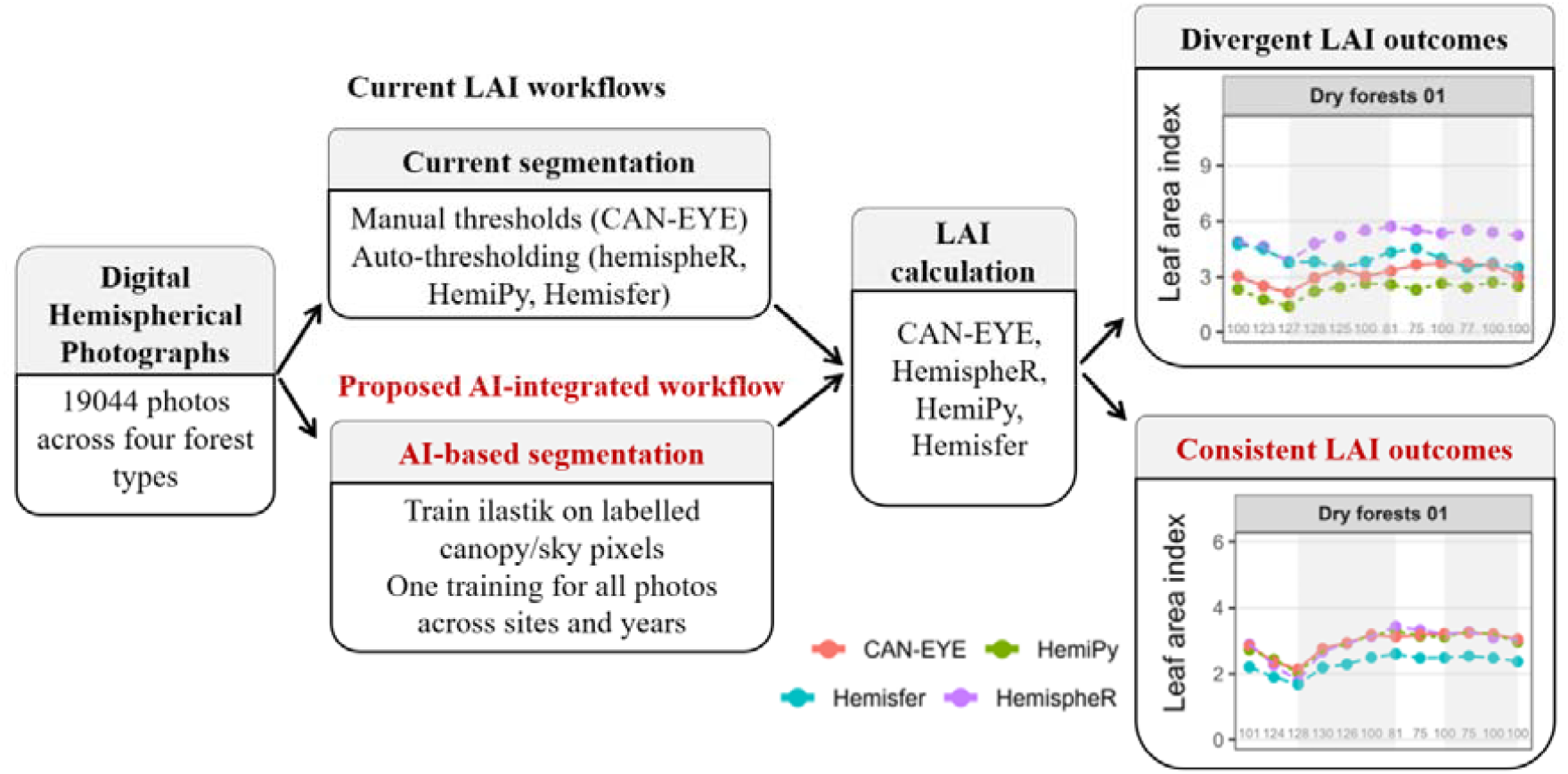
Overview of the AI-integrated workflow for deriving LAI from digital hemispherical photographs. The workflow decouples canopy–sky segmentation (performed in ilastik) from LAI calculation (performed in CAN-EYE, HemispheR, HemiPy or Hemisfer), and is illustrated using our case study dataset.

### Protocol availability and accessibility

To help future researchers, we have provided a step-by-step guide showing how to integrate ilastik with LAI estimation software (see Supplementary Materials). The protocol consists of two key steps: 1) image segmentation (using ilastik); 2) LAI calculation (using HemispheR, HemiPy, and CAN-EYE). We focused on providing additional details for CAN-EYE because HemispheR and HemiPy require programming experience, whereas Hemisfer requires a paid subscription. Our goal is to maximise accessibility by ensuring the protocol is free of charge and requires no coding knowledge. To improve clarity, the protocol includes detailed annotated screenshots and step-by-step instructional videos (see Supplementary Video 1). The protocol documents all parameters for both ilastik and CAN-EYE used for hemispherical photo processing. Additionally, it is compatible with budget-friendly laptops that lack GPU acceleration.

### AI-based image segmentation using ilastik

We used ilastik, an interactive image analysis software that utilises machine learning techniques (Berg et al., 2019), to process hemispherical photos to classify canopy and sky pixels, producing black-and-white images. Two AI models were trained on images with contrasting canopy conditions: 1) ANK and BOB sites (both with closed canopies), and 2) the KOG site, where the sky background can be heterogeneous. For each model, we randomly selected twenty fisheye photographs across plots and seasons. Pixel annotation was performed using the Pixel Classification workflow in ilastik, with manual labelling of leaf and sky pixels. The software then learned from these annotations to build two classification models, which were applied in batch mode to process all images. Further details of the image processing steps are provided in the protocol (in the supplementary materials and on Zenodo). In the case study, we compared two versions of hemispherical photos: (1) raw colour photos (without AI) and (2) black-and-white photos from *ilastik* (AI integration). We grouped photos into plot-month packages; each package contained 25 photos (representing 25 image acquisition points within each plot).

### LAI calculation using HemispheR, HemiPy and Hemisfer

Raw colour photos and black-and-white transformed photos were processed using HemispheR, HemiPy and Hemisfer, which generated LAI results both with and without AI. The same parameters were used for both sets of photos to ensure comparability of results. Explanations of each parameter are available in the protocol. It should be noted that HemiPy and CAN-EYE produce one LAI value per plot-month package, whereas HemispheR and Hemisfer provide one LAI value per photo. This difference in methodology resulted in variations in the number of LAI values and subsequently their standard errors.

### LAI calculation using CAN-EYE and manual segmentation experiments

All ilastik-derived black-and-white images were analysed in CAN-EYE following the protocol. Because CAN-EYE lacks auto-thresholding, raw colour images require manual sky –canopy classification; processing the full dataset was impractical, so we analysed only raw images collected after January 2018 (a time-consuming limitation of CAN-EYE). Raw images were processed in the Color DHP (RGB) interface. For each plot–month package (25 photos) from January 2018 onwards, we allowed up to 2 minutes for manual classification, labelling sky and canopy pixels in 2– 3 representative photos, after which CAN-EYE determined a brightness threshold for the package.

Because CAN-EYE outputs can be influenced by subjective error (Jonckheere et al., 2005a), we ran two additional Color DHP experiments: (1) Unlimited time, with no constraints on the number of photos or markings, continuing until satisfactory sky–canopy accuracy was achieved; and (2) 30-second limit, in which one photo per 25-photo set was manually classified 3–4 times within exactly 30 s to represent rushed users. In all experiments, LAI was computed after segmentation. Segmentation operators did not calculate LAI, viewed only sky-percentage outputs, and were blinded to LAI values to avoid bias. We refer to the four CAN-EYE result sets as ‘CAN-EYE + AI’, ‘2 minutes’, ‘30 seconds’, and ‘unlimited time’.

## CASE STUDIES

### Case study site

West African tropical forests exhibit a well-defined ecological gradient, transitioning from coastal wet evergreen forests to savanna woodlands as they approach arid regions. Our study focused on four distinct forest subtypes: 1) three wet evergreen plots, 2) three semi-deciduous plots, 3) two dry forest plots, and 4) two woody savanna plots. Each plot covers one hectare. All plots were located within three protected reserves in Ghana (ANK, BOB, and KOG) (Table 1). All study sites experience a bimodal rainfall pattern, with the primary wet season peaking in June and a secondary wet season in October (Appendix A3 and Supplementary Figure 1). December and January are the driest months. Leaf flushing and shedding occur twice annually, corresponding to the two wet seasons (Lieberman, 1982). The study sites form part of the Global Ecosystem Monitoring (GEM) network (Malhi et al., 2021) and are registered in the ForestPlots.net database. More information about these study sites could be found in previous publications(Zhang-Zheng et al., 2024a, 2024b).

**Table 1.**
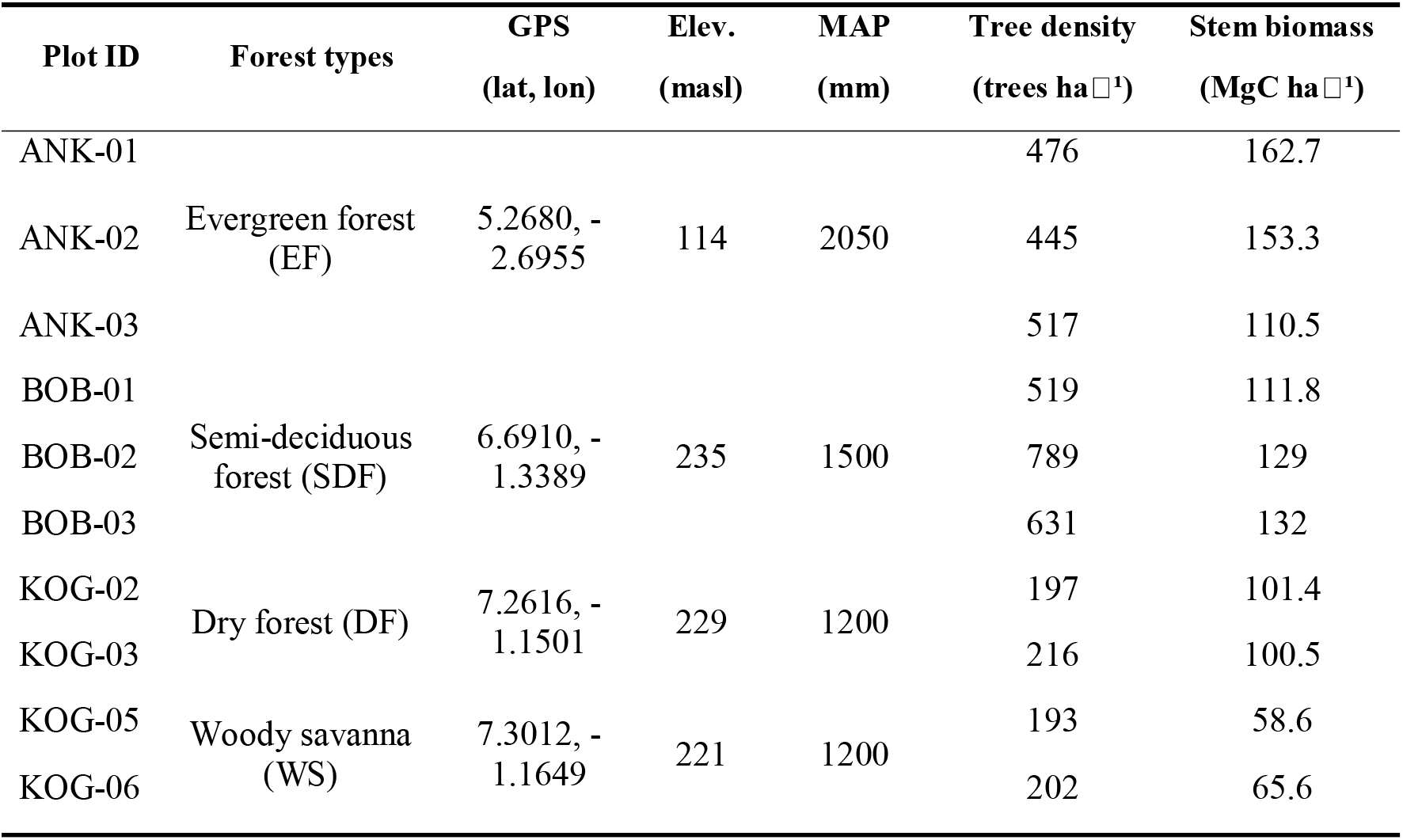
Key characteristics of study sites. We show Plot ID, Forest types, Coordinates (GPS), Elevation (elev.), Mean annual precipitation (MAP), Tree Density and Aboveground stem biomass.

We analysed 19,044 hemispherical photographs collected monthly (2012–2019) (Appendix A4) from these study sites along a rainfall seasonality gradient (Figure 2). LAI seasonality and spatial differences derived from this dataset were evaluated against independent leaf-flushing phenology from a top-down camera and published phenological descriptions for the region. Phenology and spatial variation of these study sites were described in the supplementary (Appendix A1 and A2).

**Figure 2.**
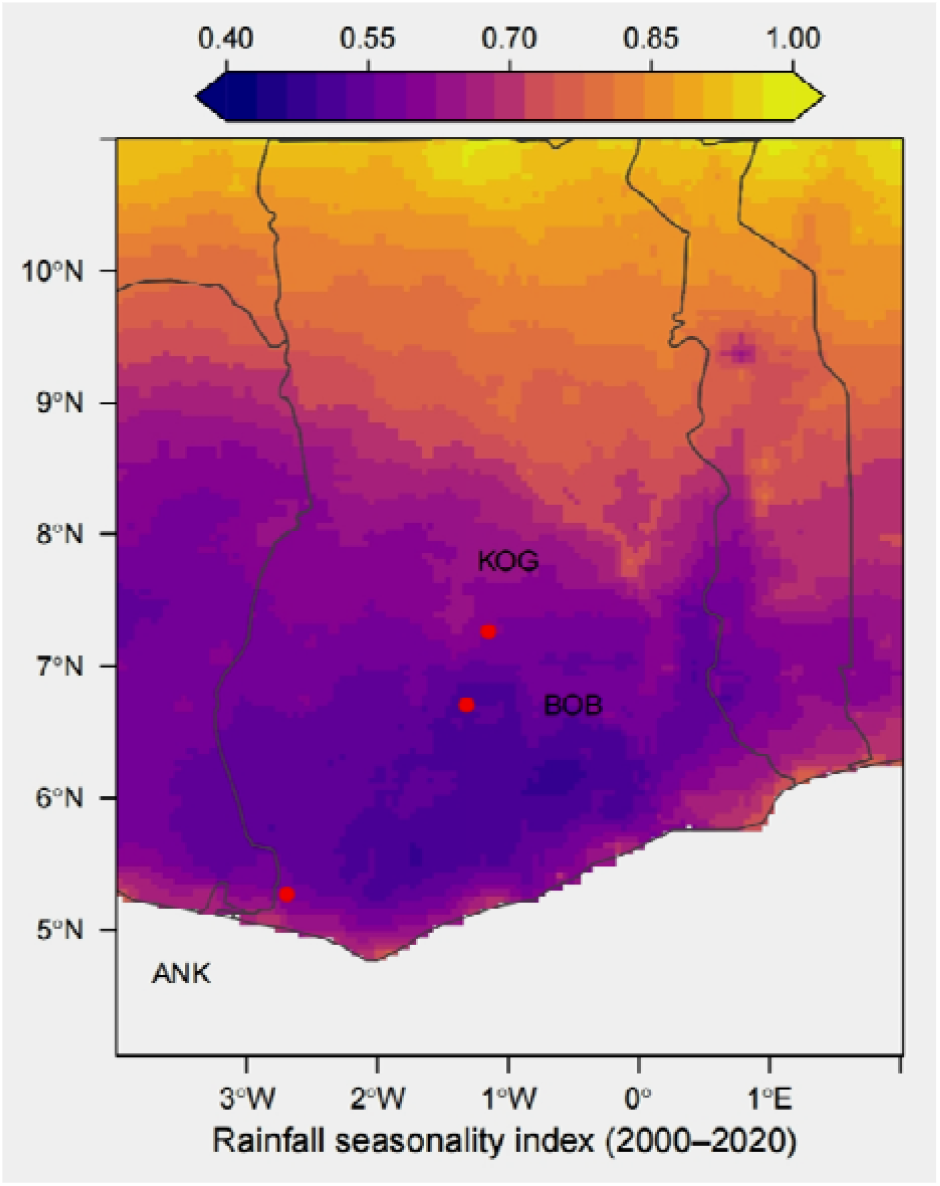
Map of the study site and rainfall seasonality index of Ghana, West Africa. The background shows the seasonality index of rainfall. Typically, values smaller than 0.2 suggest no seasonality. Values around 0.45 suggest moderate seasonality with a short drier season. Values larger than 1.0 suggest most rainfall occurs in less than 3 months (Nair et al., 2014). Abbreviations: ANK = Ankasa; KOG = Kogyae; BOB = Bobiri.

### Sensitivity of CAN-EYE LAI estimates to manual segmentation time

In traditional CAN-EYE colour image processing, LAI outcomes are affected by the time spent on manual segmentation (Figure 3). The ‘Unlimited time’ experiment does not provide the best performance, but instead consistently yields higher LAI values than other experiments (Figure 3, Supplementary Figure 8) and produces an incorrect seasonal pattern. Consequently, the ‘30 seconds’ experiment likely omitted some leaves due to time constraints, but ‘Unlimited time’ tends to classify leaves excessively, leading to an overly permissive or overly conservative threshold.

**Figure 3.**
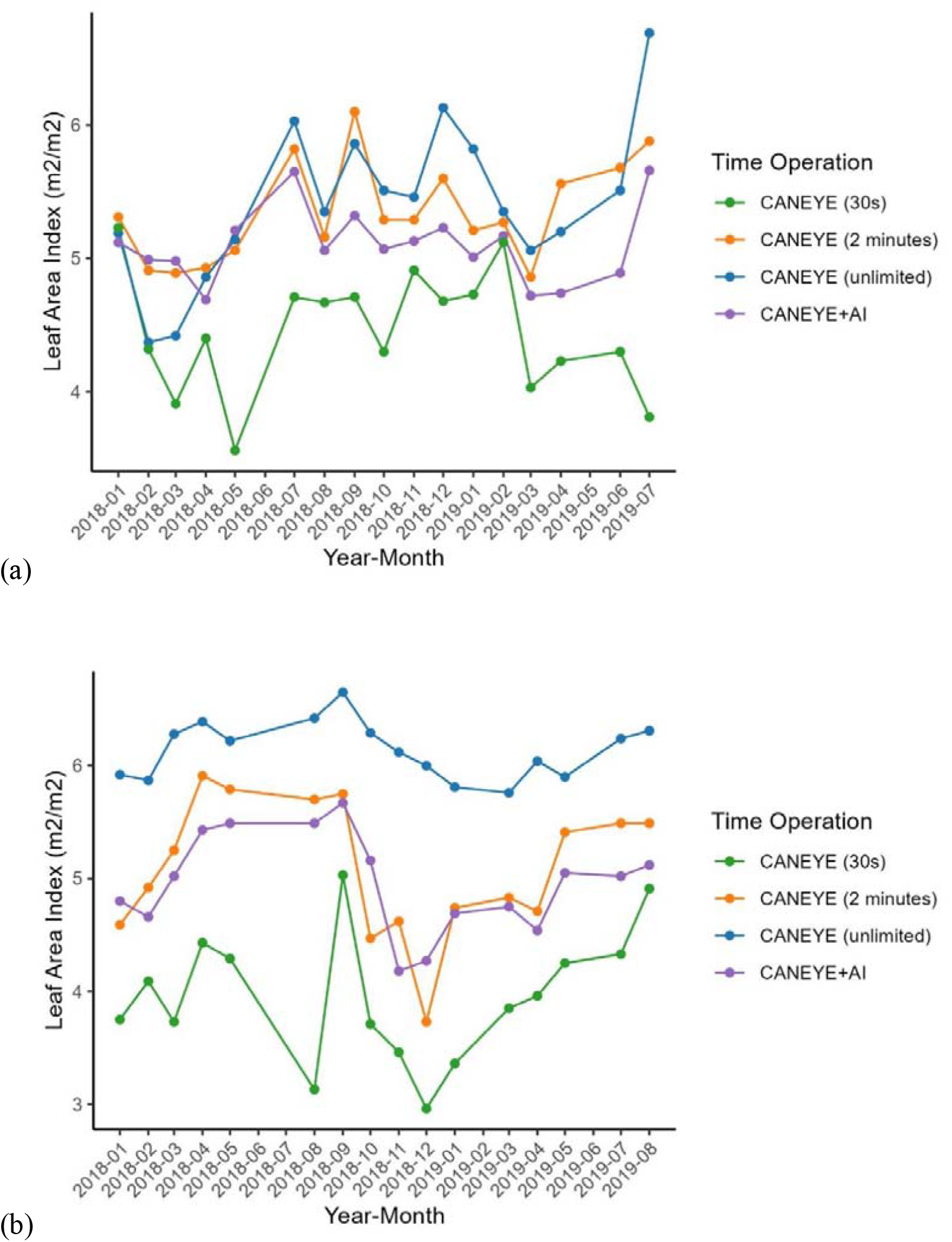
Seasonal variation of Leaf area index for different experiment settings in (a) Wet evergreen 01 ANK-01 and (b) semi-deciduous 01 BOB-01. The purple line represents the AI integration, using ilastik to process images, which has no operation time, as it only needs to be trained once. The remaining colours represent traditional manual image processing experiments (including 30 seconds, 2 minutes and unlimited time), where Manual pixel classification was conducted with different timings.

The ‘2 minutes’ method captures a seasonal trend broadly similar to ‘AI + CAN-EYE’, but its LAI values are significantly higher (Supplementary Figure 8 and Figure 9). At ANK-01, visual inspection indicates fewer leaves in April 2019 than in February 2019 (Supplementary Figure 3). This is reflected by AI + CAN-EYE, whereas CAN-EYE-only does not reproduce this contrast (Figure 3a). Moreover, the pronounced month-to-month spikes in CAN-EYE-only (2 minutes)—from July (high) to August (low) and back to a very high September—are biologically implausible, supporting the conclusion that AI + CAN-EYE provides more realistic seasonal variation.

Across other plots (Figure 4), similar spiky fluctuations occur in wet evergreen 02 and semi-deciduous 02. Overall, both approaches show broadly comparable seasonality, but AI + CAN-EYE is consistently more reasonable and aligns better with the expected seasonal pattern.

**Figure 4.**
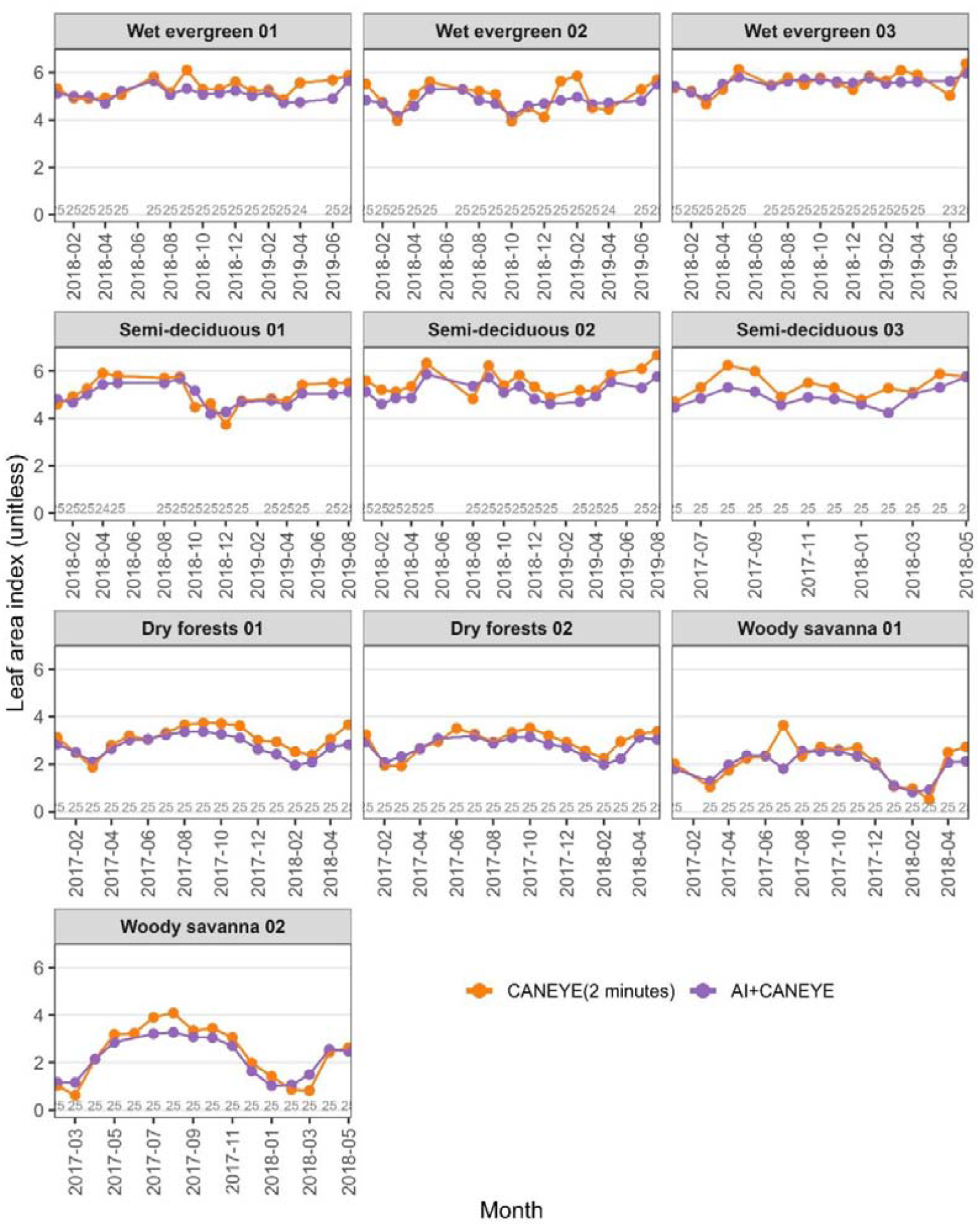
Seasonal variation of Leaf area index (LAI) for CAN-EYE with and without AI. The purple line represents the protocol which uses ilastik to process images. The orange line represents the two-minute manual processing experiment using CAN-EYE. The grey numbers at the bottom represent the number of photos behind each LAI value. Manual processing of photos is highly time-consuming, and thus, CAN-EYE (2 minutes) was conducted for the last two years only. AI+CAN-EYE in this figure also includes the last two years only for fair comparison, but AI+CAN-EYE can be run automatically for all photos, the results of which are available in Figure 6.

### Reduction of inter-software discrepancies through AI-based segmentation

When segmentation relied on each software’s original routine (i.e., without AI), LAI estimates diverged strongly among packages (Figure 5). For wet evergreen forests, mean annual LAI ranged from 2.5 to 6.5 depending on software. Seasonal patterns also disagreed, most notably at woody savanna 01: Hemisfer and HemispheR produced anomalously high February LAI (relative to other months), inconsistent with phenological expectations, whereas CAN-EYE and HemiPy showed the opposite pattern, consistent with known phenology.

**Figure 5.**
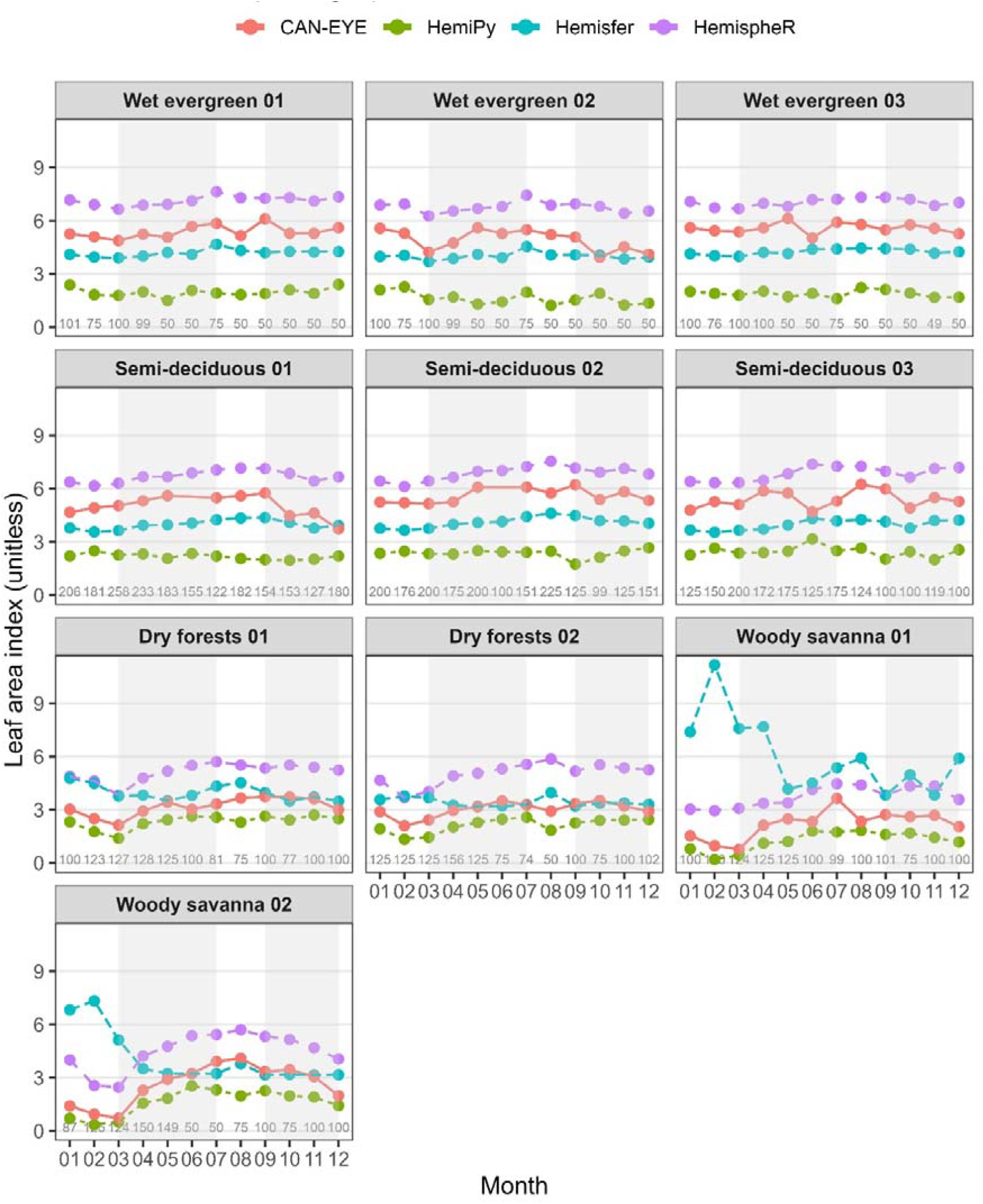
Seasonal variation of leaf area index (LAI) estimated by four software without AI-based segmentation. Monthly LAI (y-axis; unitless) is shown for each of the 10 forest plots across wet evergreen, semi-deciduous, dry forest, and woody savanna sites. The x-axis indicates month (01–12). Colours represent different DHP software: CAN-EYE (red), HemiPy (green), Hemisfer (blue), and hemispheR (purple). Each point corresponds to the monthly mean LAI for a plot, and numbers above the x-axis indicate the number of hemispherical photos used for that month. Light grey background shading marks the rainy seasons to aid visual interpretation. Each panel displays one plot.

Replacing the segmentation step with AI (ilastik) substantially improved agreement (Figure 6). Across all plots, the four packages converged on consistent seasonal trajectories. For example, at woody savanna 01 and 02, all methods identified the lowest LAI in February, with January higher than February but lower than summer values, matching the vegetation leafing phenology (Appendix A1). The AI-based results also revealed the expected increase in seasonality from wet evergreen forests to semi-deciduous and dry forests, with the strongest seasonality in woody savanna. Even in wet evergreen 01–03, a subtle but coherent seasonal signal emerged, with all packages agreeing that July had the highest LAI.

**Figure 6.**
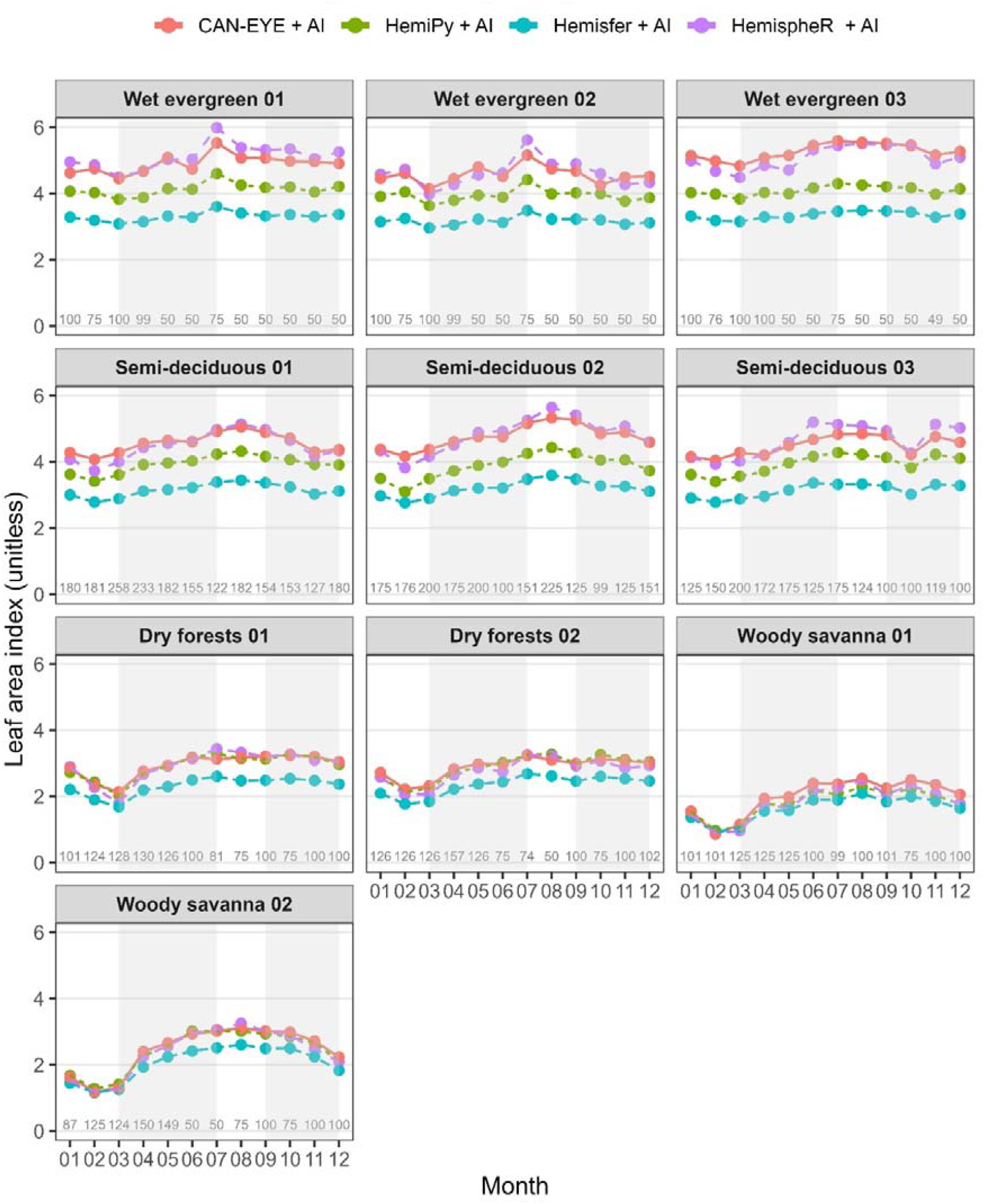
Seasonal variation of leaf area index (LAI) estimated by four software with AI-based segmentation. Same as figure 4, but here all software was fed with photos already processed by ilastik. Monthly LAI (y-axis; unitless) is shown for each of the 10 forest plots across wet evergreen, semi-deciduous, dry forest, and woody savanna sites. The x-axis indicates month (01–12). Colours represent different DHP software: CAN-EYE (red), HemiPy (green), Hemisfer (blue), and hemispheR (purple). Each point corresponds to the monthly mean LAI for a plot, and numbers above the x-axis indicate the number of hemispherical photos used for that month. Light grey background shading marks the rainy seasons to aid visual interpretation. Each panel displays one plot.

AI integration also reduced discrepancies in LAI magnitude. Differences were almost eliminated in dry forest and woody savanna plots, and reduced but not removed in wet evergreen and semi-deciduous plots (e.g., wet evergreen 01 still ranged from 3.1 to 5.0 in mean annual LAI). Overall, HemispheR produced a reasonable seasonal pattern with and without AI, whereas Hemisfer misinterpreted seasonality in dry forest/woody savanna and HemiPy in semi-deciduous forests when AI was not used. With AI integration, all packages delineated a reasonable and consistent seasonal pattern.

### Spatial patterns of LAI across forest types

Without AI segmentation, LAI magnitude and spatial patterns differed substantially among software (Figure 7b): CAN-EYE and HemispheR agreed with the expected pattern (Appendix A2), whereas HemiPy and Hemisfer did not. After AI integration, discrepancies in LAI magnitude were greatly reduced and all four packages converged on a consistent spatial pattern (Figure 7a), with the lowest LAI in woody savanna, intermediate values in dry forest, and higher LAI in wet evergreen and semi-deciduous plots. All methods also indicated slightly higher mean LAI in wet evergreen (ANK) than in semi-deciduous (BOB) plots (Figure 7), although this small difference was not apparent from field observation (Supplementary Figure 5).

**Figure 7.**
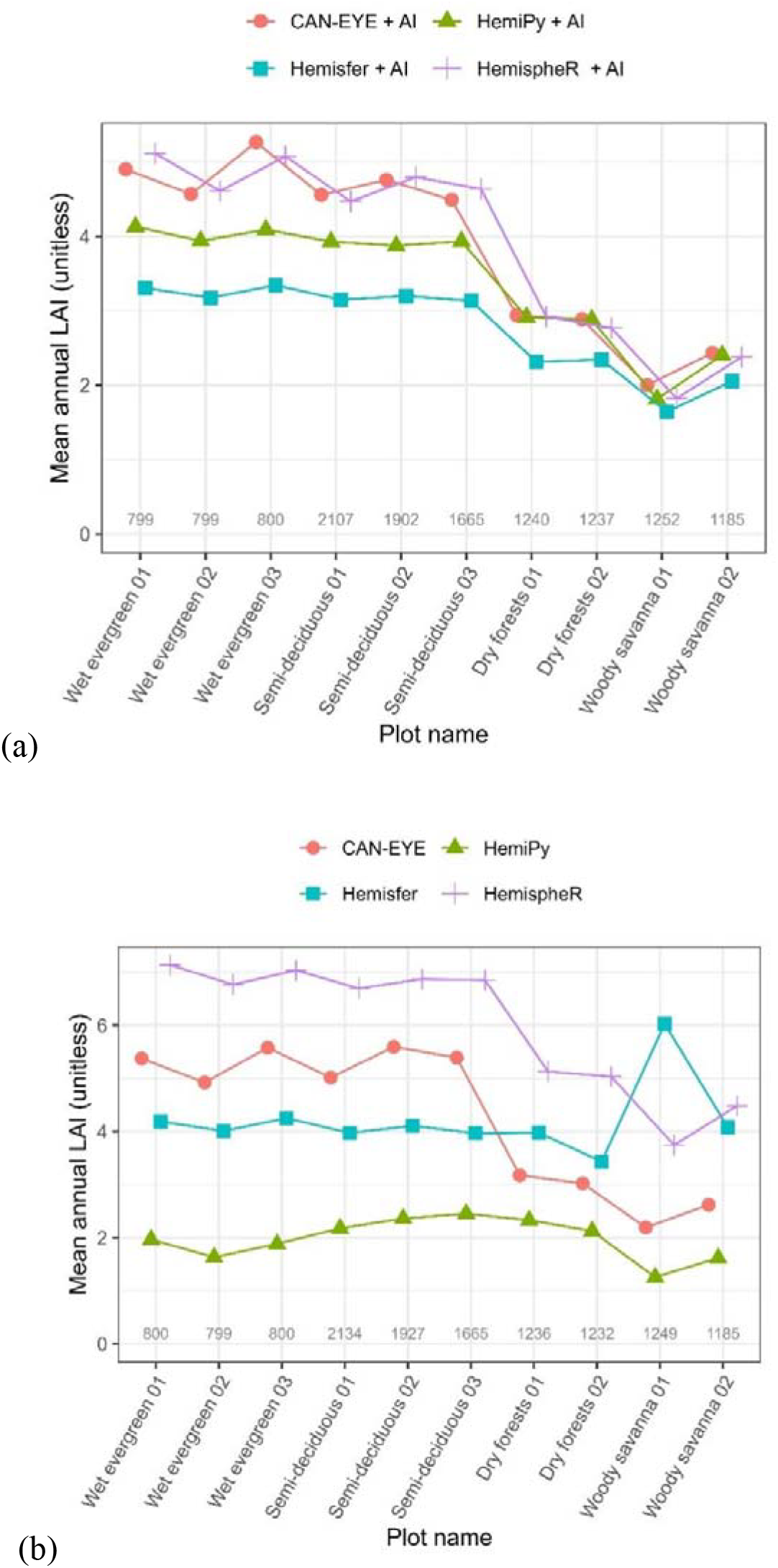
Spatial variation of mean annual leaf area index (LAI) across 10 tropical forest plots. Panel (a) shows results with AI-based image segmentation, and Panel (b) shows results without AI. In both panels, the y-axis shows mean annual LAI (unitless), and the x-axis lists the 10 forest plots spanning wet evergreen, semi-deciduous, dry forest, and woody savanna sites. Colours denote software packages: CAN-EYE (red), HemiPy (green), Hemisfer (blue), and HemispheR (purple). Each point represents the annual mean LAI at a plot. Numbers above the x-axis indicate the total number of hemispherical photographs used at each plot.

### Limitation

A limitation of the case study is the absence of an independent reference dataset for LAI, although we provide independent evidence of leaf phenology. Litterfall-based LAI can serve as ground truth in deciduous forests (e.g., benchmarking HemispheR and HemiPy) (Brown et al., 2023), but it is infeasible in evergreen tropical systems, where destructive sampling would be the only alternative. To our knowledge, all published LAI values for West African forests are derived from hemispherical photographs (Ametsitsi et al., 2020; Issifu et al., 2019); in such contexts, visual inspection and consistency with expected ecological patterns are widely used validation criteria (Jonckheere et al., 2005a). We therefore provide example photos (Supplementary) and triangulate plausibility using stem biomass (Table 1), rainfall seasonality (Supplementary Figure 1) and regional leafing phenology (Lieberman, 1982)(Supplementary Figure 11–13). Another limitation is variable exposure in field fisheye photos: although overcast conditions are recommended, this is difficult in tropical forests due to travel/time-of-day logistics, and light contrast is extreme—sky regions can remain dazzling even during rain (de Wasseige et al., 2003). With 19,044 photos, exposure noise may partly average out in mean LAI.

## Discussion and conclusion

### Comparison with manual pixel classification

Building on the results above, we show that manual pixel classification is prone to strong operator subjectivity (Figure 3). A key source of bias is a “grey zone” along leaf edges, where the human eye cannot unambiguously determine whether pixels belong to leaves or sky; operators must therefore make subjective decisions. Although sunny conditions are generally discouraged, tropical forest images can still appear very bright, with sunbeams, due to dense canopies and field logistics (Supplementary Figure 6). Under these conditions, sun-bleached leaf edges or light-green areas are easily misclassified as sky. Manual segmentation is further challenged at the woody savanna sites during the dry season, when fine leaves and twigs are difficult to segment (Supplementary Figure 6).

Accordingly, results vary among operators even for the same image, depending on annotation time, experience, and judgment criteria. Previous work has shown that human judgement is inherently subjective and can substantially affect data quality (Gilardelli et al., 2018; Jonckheere et al., 2005b). Consistent with our findings, spending more time on annotation leads operators to label more grey-zone pixels as leaves, producing progressively higher LAI values (Figure 3). In addition, CAN-EYE’s edge processing lacks precision and often introduces erroneous pixel clusters along sky boundaries (Supplementary Figure 7), which can further inflate LAI; similar artefacts have been reported previously (Duveiller and Defourny, 2010; Zhu et al., 2023).

Differences among time-limited settings can also be explained by operator strategy. Operators typically label sky first and then iteratively add leaves and branches while repeatedly checking the segmentation. Under strict time constraints (e.g., 30 s), some leaves are likely omitted, whereas extended annotation time (“Unlimited time”) promotes increasingly permissive inclusion of ambiguous pixels as leaves. This can over-include grey-zone pixels, inflate LAI, and distort seasonal trajectories, as observed in our results.

For the ilastik AI approach, the model must also place a boundary within grey zones, which may not always be exact. However, once trained, the same model is applied consistently across months and sites, substantially reducing operator-dependent variability. Automated thresholding likewise improves consistency relative to manual classification, as discussed below.

### Comparison with Automated thresholding

Previous research shows that classical automated thresholding can reduce the subjectivity of manual pixel classification (Jonckheere et al., 2005a; Nobis and Hunziker, 2005). HemispheR, HemiPy and Hemisfer all rely on automated thresholding for leaf–sky separation, yet threshold choice itself can introduce substantial uncertainty, with gap-fraction deviations of up to 17% reported among algorithms (Xie et al., 2023). A systematic assessment of 35 thresholding algorithms using 300 hemispherical photographs from Belgian deciduous and coniferous stands likewise revealed large performance differences, closely mirroring our tropical-forest results (Figure 5).

In our study, replacing the automated thresholding step in these software packages with AI-based segmentation markedly increased cross-software agreement (Figure 6 and 7). Inter-method discrepancies in hemispherical approaches are widely documented, with field studies often reporting either averaged LAI or method-specific estimates (Calders et al., 2018; Pastorella and Paletto, 2013; Thimonier et al., 2010). Our results indicate that AI segmentation can substantially reduce this source of methodological divergence. A modest residual discrepancy remained at wet evergreen and semi-deciduous sites, likely reflecting differences in gap-fraction or clumping calculations rather than segmentation per se (Liu et al., 2021).

The improved agreement likely stems from higher-quality segmentation by the AI workflow (Supplementary Figure 10). In dry-forest and woody-savanna plots, some sampling points contain large, inhomogeneous sky openings (Supplementary Figure 4), so plot–month image sets can mix open and closed canopy conditions. CAN-EYE applies a single threshold across each image set, which can perform poorly under such heterogeneity. HemispheR, Hemisfer and HemiPy use image-specific thresholds, partially alleviating this limitation, but large openings remain challenging; as noted in the HemispheR documentation (Chianucci and Macek, 2023), portions of large sky gaps can still be misclassified as leaves, potentially explaining the unrealistically high February LAI at woody-savanna site 01 (Figure 5).

By contrast, ilastik is a supervised classifier that, once trained, applies consistent rules based on colour, texture and edge information rather than relying solely on brightness histograms (Berg et al., 2019). Many thresholding methods assume two distinct brightness clusters for sky and foliage (Macfarlane et al., 2014), an assumption readily violated by illumination and field conditions, reducing robustness.

### A low-cost, coding-free, and widely applicable protocol

A key feature of our AI integration is that it is low-cost, coding-free and user-friendly, making it suitable for teams with limited resources. Because both ilastik and the LAI software in our protocol are free, the main equipment requirement is a digital camera with a fisheye lens (typically **< £1**,**000**)(Zhu et al., 2023); a smartphone with a fisheye lens can also be used if appropriate parameters are applied. ilastik implements established classical machine-learning methods and runs on a standard commercial laptop (≈ **£600**)(Kreshuk and Zhang, 2019), requiring neither a high-performance cluster nor a GPU (Sommer et al., 2011).

As shown here, the workflow can be adapted to a range of forest types and woody savannas via ecosystem-specific training, enabling spatial and temporal comparisons (Obels, 2021).

Overall, AI integration substantially improves cross-software consistency in LAI estimation and largely resolves inter-software discrepancies. The workflow offers three advantages:

1. Reliability, with consistent detection of subtle seasonal and spatial LAI variation across four tropical forest subtypes.
2. Efficiency, because a trained model can be applied across sites and years, reducing operator dependence and processing time.
3. Accessibility and cost-effectiveness, through free, intuitive software and minimal equipment requirements.

## Supporting information

Supplementary protocol

## Competing interest statement

The authors declare no competing interests.

## Inclusion & Ethics statement

Local researchers were engaged throughout the research process, with clear roles and responsibilities established, particularly through contributions from the Forestry Research Institute of Ghana. The research focused on locally relevant topics and was designed in collaboration with local partners. Capacity-building workshops were conducted during the research process. Relevant local and regional studies are cited, as evidenced by the inclusion of West Africa-specific studies. Risk management plans were developed for the fieldwork in accordance with the universities’ regulations.

## Notes

### Competing Interest Statement

The authors have declared no competing interest.

### Summary of Updates

This version of the manuscript has been substantially revised to include a new comparison with mainstream automatic thresholding software, including hemispheR, HemiPy, and Hemisfer. The revised manuscript now evaluates the proposed AI based, coding free workflow against these commonly used methods, providing a broader assessment of its performance and applicability for hemispherical image segmentation and LAI calculation. In addition, the author list has been updated to include new contributors.

