## Supplementary for "An AI-based and coding-free integration for forest Leaf Area Index calculation"

### Appendix Table of Contents

### Appendix A. Supplementary Methods

***A1.Leaf phenology at the study sites***

Increasing rainfall seasonality (Supplementary Figure 1) leads to greater deciduousness from wet to dry sites. The leafing phenology of Ghanaian forests has been documented in the literature (Lieberman, 1982). The study site in that work shared some species with ours. The main leaf-flushing period occurs from mid-February to July. The second minor leaf flushing period occurs in October and November. Leaf fall also occurs throughout the year but peaks during the major dry season from late November to March, and at the lowest level during the wet season from May to August. According to GCC at our semi-deciduous plot 02, leaf flushing occurred from December 2022 to May 2023, with some minor flushing in July and October 2023 (Supplementary Figures 11 and 12). The GCC pattern is broadly consistent with previous observations (Lieberman, 1982). It should be noted that GCC was obtained from top-canopy species not included in the observations reported in the literature (Lieberman, 1982), while the cited study (Lieberman, 1982) surveyed many lianas and lower-canopy tree species. LAI from hemispherical photos should include all canopy layers and lianas in principle.

Based on leafing phenology, the thinnest canopy occurs in February (as indicated by the number of leafless species reported in the literature (Lieberman, 1982)). The thickest canopy should be observed at the switch from leaf flushing to leaf fall, which occurs in mid-July. This pattern becomes more pronounced in drier sites compared to wet sites. In the driest plots (woody savanna) and during the driest month (February), a small portion of trees still retains leaves, as shown in our hemispherical photographs, whereas the wet evergreen plots remain fully green (Supplementary Figure 2). The seasonal pattern of LAI is clearly visible in hemispherical photos in the woody savanna (Supplementary Figure 13), but not visually obvious at other sites (Supplementary Figure 5).

***A2.Spatial variation in the study sites***

Given the differences in tree number and aboveground biomass between study sites, it is reasonable to assume that LAI in woody savanna plots would be lower than that in dry forest plots, which would in turn be lower than that in semi-deciduous plots. However, the difference between semi-deciduous plots and wet evergreen plots is difficult to discern through visual inspection of the colour photographs (Supplementary Figure 5), and these two plot types have very similar stem biomass (Zhang-Zheng et al., 2024).

***A3.Climate data and rainfall seasonality***

The Seasonality Index (SI) (Figure 1) was computed using the following formula:

SI = (1/R) Σ |Xj - R/12|

where Xj is monthly precipitation and R is total annual rainfall (Imteaz and Hossain, 2023).

We processed the CHIRPS daily precipitation dataset (2000–2020) using Google Earth Engine (Funk et al., 2015), where we aggregated rainfall data to monthly totals, computed annual rainfall sums for each pixel, and derived the SI values for each year. The final output represents the 21-year mean SI, which characterises long-term rainfall seasonality patterns across the study area.

***A4.Hemispherical photography data collection***

Twenty-five hemispherical photo sampling points were systematically distributed within each 1-ha plot following the GEM protocol (Malhi et al., 2021). This sampling approach is in line with common hemispherical photography sampling strategies used in forest canopies (Brown et al., 2023; Demarez et al., 2008; Weiss et al., 2004). Monthly photographs were collected from March 2012 to July 2019 at BOB, from June 2013 to July 2019 at KOG, and from January 2016 to July 2019 at ANK. All images were acquired using a Nikon Coolpix 4500 digital camera equipped with a Nikon Fisheye Converter FC-E8 0.21× lens. To aid interpretation of seasonal LAI variations, we present representative field photographs and their corresponding hemispherical images (Supplementary Figures 2–5).

***A5.Green Chromatic Coordinate***

To provide independent evidence of canopy phenology beyond hemispherical photos (bottom-up), we quantified leaf flushing phenology using Green Chromatic Coordinate (GCC) at BOB-02. Data were collected via digital repeat photography from a top-down camera mounted on a 60 m tower overlooking the canopy (Supplementary Figure 12). Images were captured every 10 minutes, 24 hours a day, from November 2022 to November 2023. To maintain consistent lighting and minimise shadow effects, a single daily image was selected as close to solar noon as possible. Images obscured by nightfall, poor weather, fog, or low illumination were manually excluded.

Image analysis and vegetation index extraction were performed using the Phenopix package in R. We focused on dominant species at the study plot, including Triplochiton scleroxylon, Terminalia superba, Pterygota macrocarpa, Sterculia rhinopetala, Alstonia boonei, and Guarea cedrata. The GCC shown in the study is the maximum GCC among these species. For each tree crown, we calculated GCC as: GCC = G / (R + G + B), where R, G, and B represent the red, green, and blue channels. The resulting time series were filtered to reduce noise using a sequential three-step protocol: a night filter to remove low-illumination data, a spline filter for outlier removal, and a max filter to identify values in the 90th percentile within a three-day moving window.

Brown, L.A., Morris, H., Leblanc, S., Bai, G., Lanconelli, C., Gobron, N., Meier, C., Dash, J., 2023. HemiPy: A python module for automated estimation of forest biophysical variables and uncertainties from digital hemispherical photographs. Methods Ecol. Evol. 14, 2329–2340. <https://doi.org/10.1111/2041-210X.14199>

Demarez, V., Duthoit, S., Baret, F., Weiss, M., Dedieu, G., 2008. Estimation of leaf area and clumping indexes of crops with hemispherical photographs. Agric. For. Meteorol. 148, 644–655. <https://doi.org/10.1016/j.agrformet.2007.11.015>

Funk, C., Peterson, P., Landsfeld, M., Pedreros, D., Verdin, J., Shukla, S., Husak, G., Rowland, J., Harrison, L., Hoell, A., Michaelsen, J., 2015. The climate hazards infrared precipitation with stations—a new environmental record for monitoring extremes. Sci. Data 2, 150066. <https://doi.org/10.1038/sdata.2015.66>

Imteaz, M.A., Hossain, I., 2023. Climate change impacts on ‘seasonality index’ and its potential implications on rainwater savings. Water Resour. Manage. 37, 2593–2606. <https://doi.org/10.1007/s11269-022-03320-z>

Lieberman, D., 1982. Seasonality and phenology in a dry tropical forest in ghana. The Journal of Ecology 70, 791. <https://doi.org/10.2307/2260105>

Malhi, Y., Girardin, C., Metcalfe, D.B., Doughty, C.E., Aragão, L.E.O.C., Rifai, S.W., Oliveras, I., Shenkin, A., Aguirre-Gutiérrez, J., Dahlsjö, C.A.L., Riutta, T., Berenguer, E., Moore, S., Huasco, W.H., Salinas, N., Da Costa, A.C.L., Bentley, L.P., Adu-Bredu, S., Marthews, T.R., Meir, P., Phillips, O.L., 2021. The global ecosystems monitoring network: Monitoring ecosystem productivity and carbon cycling across the tropics. Biol. Conserv. 253, 108889. <https://doi.org/10.1016/j.biocon.2020.108889>

Weiss, M., Baret, F., Smith, G.J., Jonckheere, I., Coppin, P., 2004. Review of methods for in situ leaf area index (LAI) determination: Part II. Estimation of LAI, errors and sampling. Agric. For. Meteorol. 121, 37–53. <https://doi.org/10.1016/j.agrformet.2003.08.001>

Zhang-Zheng, H., Adu-Bredu, S., Duah-Gyamfi, A., Moore, S., Addo-Danso, S.D., Amissah, L., Valentini, R., Djagbletey, G., Anim-Adjei, K., Quansah, J., Sarpong, B., Owusu-Afriyie, K., Gvozdevaite, A., Tang, M., Ruiz-Jaen, M.C., Ibrahim, F., Girardin, C.A.J., Rifai, S., Dahlsjö, C.A.L., Riutta, T., Deng, X., Sun, Y., Prentice, I.C., Oliveras Menor, I., Malhi, Y., 2024. Contrasting carbon cycle along tropical forest aridity gradients in west africa and amazonia. Nat. Commun. 15, 3158. <https://doi.org/10.1038/s41467-024-47202-x>

### Appendix B. Supplementary Figures

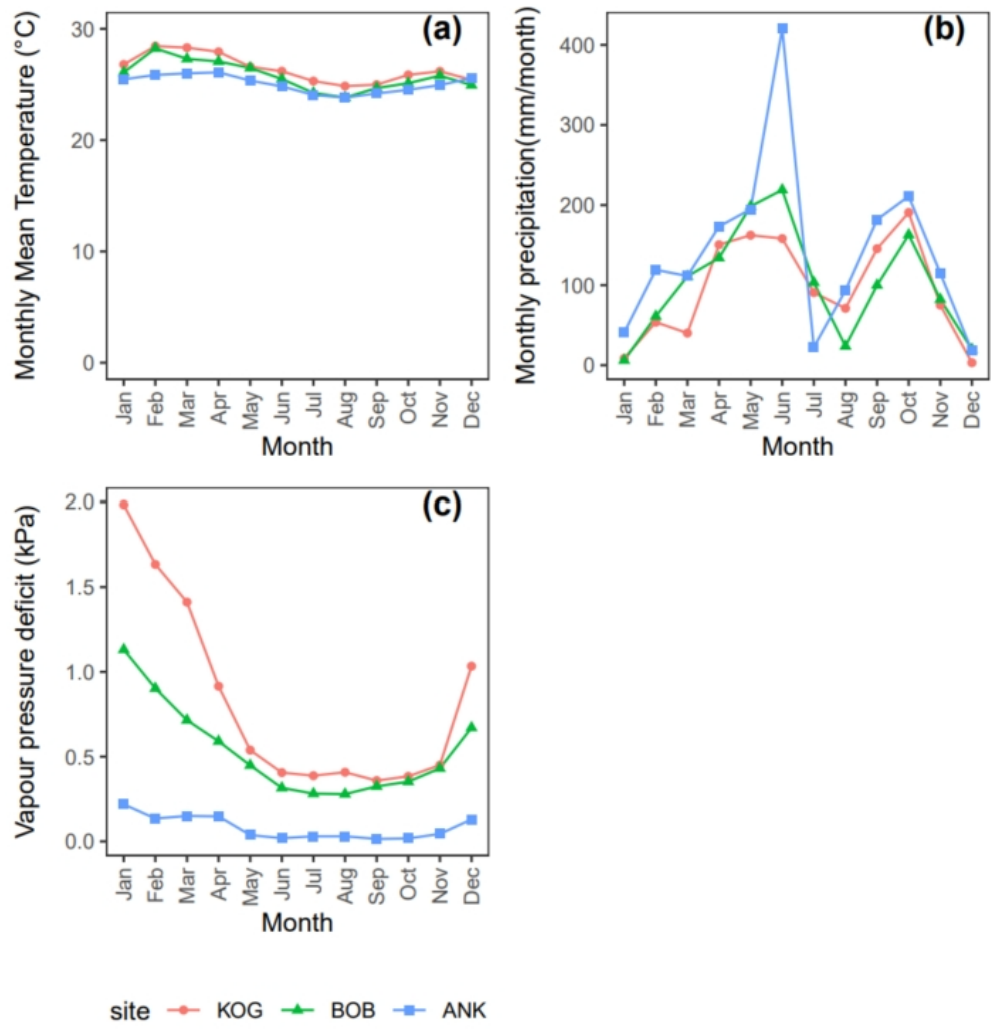

Supplementary Figure 1 Study site seasonality. We show (a) Monthly mean temperature (oC) (b) monthly sum precipitation (mm/month) and (c) vapour pressure deficit (kPa) for the three study sites. Monthly mean LAI was a mean across 2016-2018(for ANK) and 2012-2019 (for BOB&KOG). Climate data were reported by local meteological stations across 2011(for ANK), 2012-2016 (for BOB), and 2013-2016(for KOG),as described in Zhang-Zheng et al. (2025).

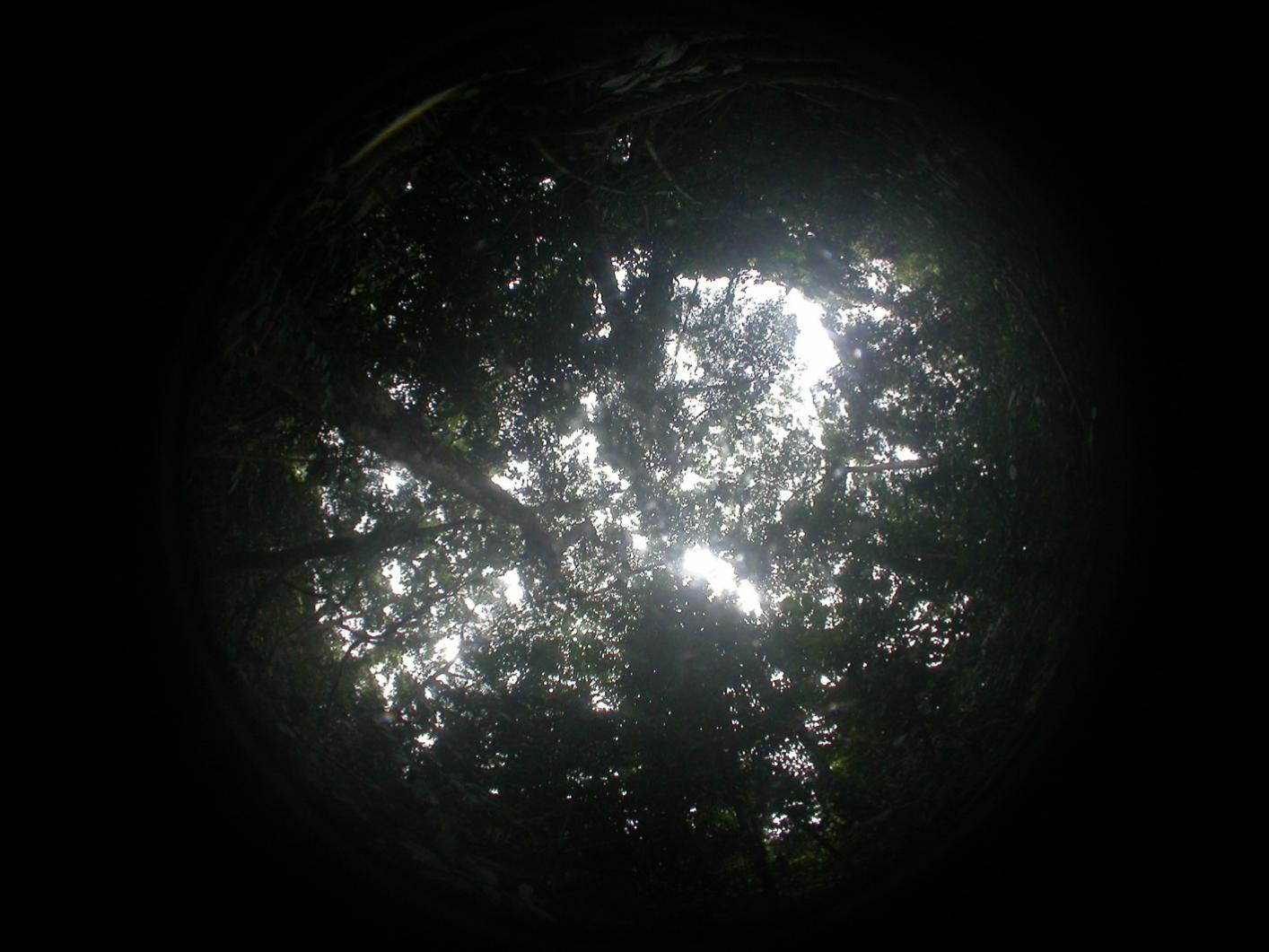

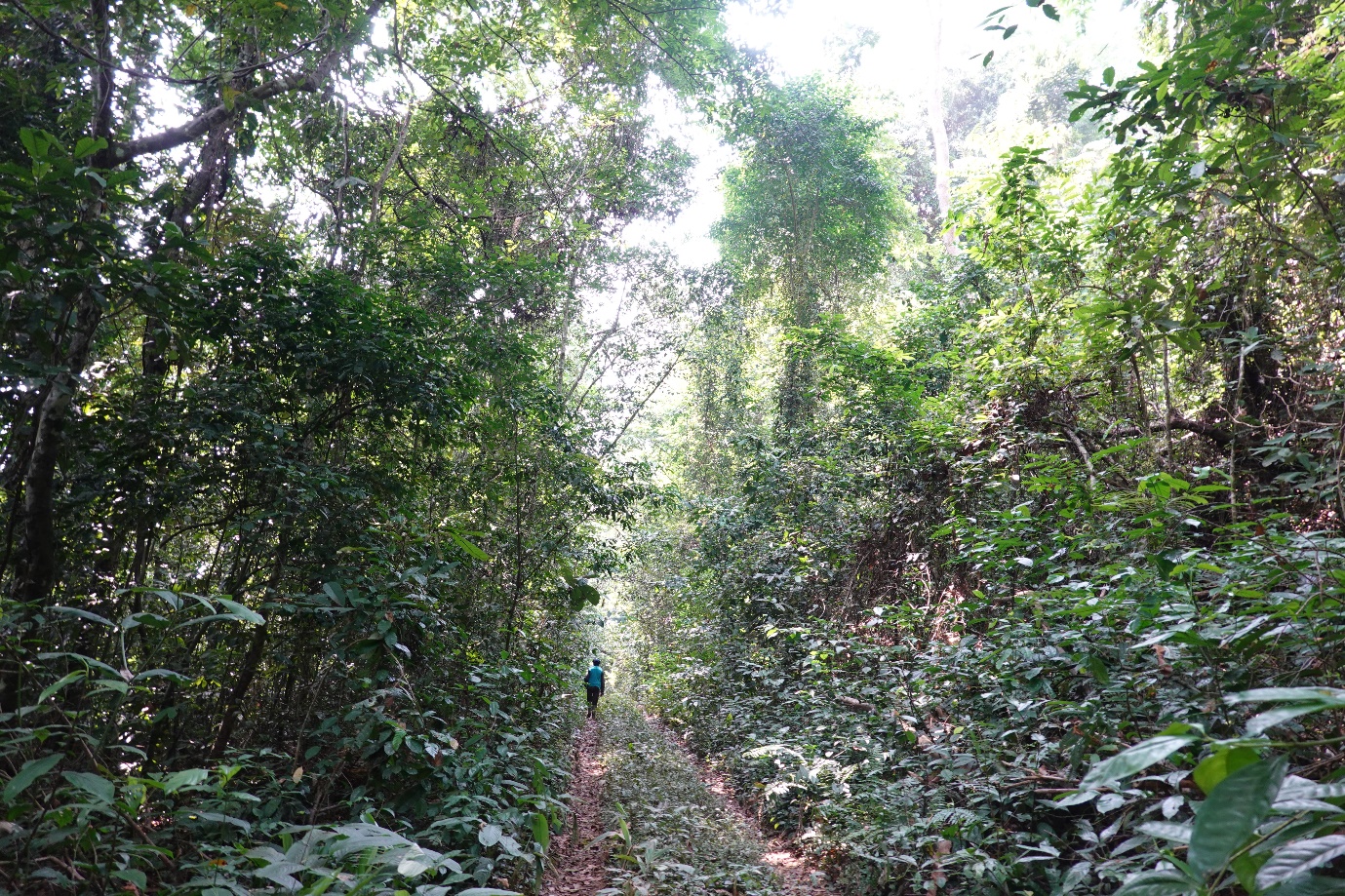

(a)

(b)

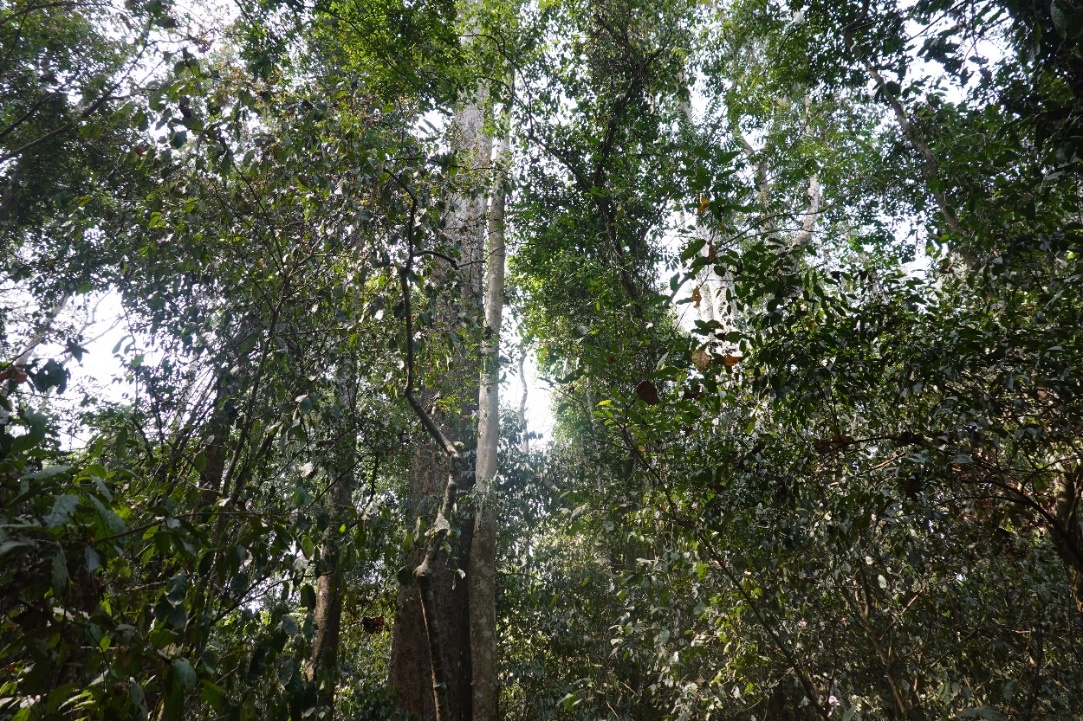

(c)

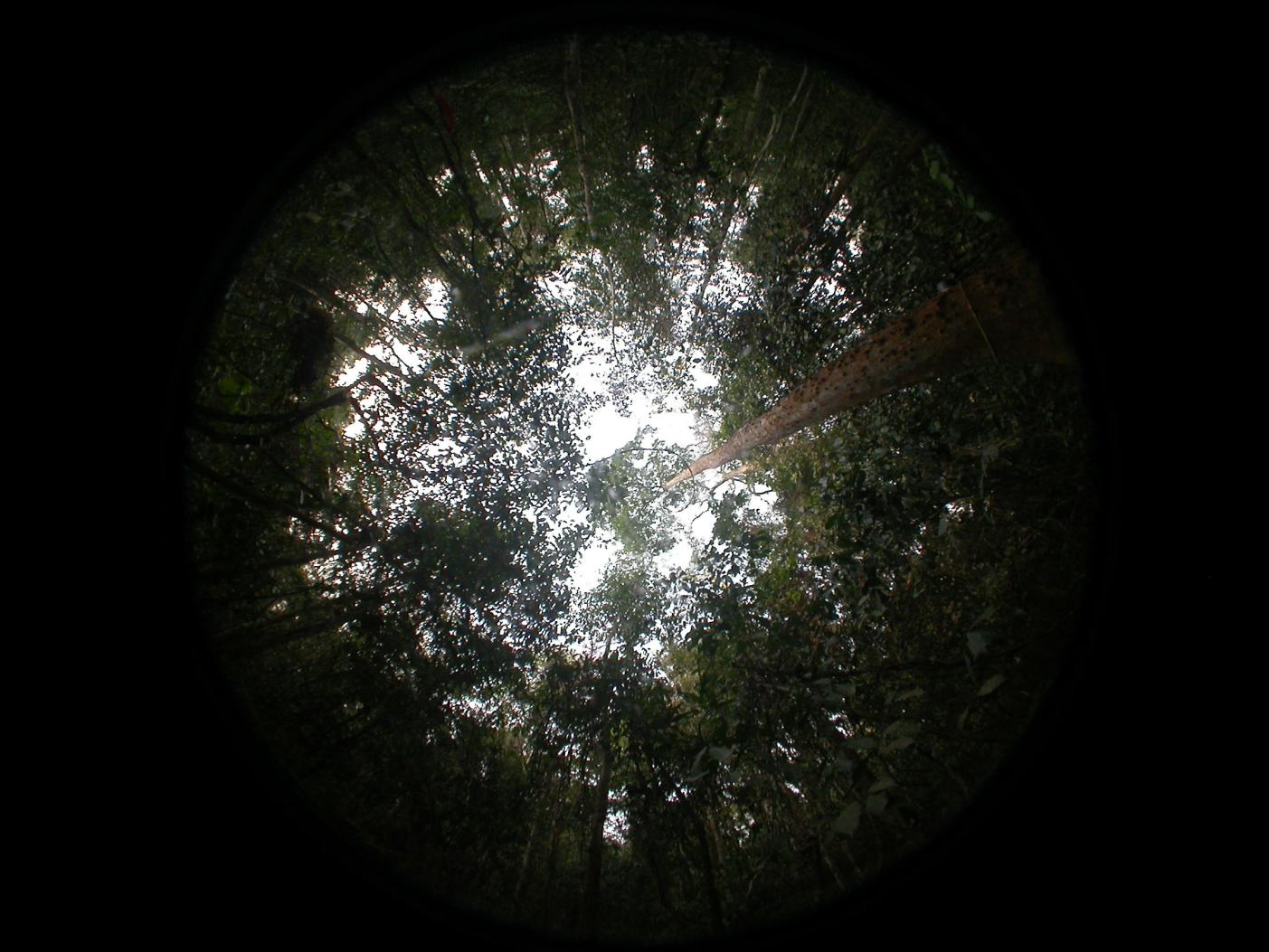

(d)

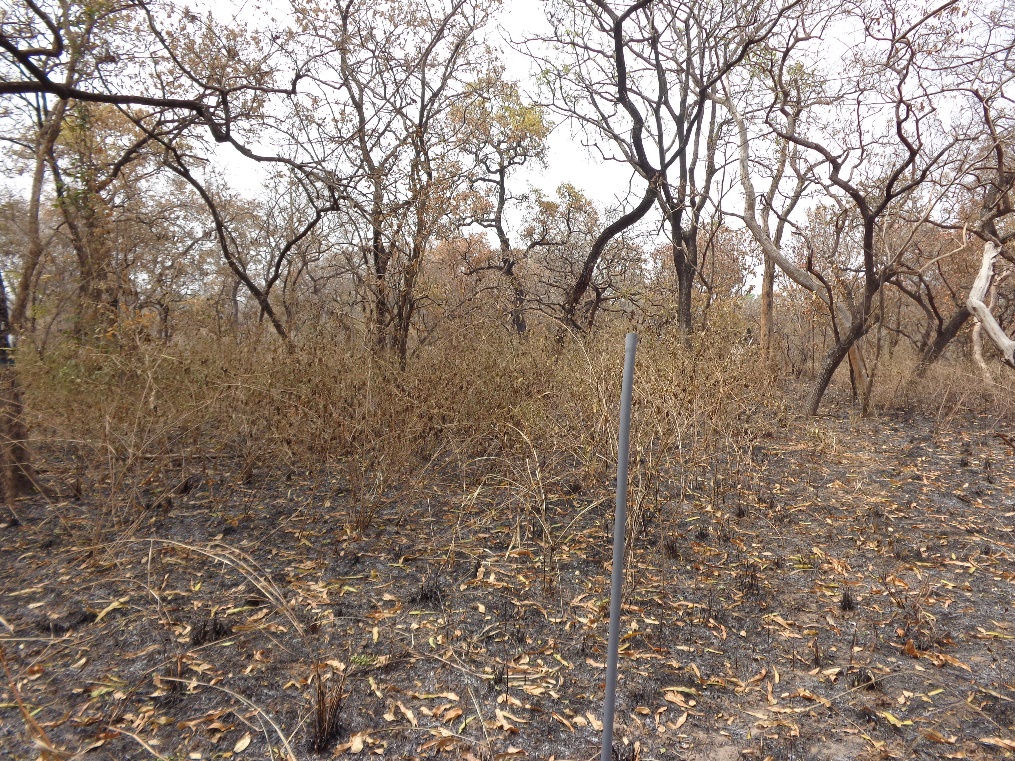

(e)

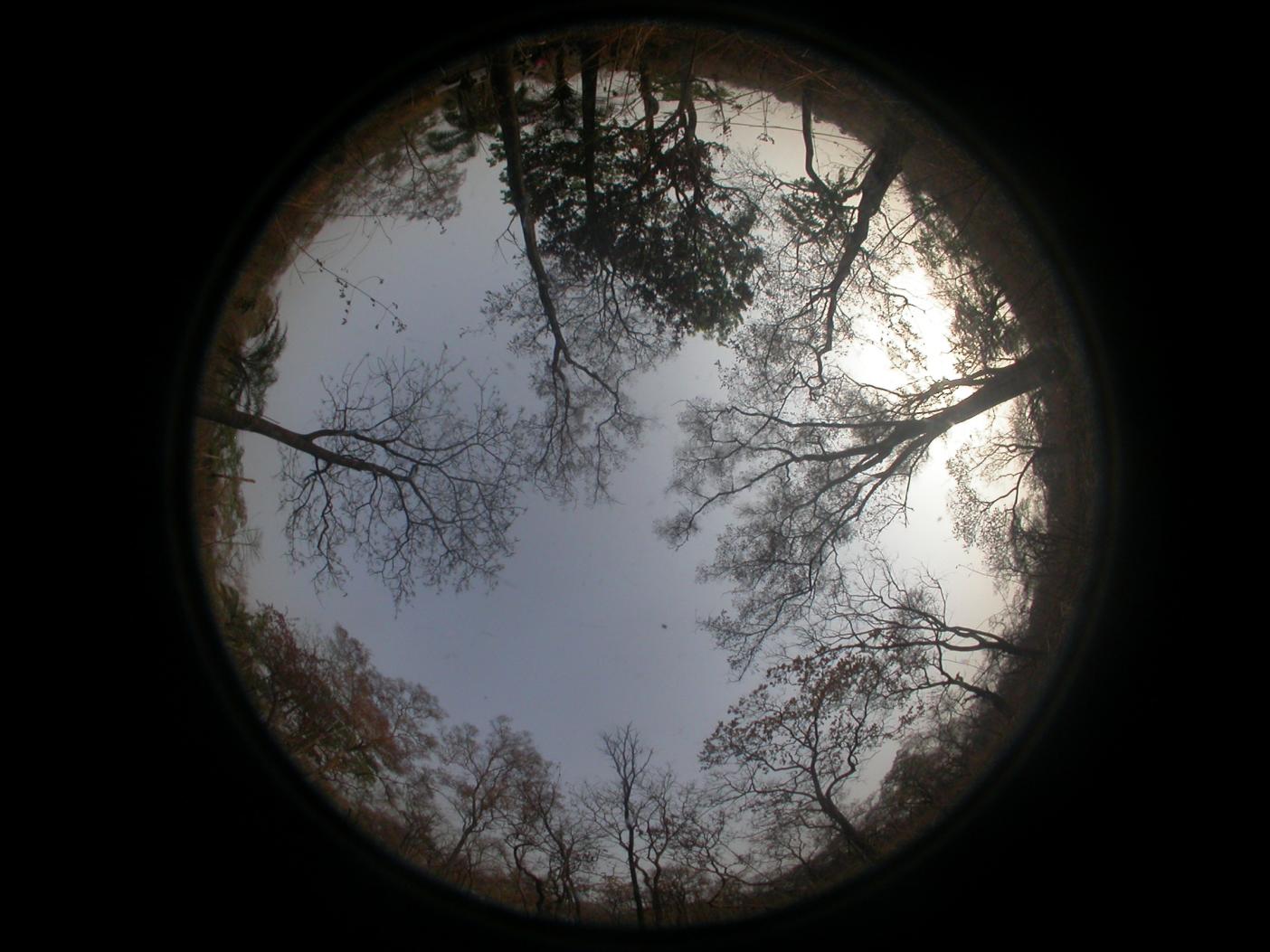

(f)

Supplementary Figure 2 Photos of the study sites and examples of their hemispherical photos. Here we show (a) the path leading to site ANK, at the entrance of ANK; At the site, the canopy often goes white in a normally exposed photo. We decrease exposure for most hemispherical photos but they still appear over-exposed. (b) hemispherical photo of plot ANK01 in February 2018 (dry season); This photo looks over-exposed, but lowering exposure will make it very dark and erase sky gap at the horizon. We could not know the weather above the canopy in the field because the forests are super dense. Therefore, photos at this plot are taken under varying weather condition(c) study plot BOB-01 (d) hemispherical photo of BOB-02 in February 2018; (e) Study plot KOG-05 after fire (f) hemispherical photo of KOG-05 taken in February 2018. The lighting condition is very different from (d).

ANK-01	2019.2

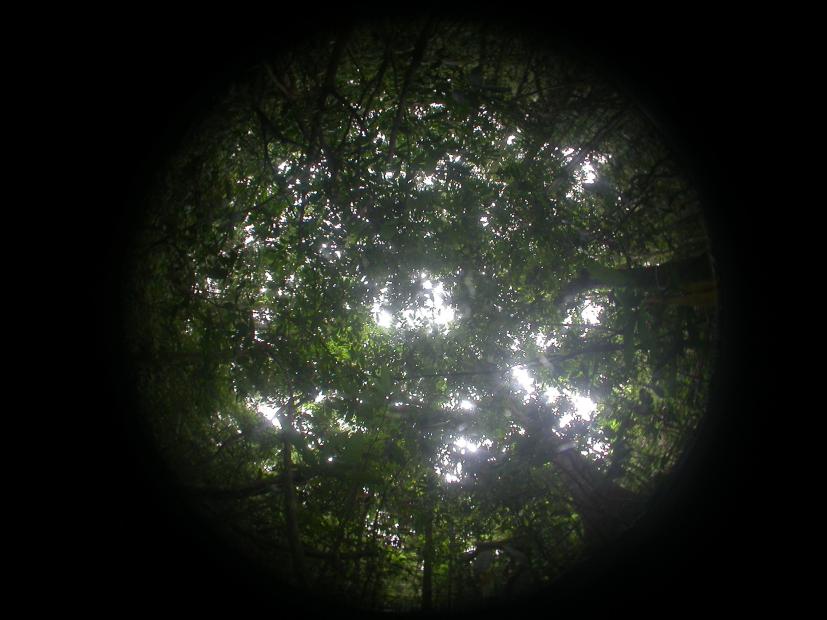

ANK-01	2019.4

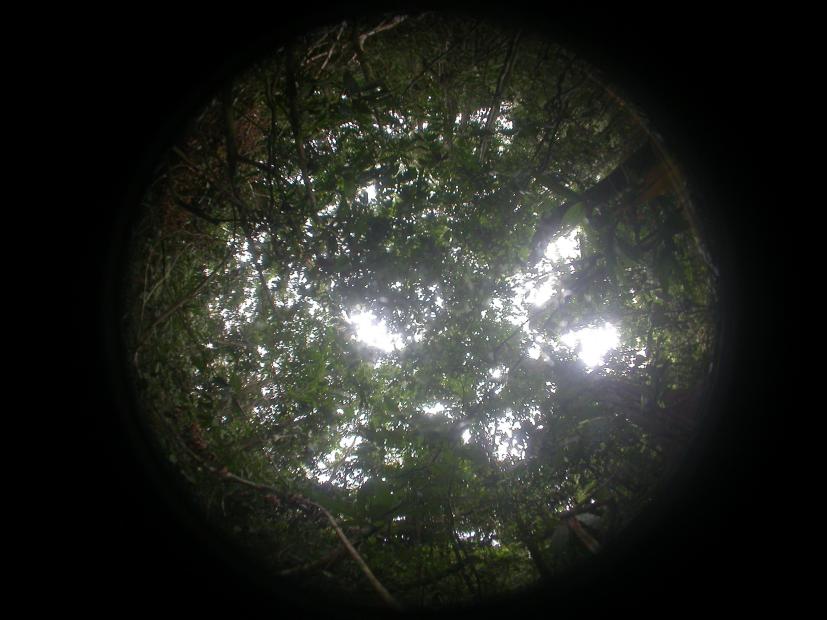

ANK-01	2019.2

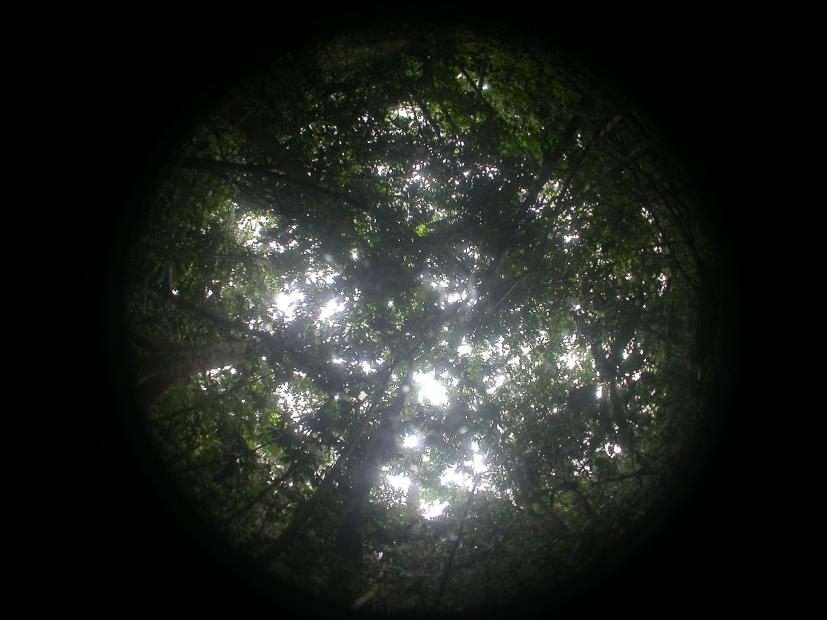

ANK-01	2019.4

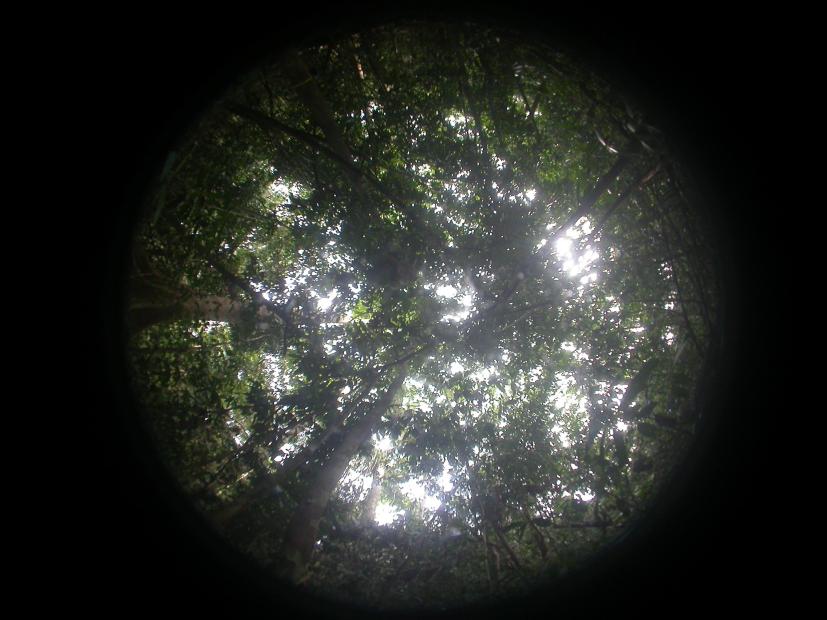

Supplementary Figure 3 Fish-eye photos (ANK-01) taken in February and April 2019. Each pair of photos (on the same page) is taken at the same photo point. By comparing with the naked eye, we can see that in the fish-eye photo from April 2019, there are fewer leaves because its sky window is larger and the edges are more fragmented. The canopy of this plot is extremely dense, and in the dry season (February), the contrast is extremely high, so most photos appear over-exposed even when we decrease exposure.

KOG-05 July 2017

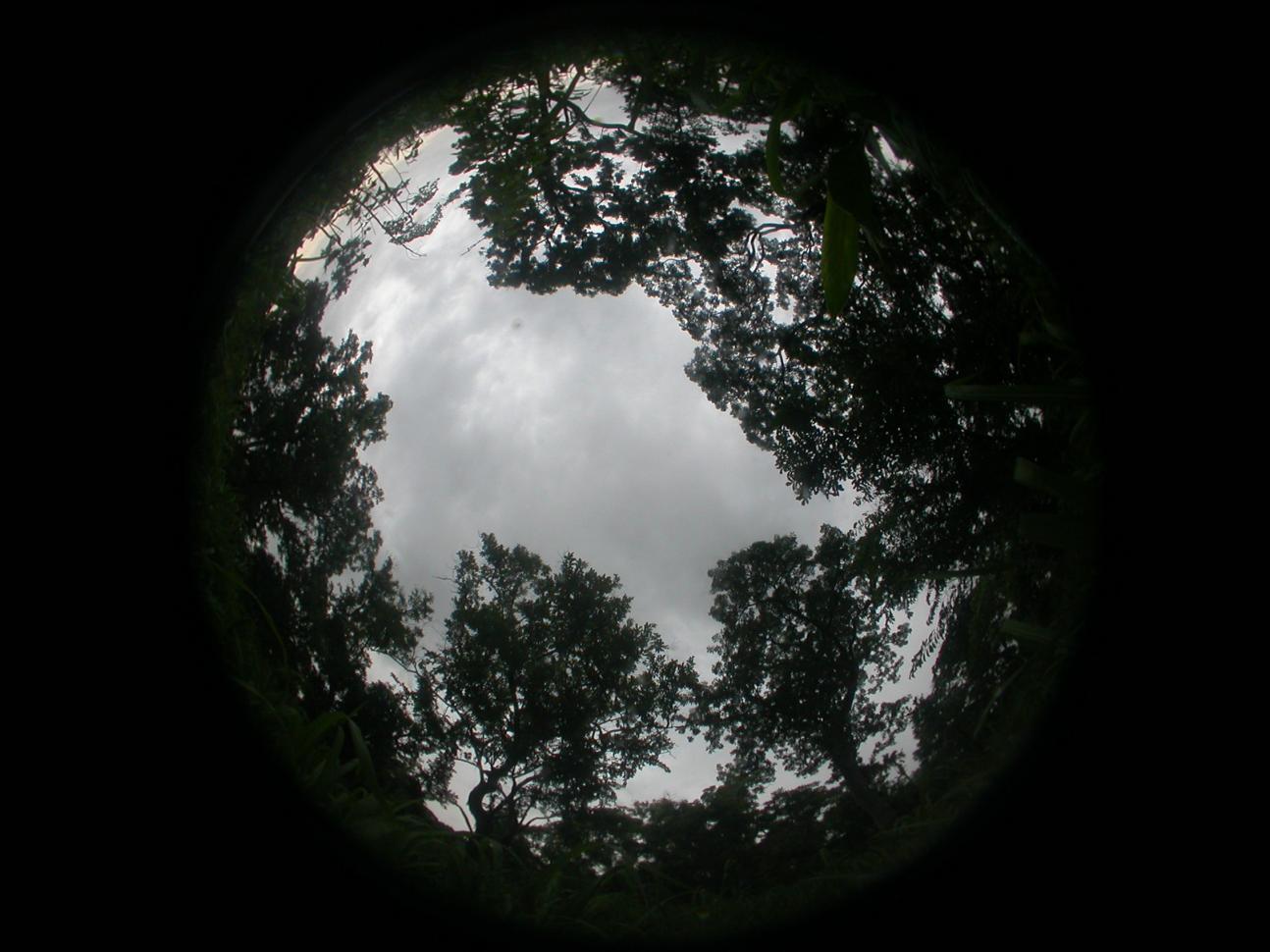

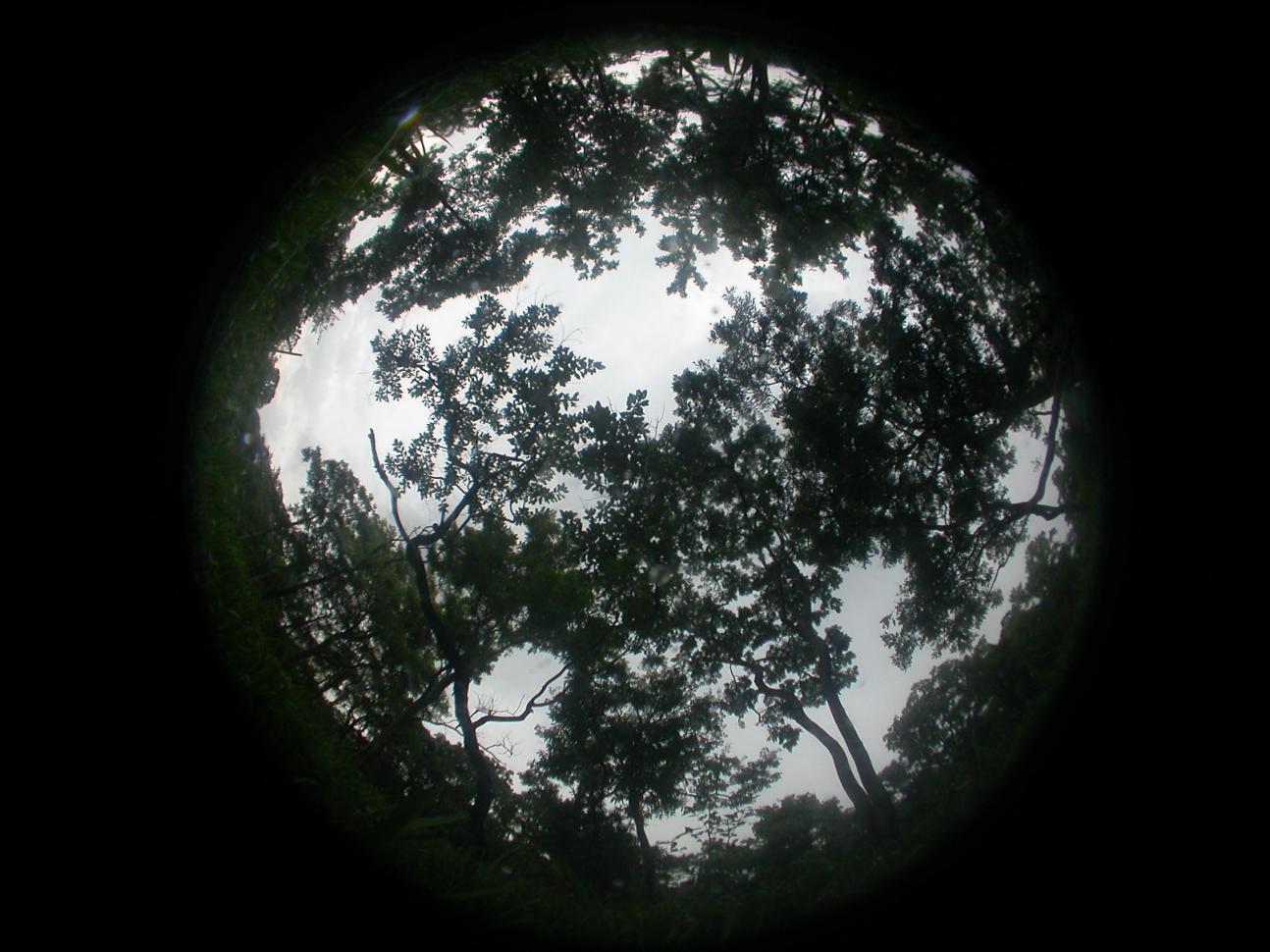

KOG-03 July 2017

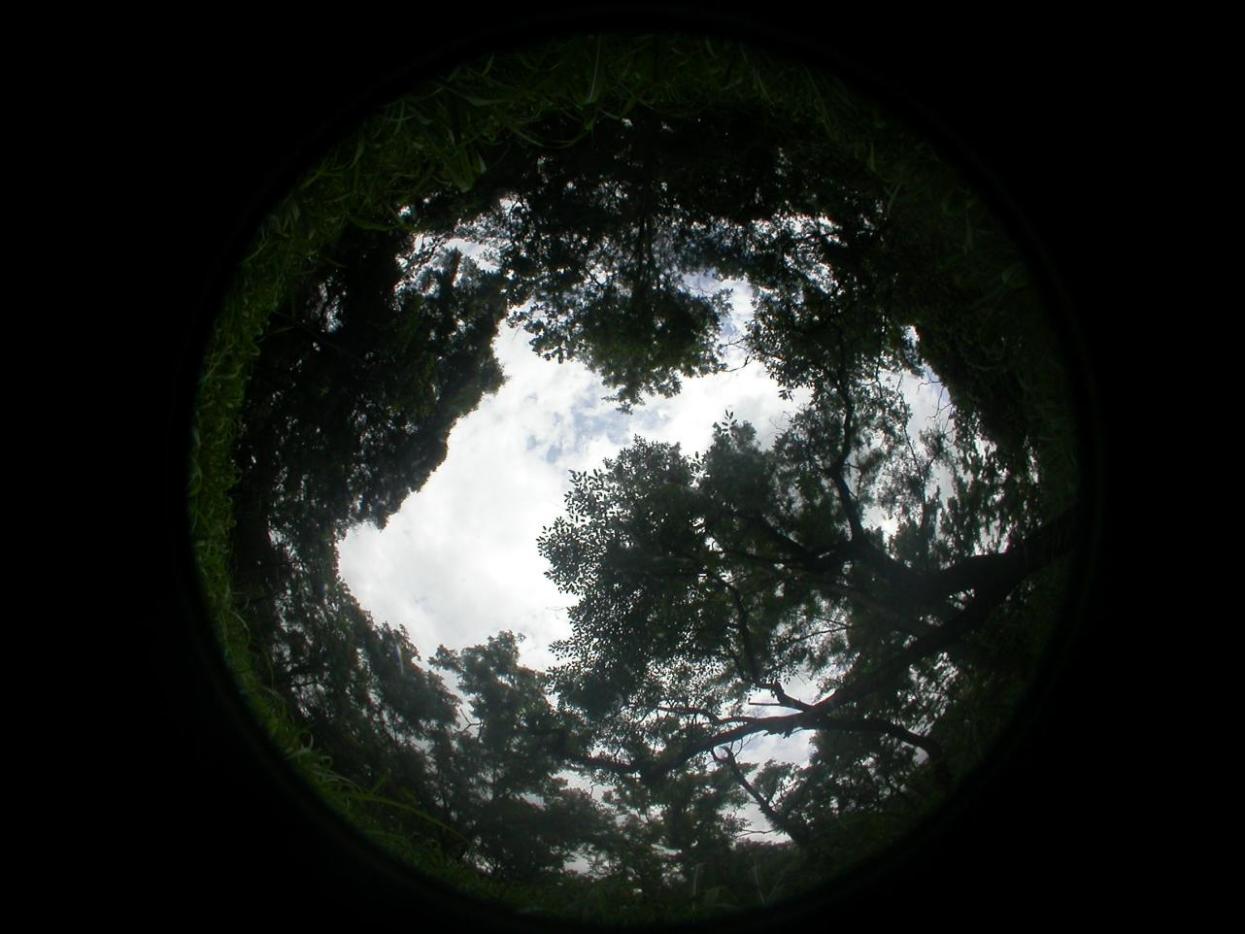

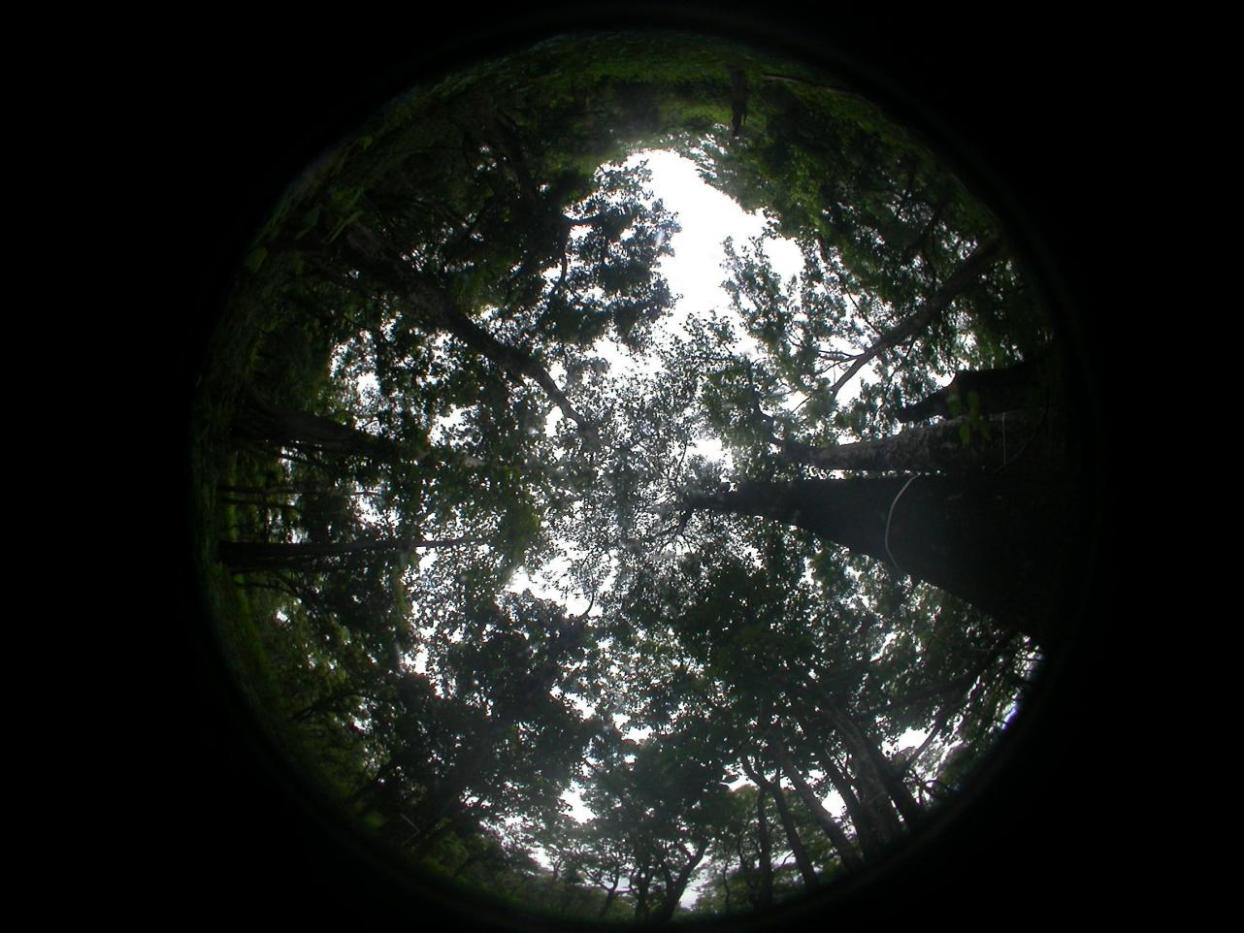

Supplementary Figure 4 Contrasting photos at the same forest plot showing the ‘large-sky’ problem (a big patch of sky that is not homogeneous). We show KOG-03 and KOG-05 in July. If the lower photo is well marked through CAN-EYE, the upper photo in the same batch of photos will put a large area of the sky into the leaves. On the contrary, if the upper image is well labelled, many leaves on the lower will be classified as the sky.

|  | ANK-03 | BOB-04 |
| --- | --- | --- |
| 2018-1 | 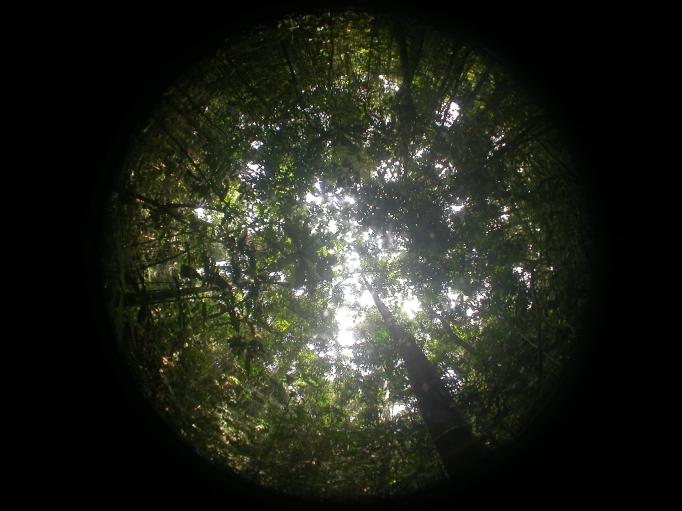 | 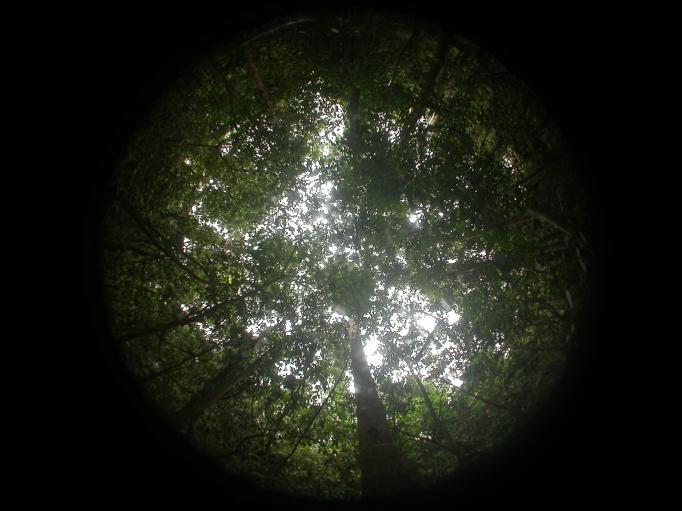 |
|  | 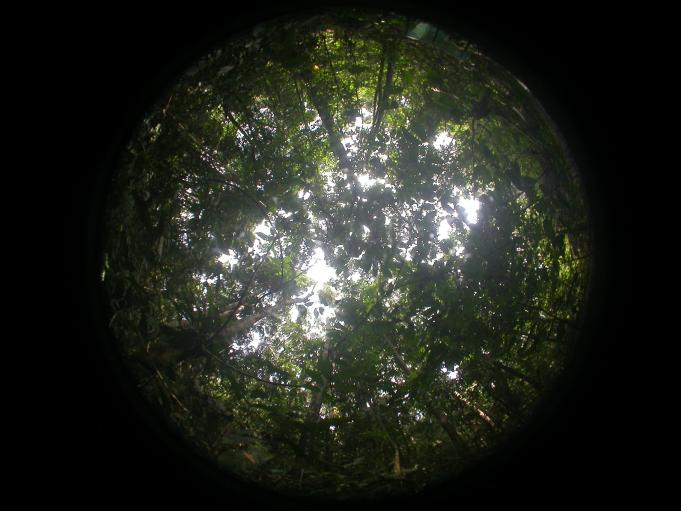 | 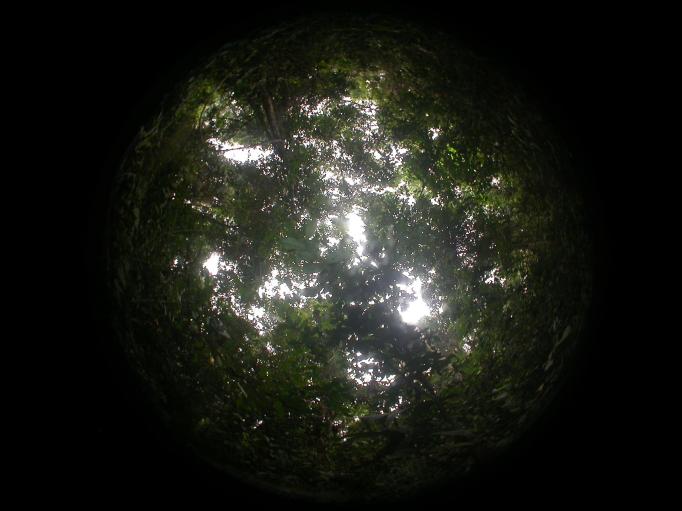 |
| 2018-3 | 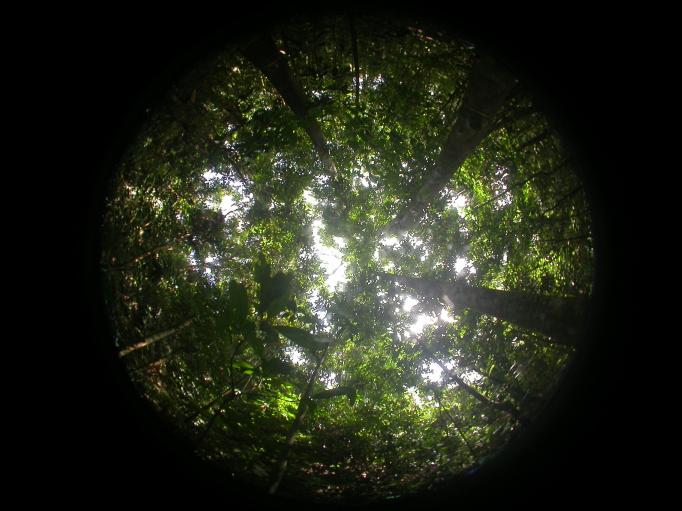 | 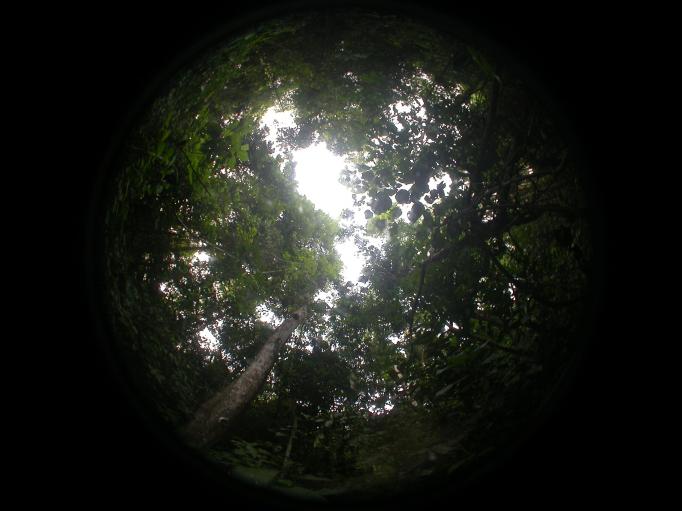 |
|  | 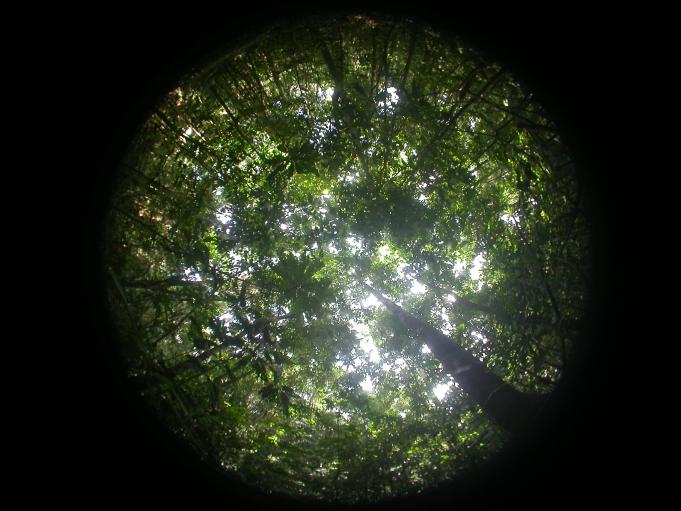 | 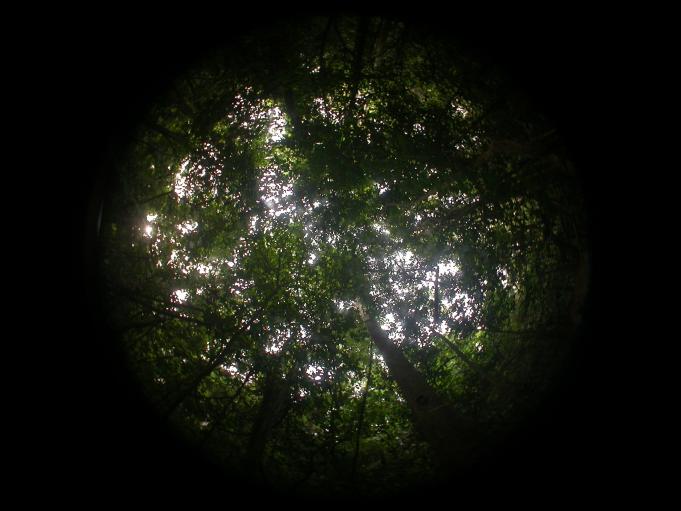 |
| 2018-5 | 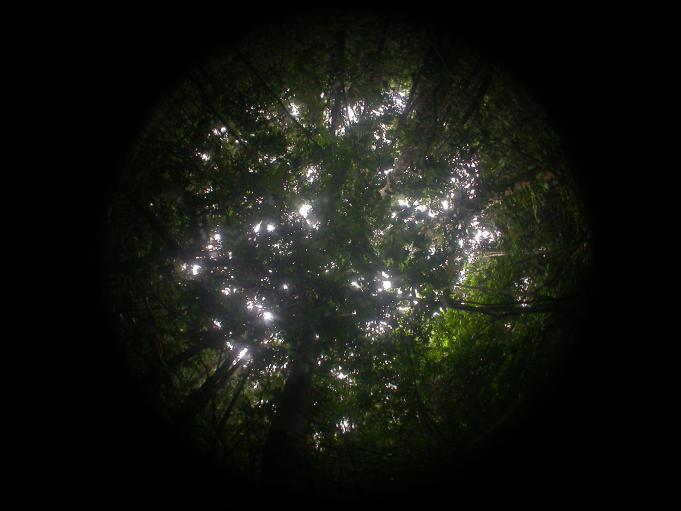 | 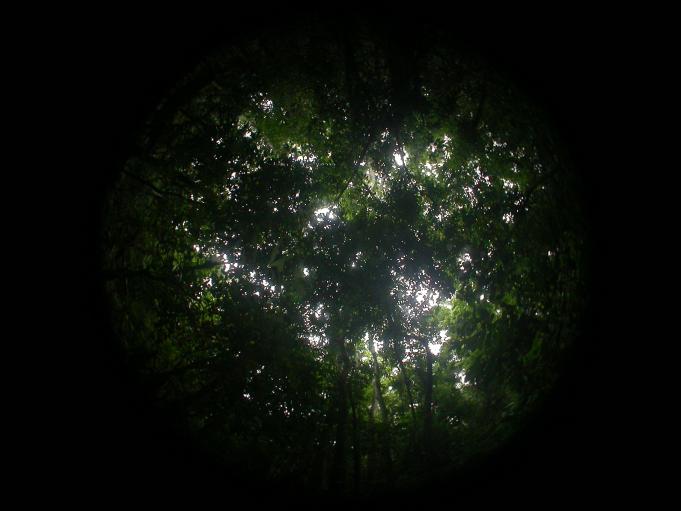 |
|  | 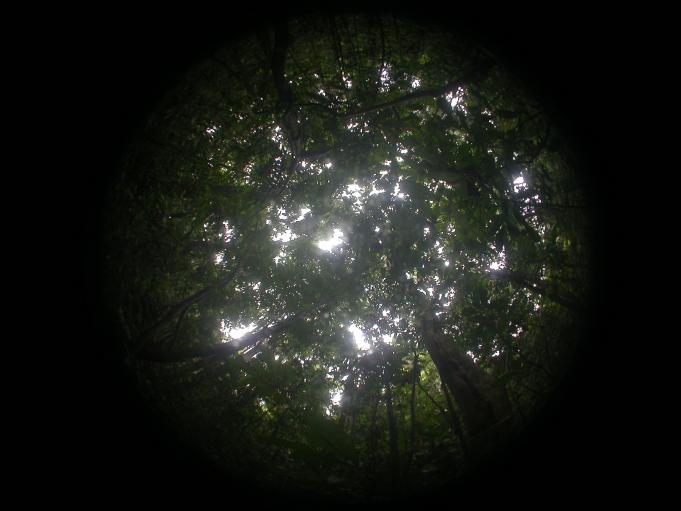 | 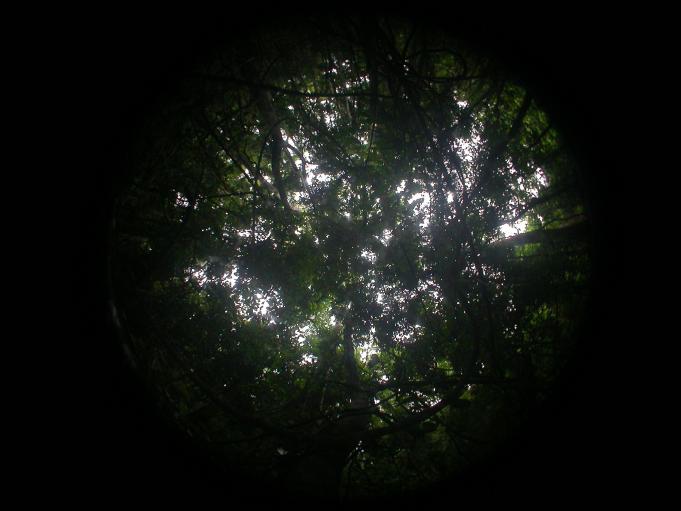 |

Supplementary Figure 5 Fisheye photos of ANK-03 (left) and BOB-04 (right). Numbers on the left are year-month. By visually comparing these fish-eye photos, it is rather difficult to say which study plot has more leaves.

Intensive sky beam

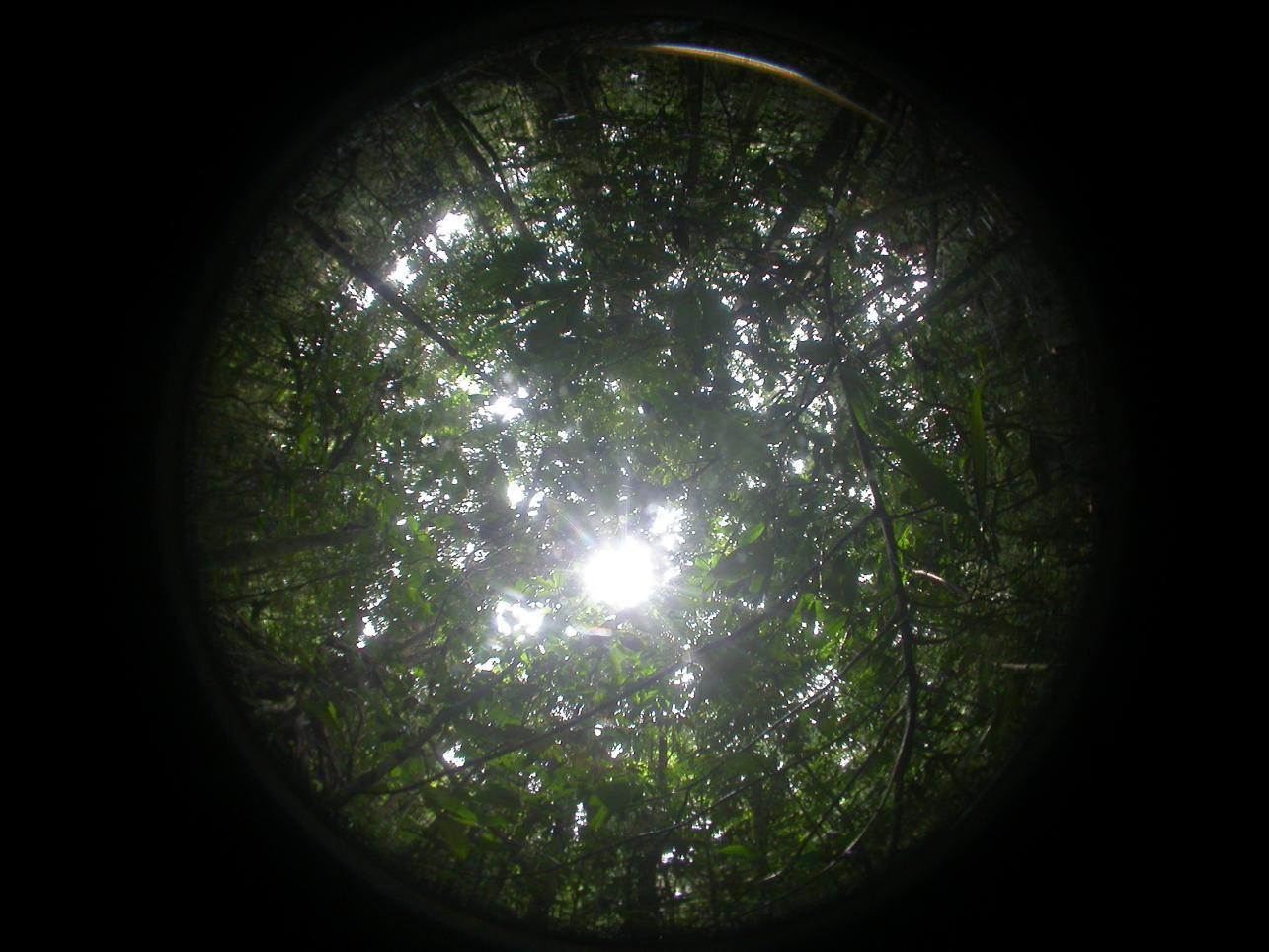

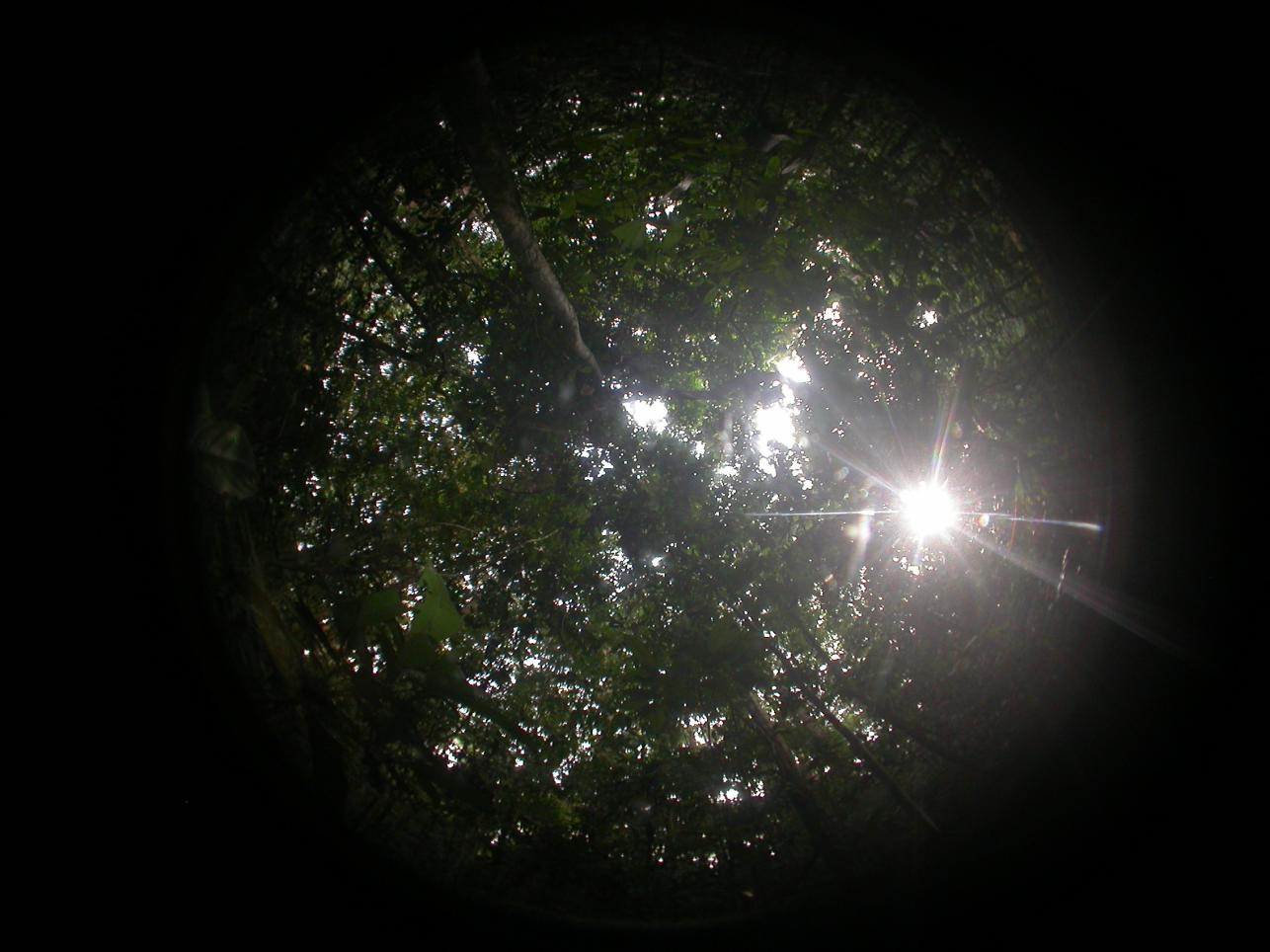

Small and dense leaf

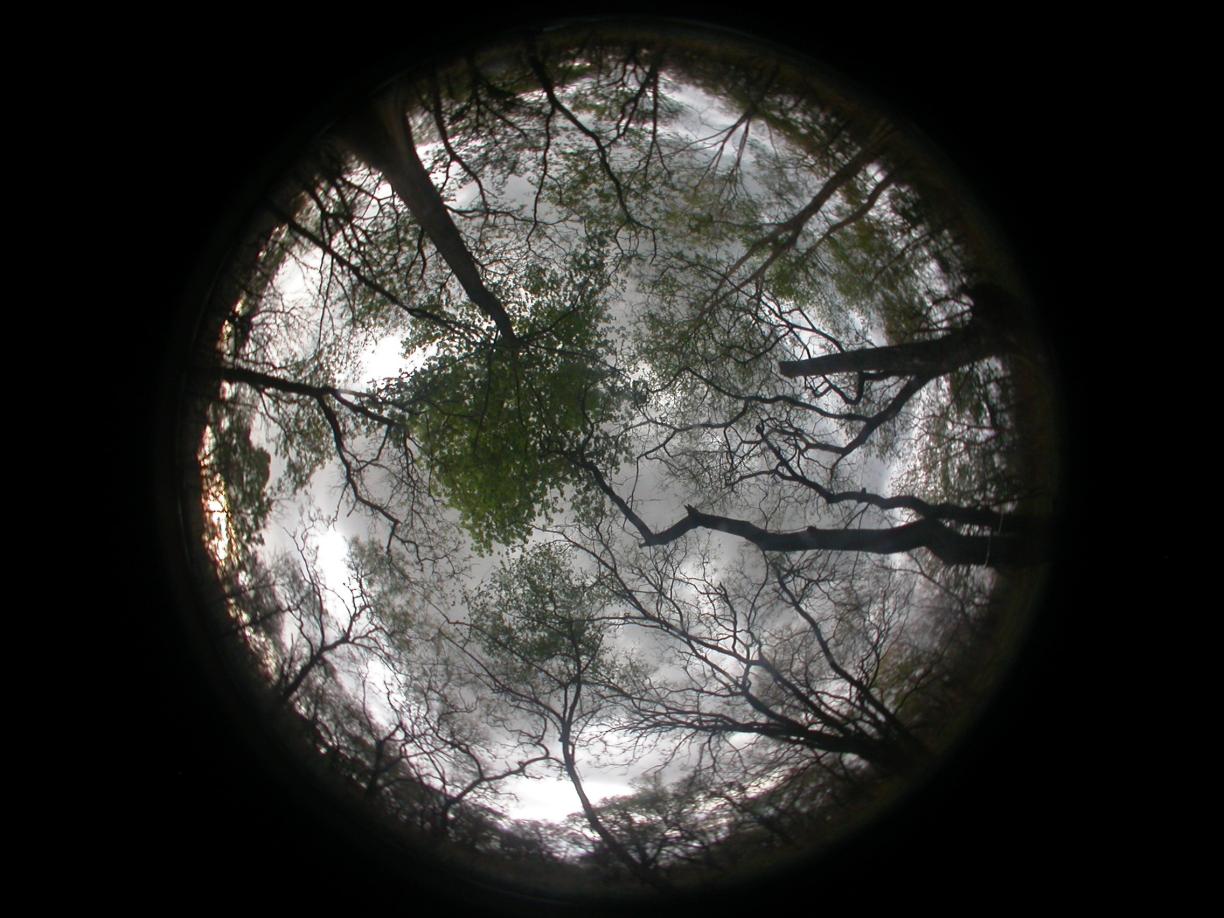

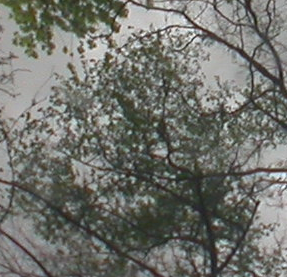

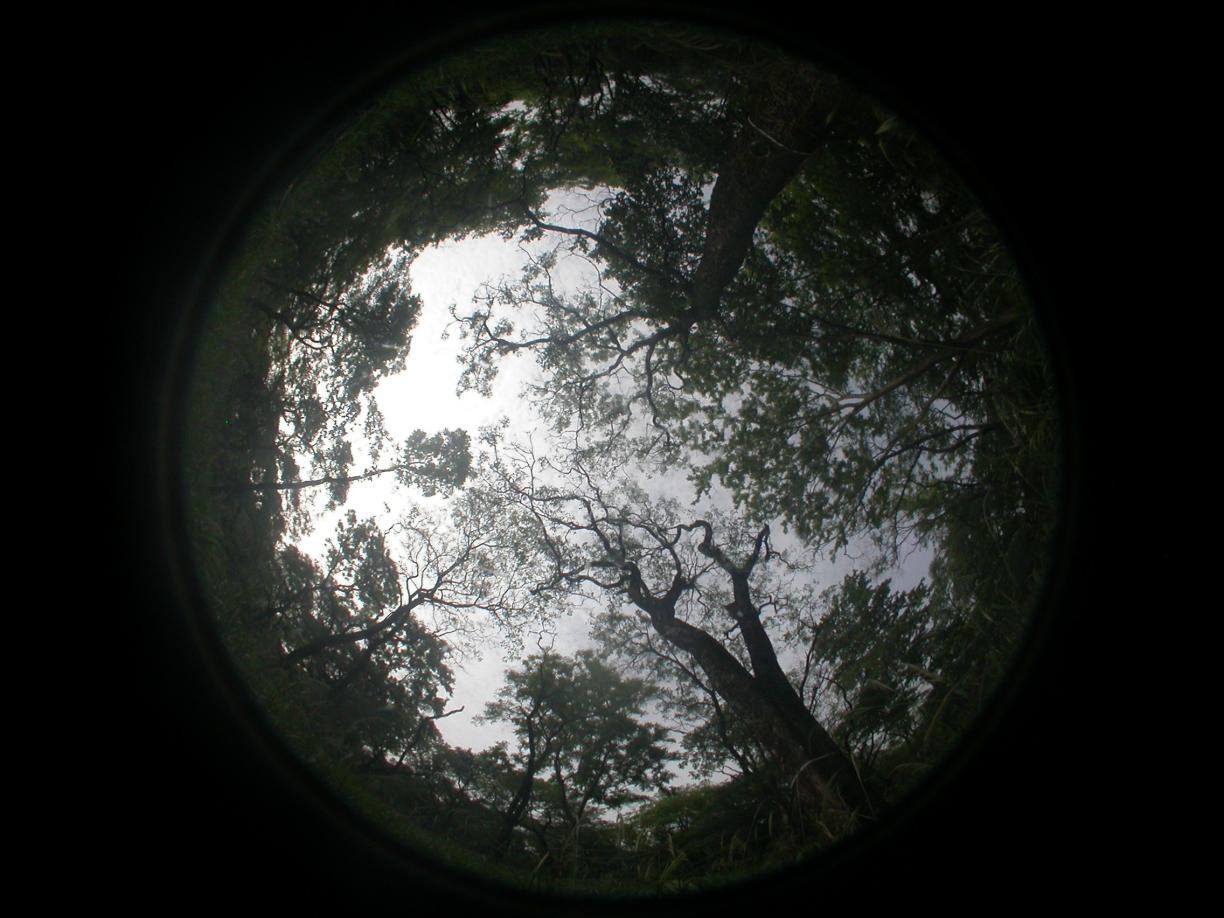

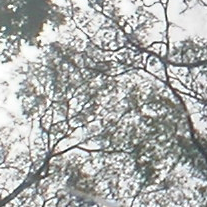

Small and dense twigs

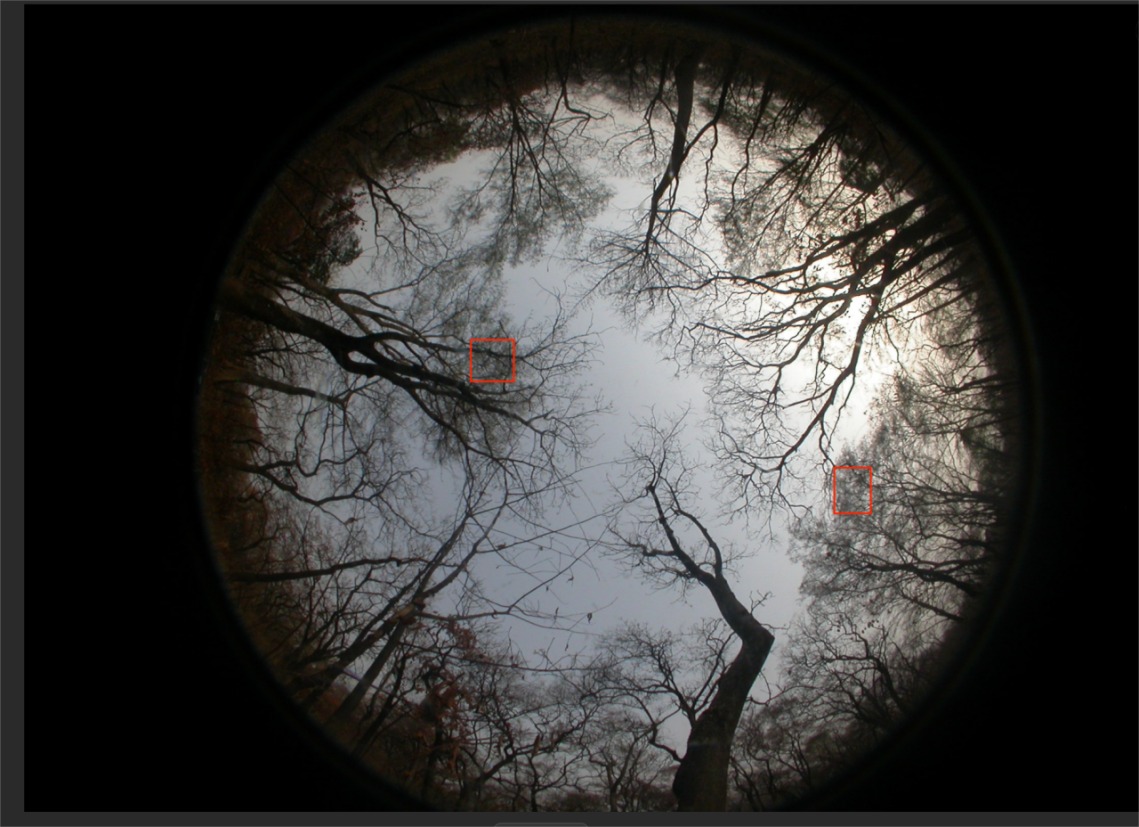

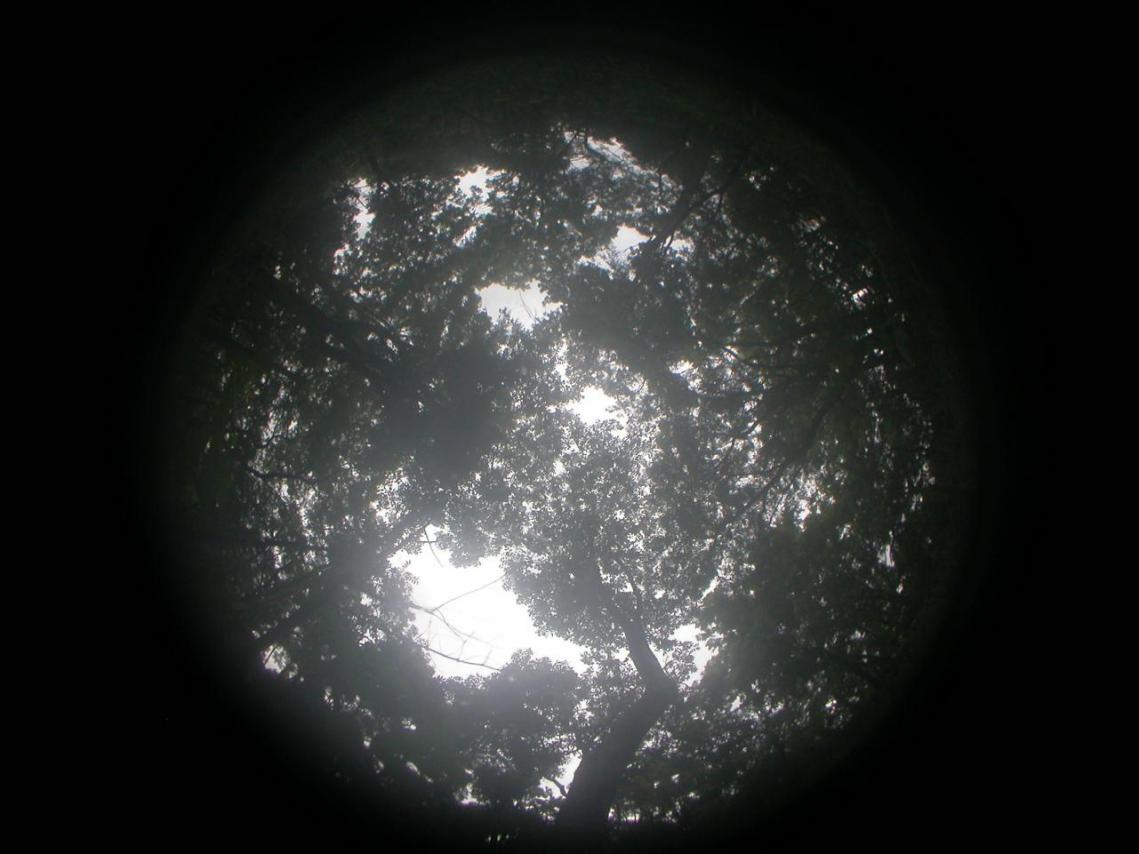

Supplementary Figure 6 The 'grey zone ' on hemispherical photos. Here are some images of ‘grey areas’, here we show two examples: (1) intensive sky beam, sunlight erosion on the leaves, making it difficult for humans to judge. It is not advisable to take photos in such sunny conditions. This is an extreme illustration – please do not take photos like this. However, a small sky beam could appear from time to time for tropical fieldwork because researchers often need to drive two hours to reach the study plot, and it is hard to anticipate the weather. The sky could appear very bright even in the rain because the canopy is super dense. (2) Small and dense leaf distribution, manual classification is hard to operate, the border between leaves and sky is fused and vague. (3) small dense twigs. The one above is a hemispherical canopy photograph from plot KOG06 in winter (February). The area marked with a red rectangle was double-checked by field staff, who confirmed that no leaves were present in this region at the time of imaging. The one below is at the same sampling point, but in summer. Overall, it is challenging for humans to make decisions when processing these areas. Subjective errors are particularly large when dealing with these photos.

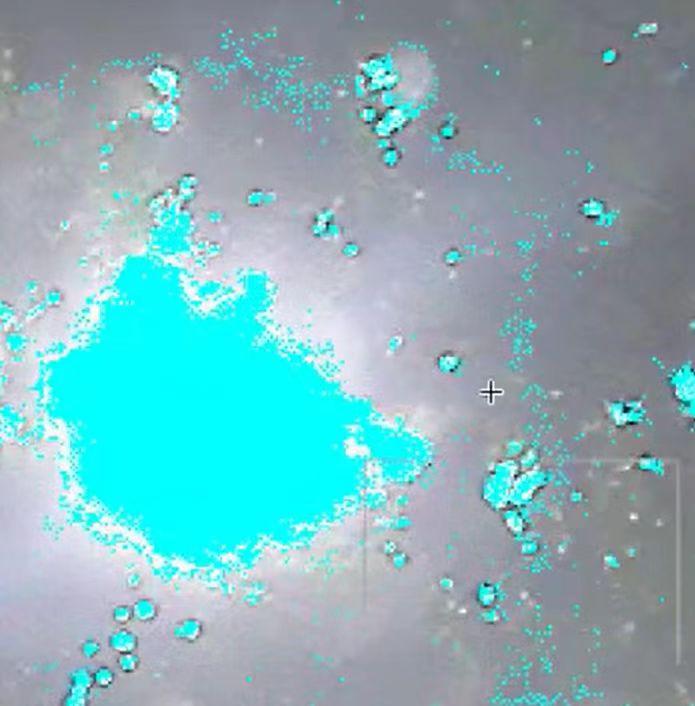

Supplementary Figure 7 Error pixel clusters in CAN-EYE processing at KOG06. In the edge regions of the sky (especially when sunlight is intense), CAN-EYE often produces dense pixel points, with gaps between them. These gaps are classified as part of the leaves, but in reality, they should be considered part of the sky. This situation is difficult to avoid, even with very careful processing. In this screenshot, blue is the classification outcome of sky, and the background is a zoom-in of a hemispherical photo.

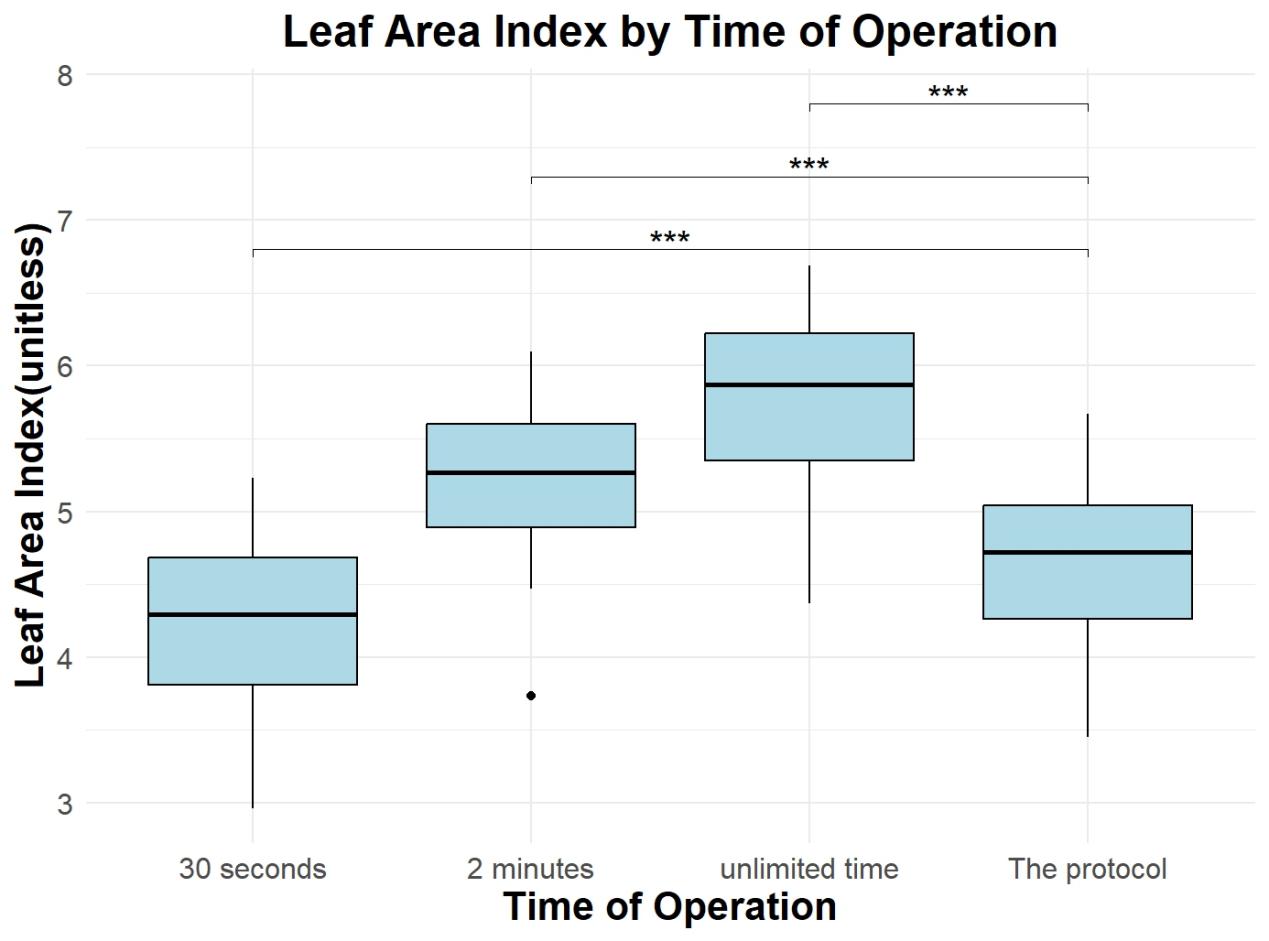

Supplementary Figure 8 Comparison of leaf area index (LAI) estimates under four different image processing methods: the protocol, 30 seconds, 2 minutes, and unlimited time. Boxplots show the distribution of LAI values obtained from two forest plots (ANK-01 and BOB-01) for each operation methods. A linear mixed-effects model was used to compare groups with the protocol as the reference and plot identity as a random effect. Asterisks (***) indicate statistically significant differences from the protocol (p < 0.001 for all comparisons).

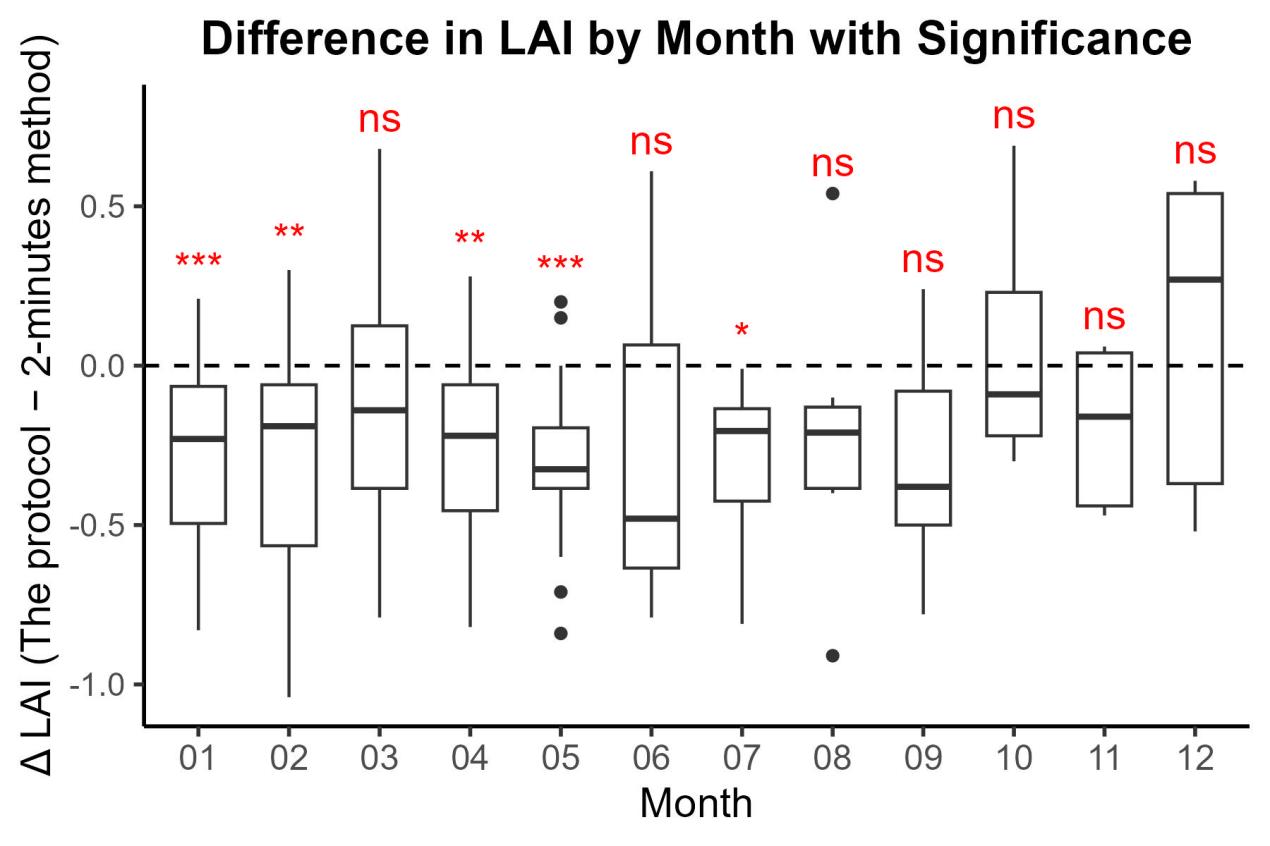

Supplementary Figure 9 Boxplots of **Δ LAI (The protocol minus 2-minutes method)** by calendar month (1 = January…12 = December). Each box depicts the distribution of LAI differences pooled across all plots in a given month. Positive Δ LAI values indicate that the protocol workflow yielded higher LAI estimates than the 2-minutes manual method. Significance of the mean Δ LAI versus zero was assessed by one-sample t-tests, with symbols denoting *p* < 0.05 (*), p < 0.01 (****), and p < 0.001 (***); “ns” indicates non-significance.

a

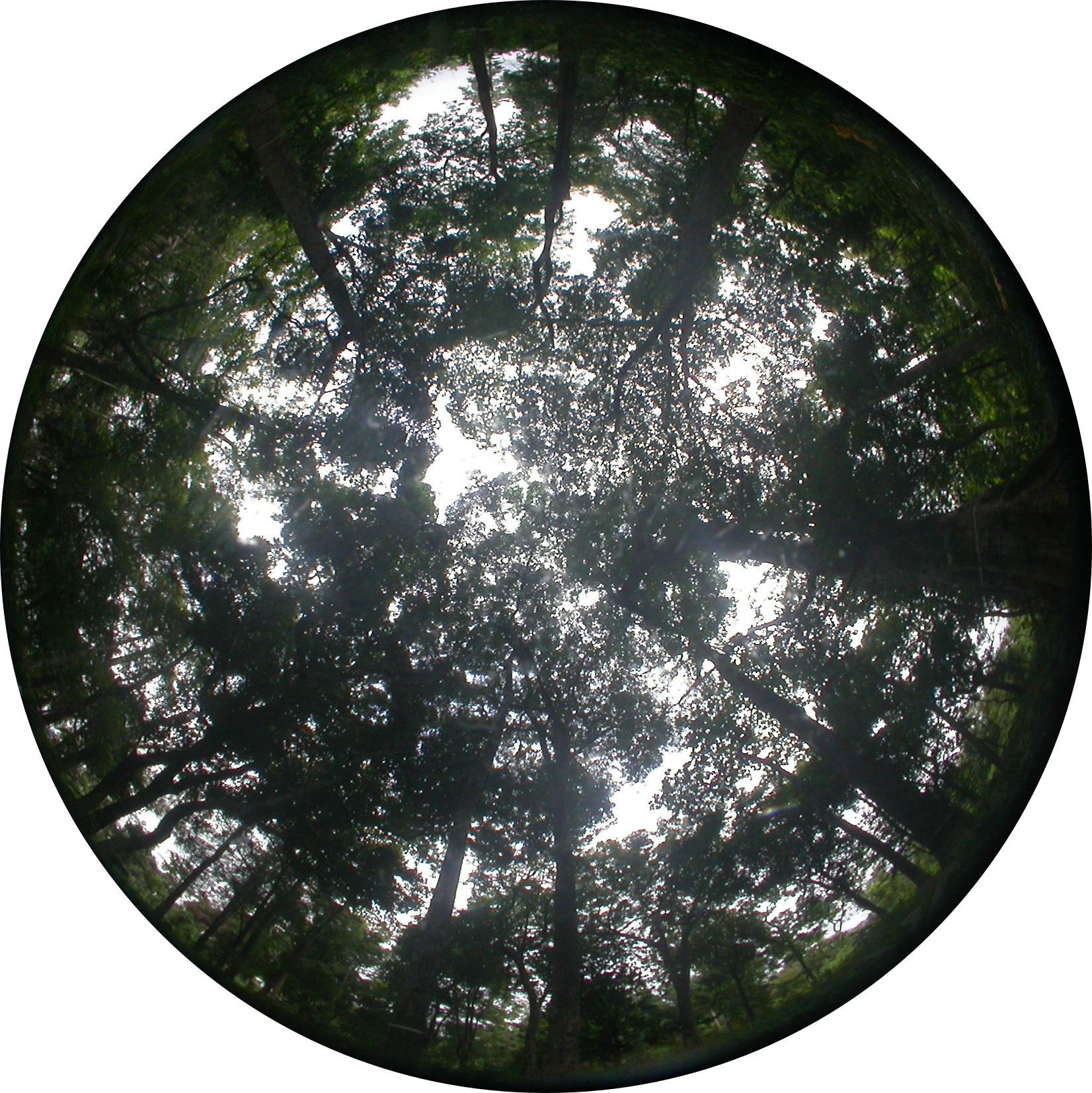

b

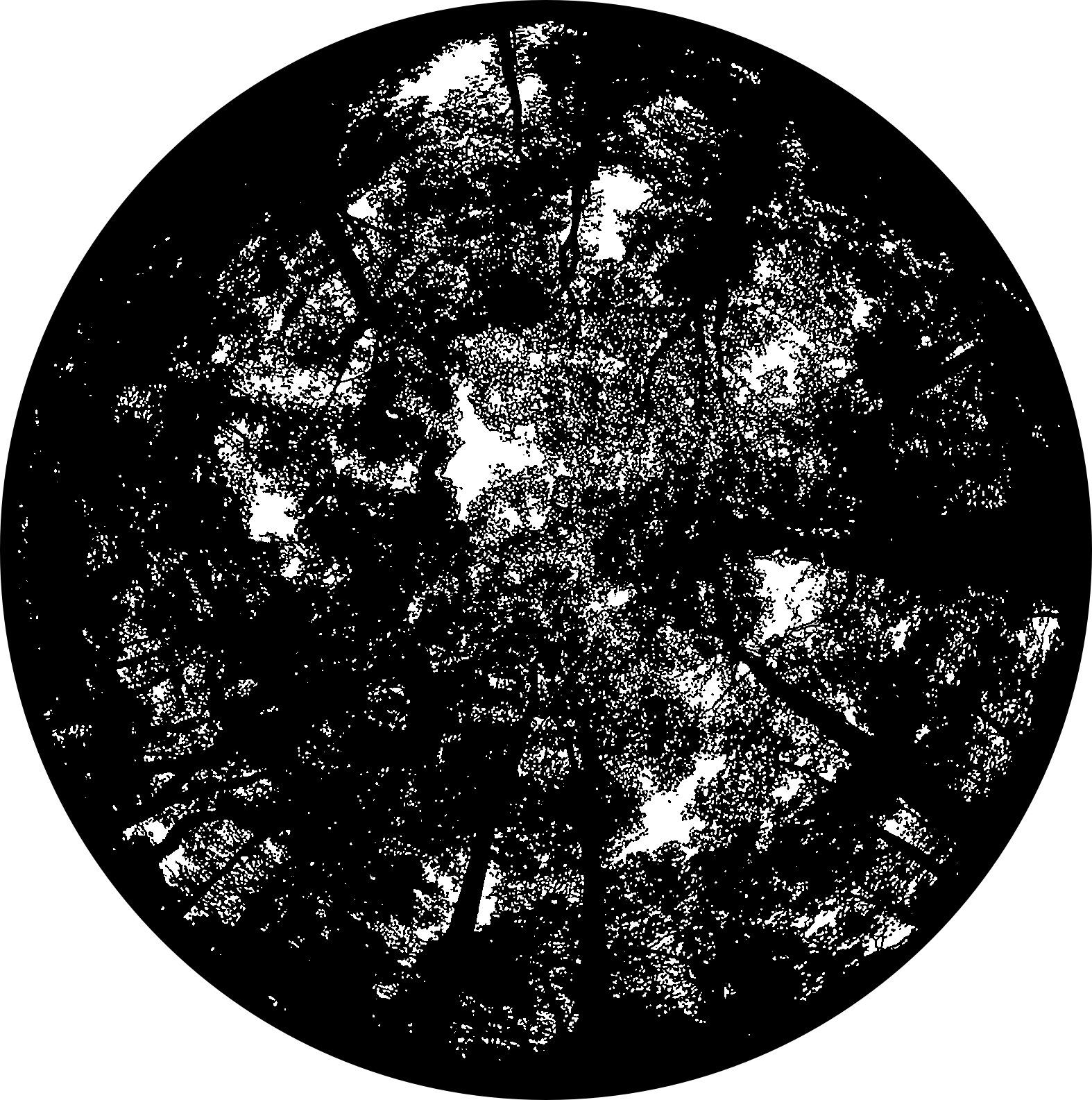

c

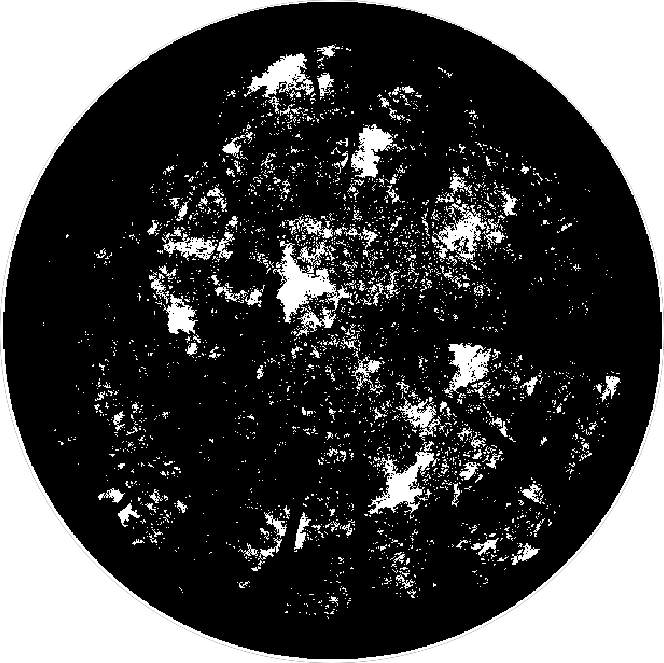

d

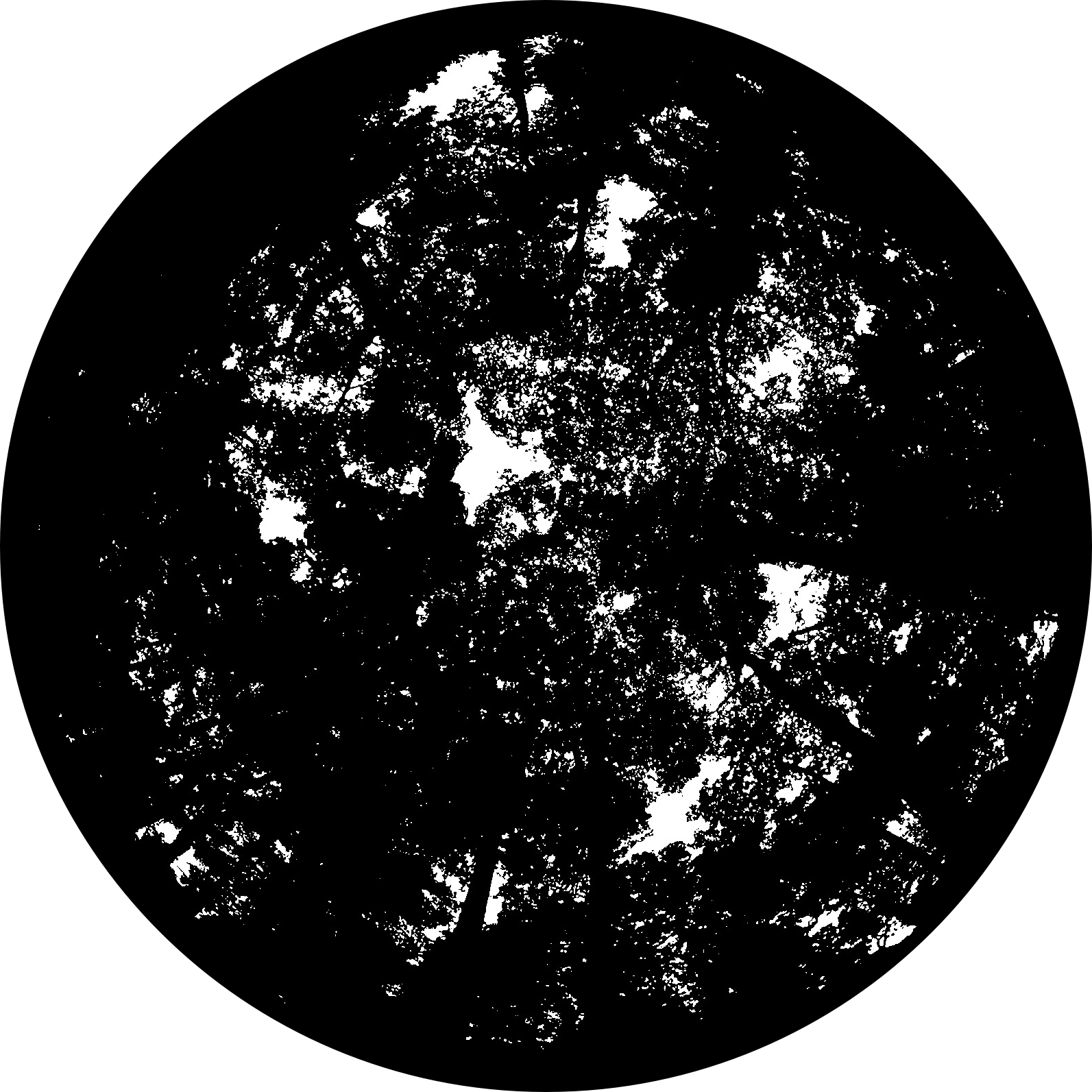

e

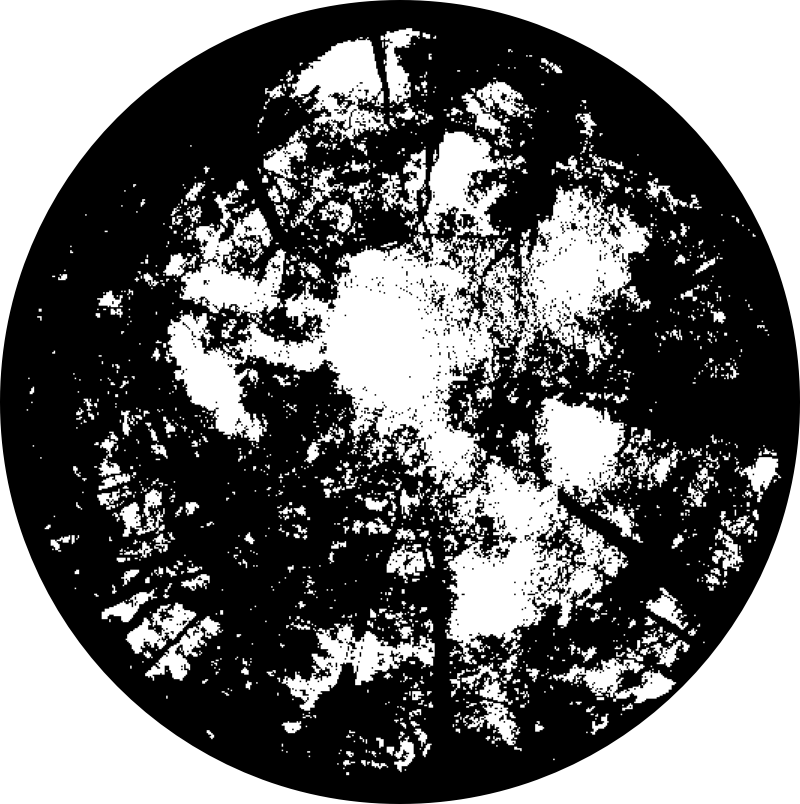

f

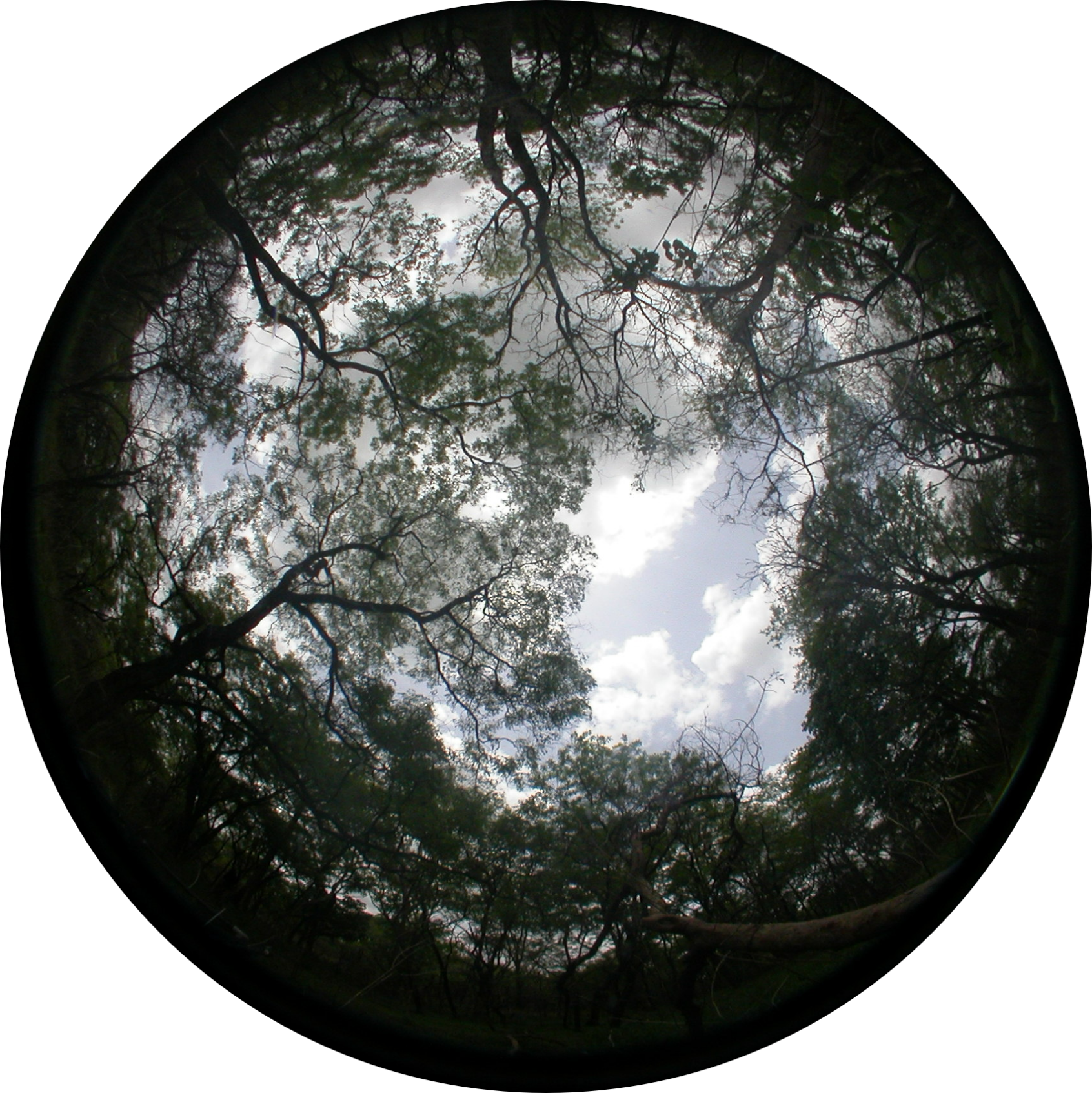

g

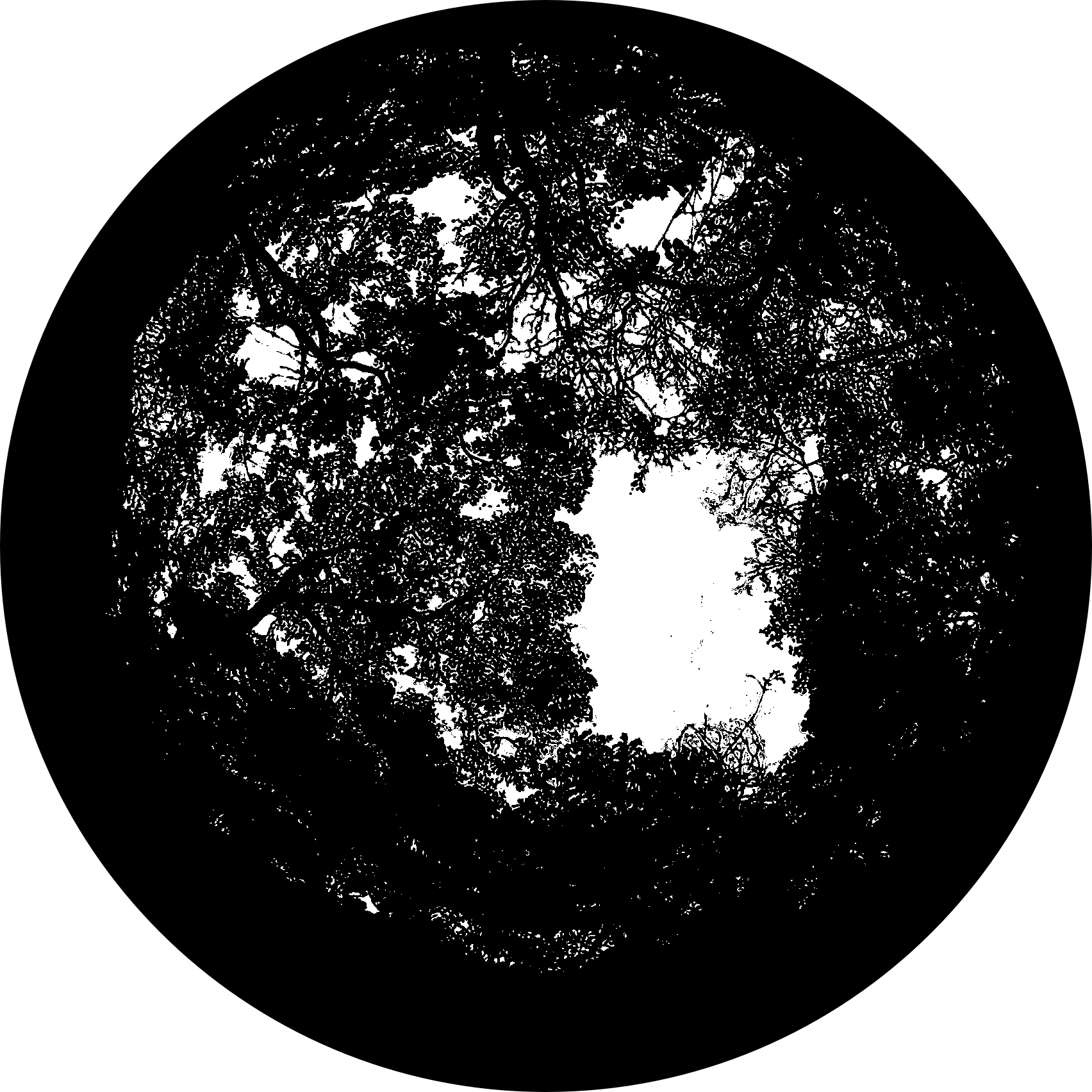

h

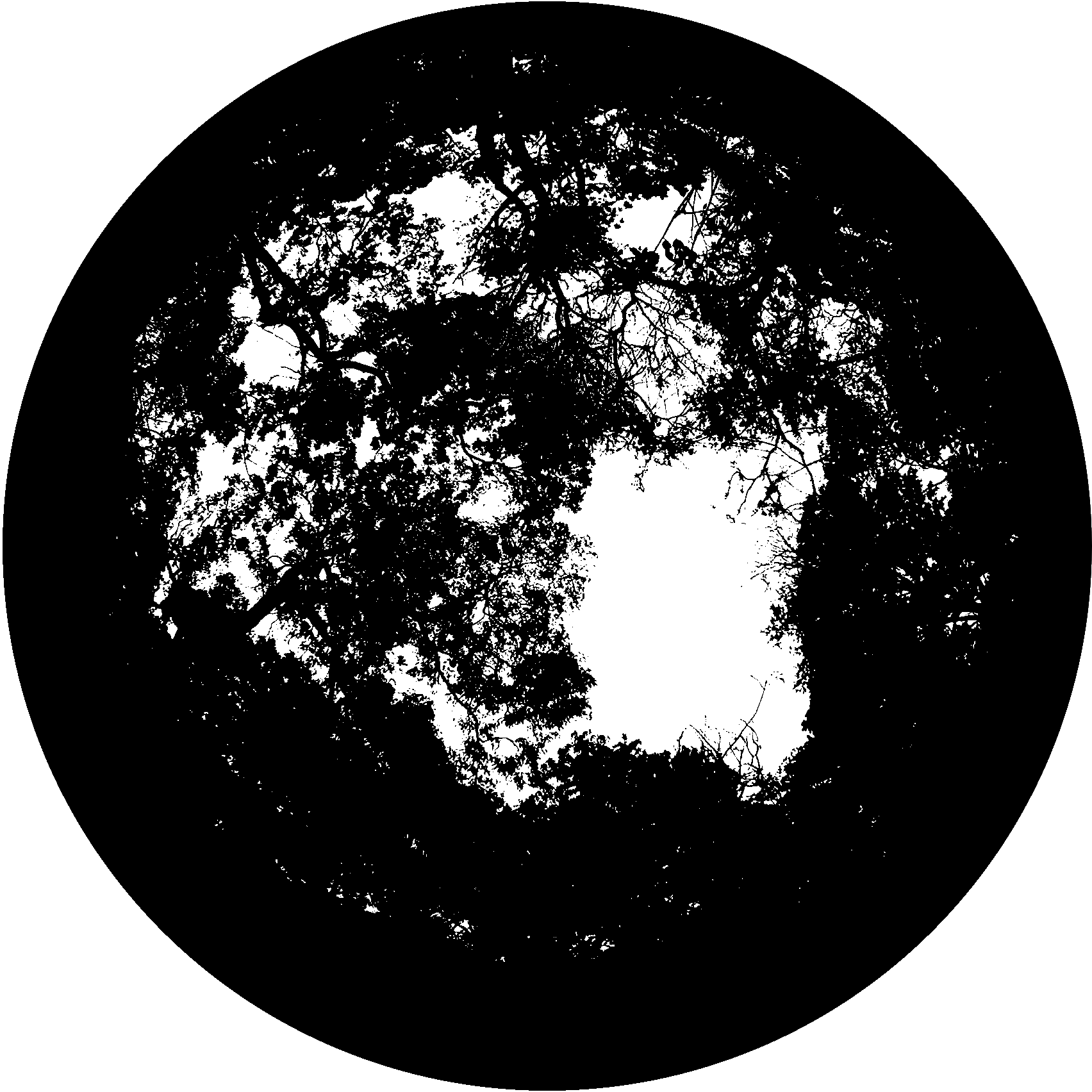

i

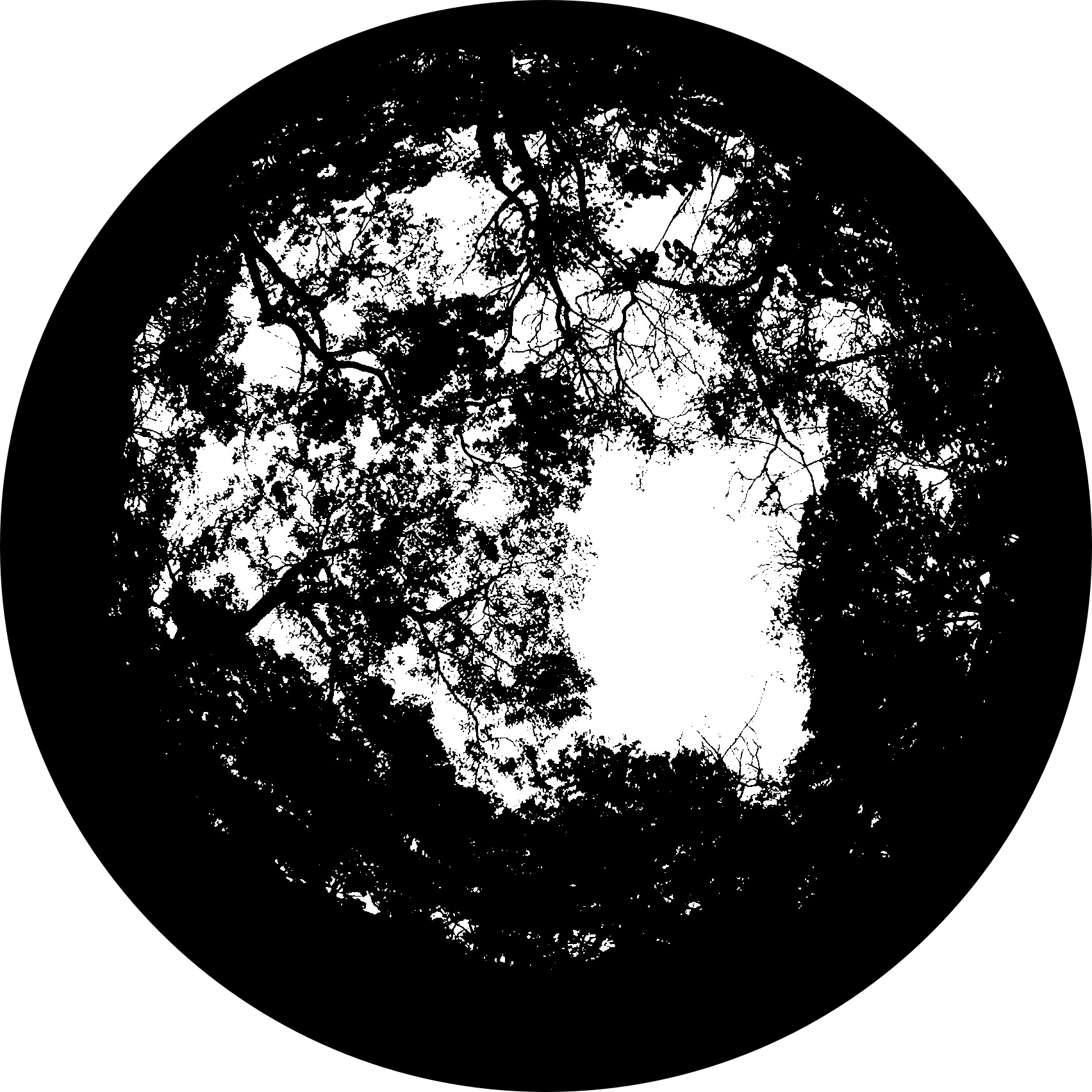

j

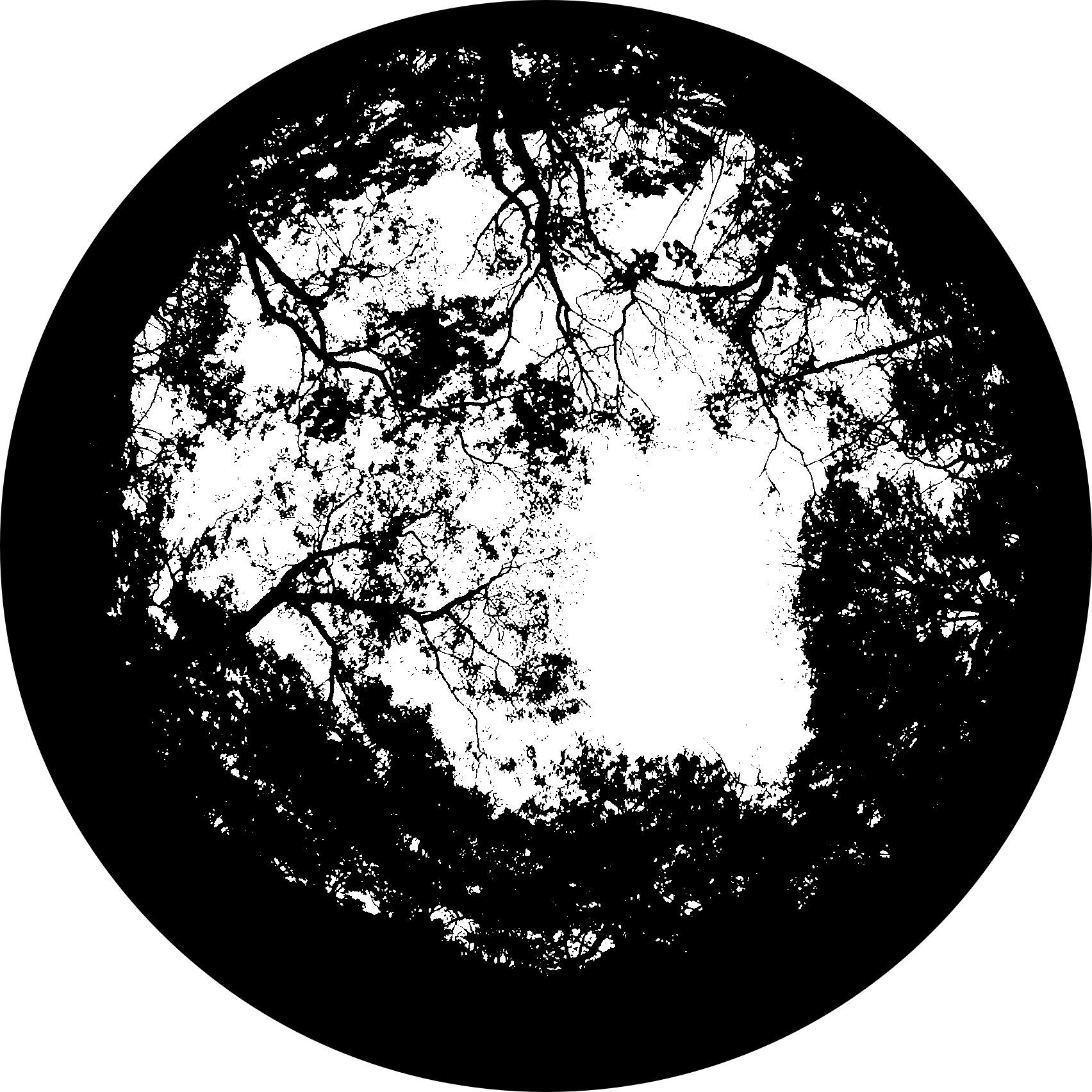

Supplementary Figure 10 Hemispherical canopy photographs and canopy/sky binary images processed by different software. (a, f) Original upward-facing hemispherical canopy photographs from two example plots (a: KOG02, September 2017; f: KOG06, April 2017). Please note that we deliberately picked two challenging photos. The first one is taken under a homogeneous and overcast sky (which is great), but during noon time in the tropics, the sky is extremely bright. The edge of the leaves is not sharp, and the horizon of the photo appears green. The second photo was not taken under a homogeneous sky. (b, g) Canopy/sky binary images generated by ilastik. (c, h) Canopy/sky binary images generated by hemispheR. (d, i) Canopy/sky binary images generated by hemisfer. (e, j) Canopy/sky binary images generated by HemiPy. In panels (b–e, g–j), black pixels represent canopy (leaves + wood) and white pixels represent sky.

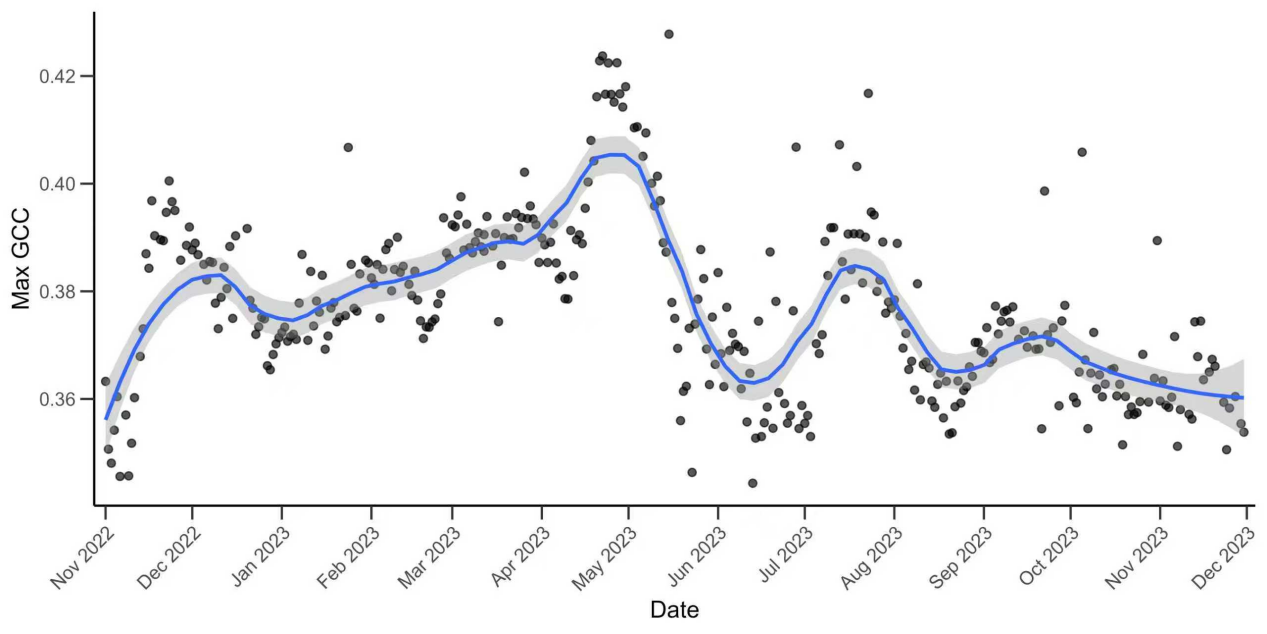

Supplementary Figure 11 Seasonal dynamics of canopy greenness (GCC) at BOB-02. Points show the maximum green chromatic coordinate (GCC) per image date across five canopy regions of interest (ROI1, ROI5, ROI6, ROI7, and ROI8). The blue curve shows a LOESS smooth (span = 0.25) fitted to the time series, and the grey band indicates the 95% confidence interval.

(a)

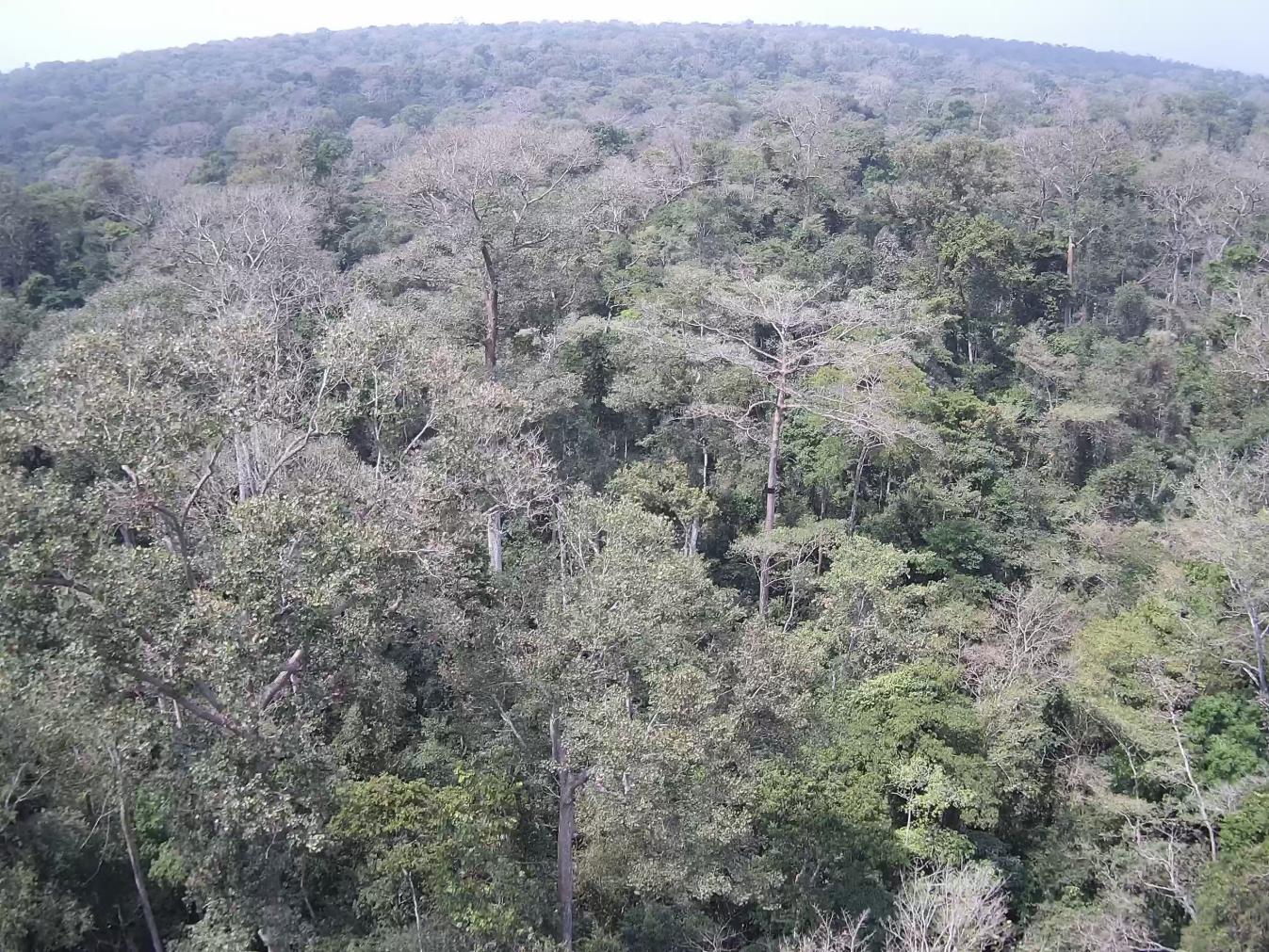

(b)

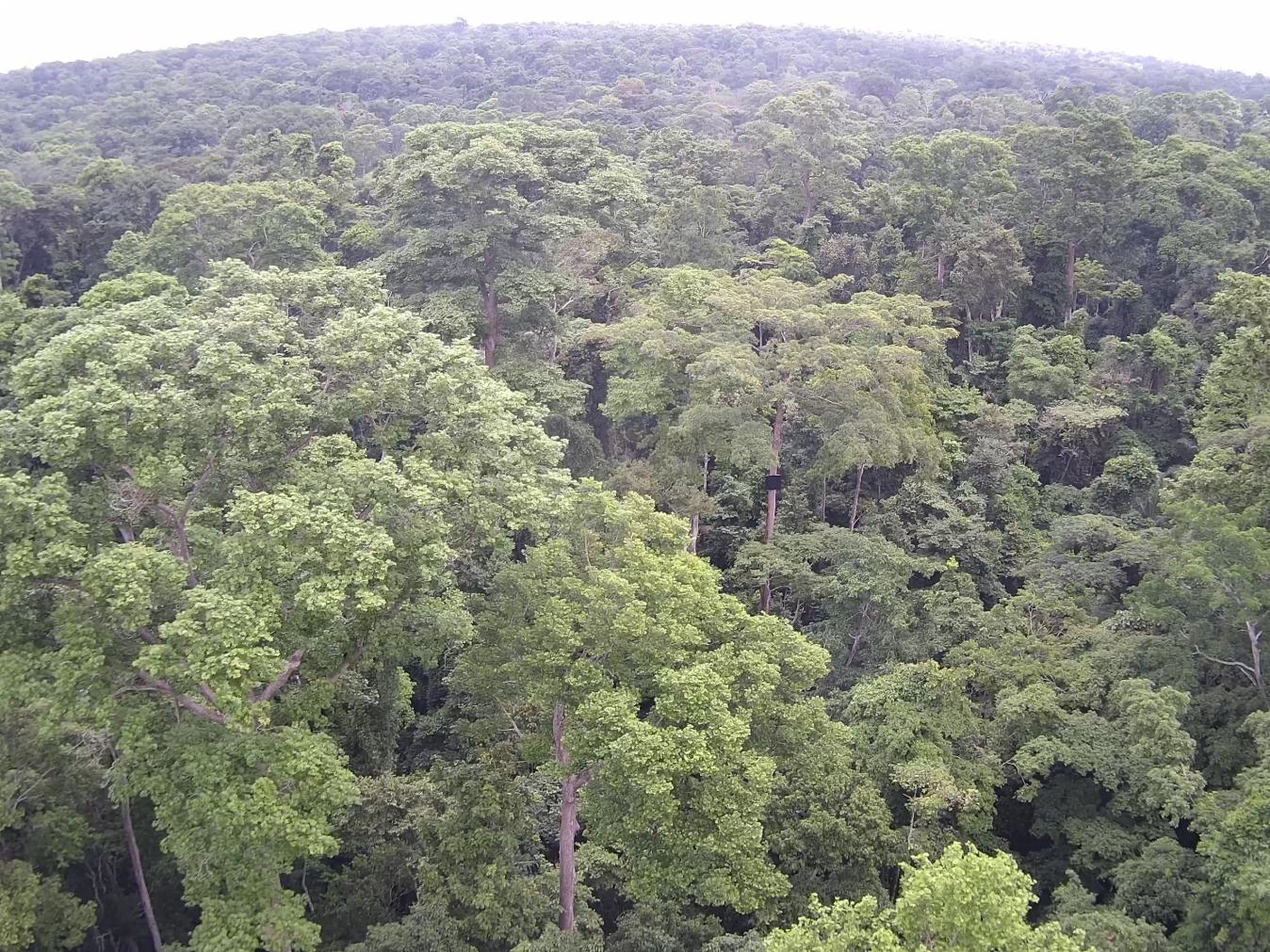

(c)

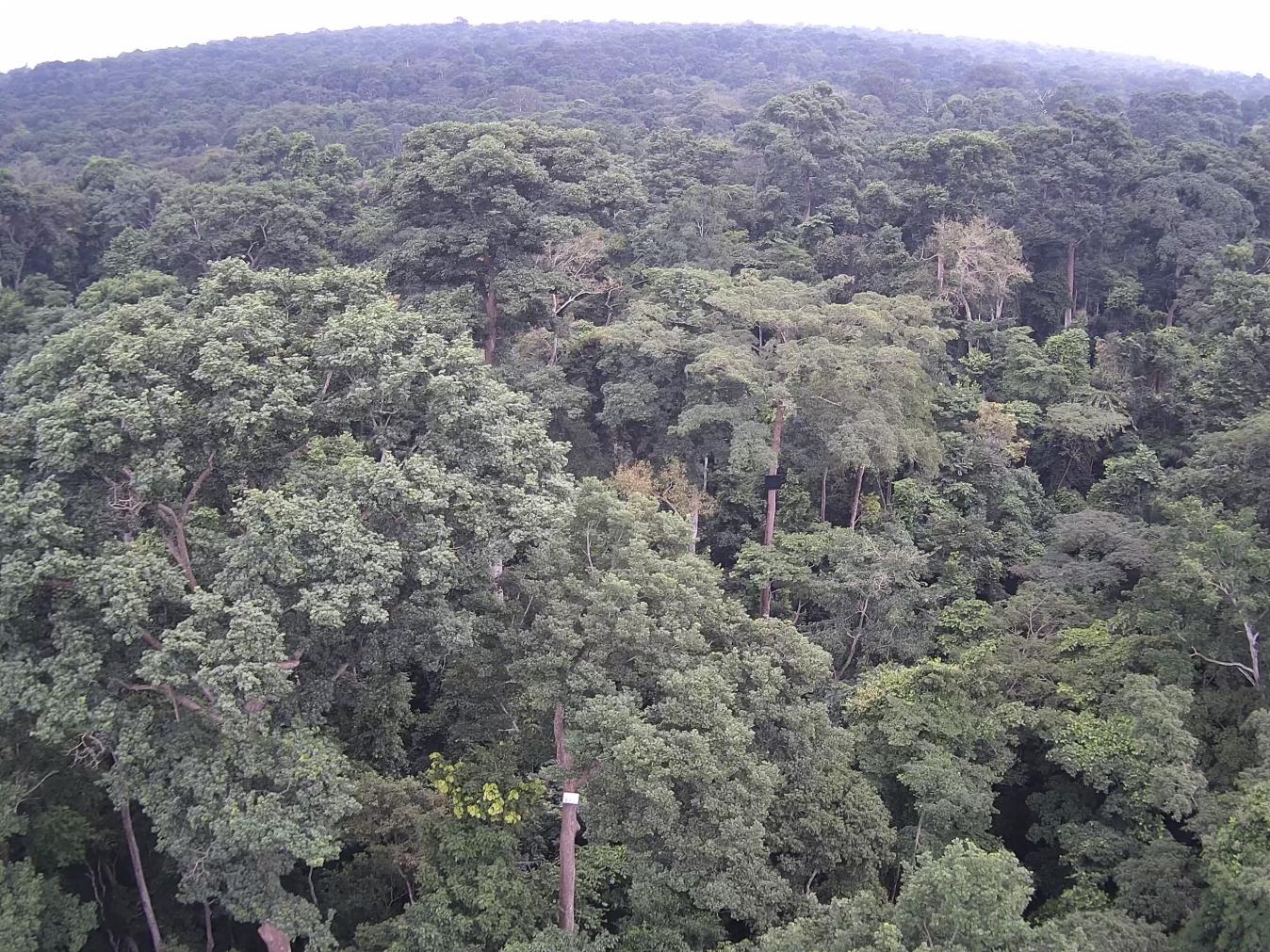

Supplementary Figure12 Example canopy photographs illustrating seasonal variation in GCC at BOB-02 (Bobiri). Representative photographs of the forest canopy at BOB-02 taken on (a) 20 January 2023, (b) 31 May 2023, and (c) 30 August 2023. These images provide qualitative visual context for the seasonal changes in canopy greenness captured by the GCC time series.

March 2017

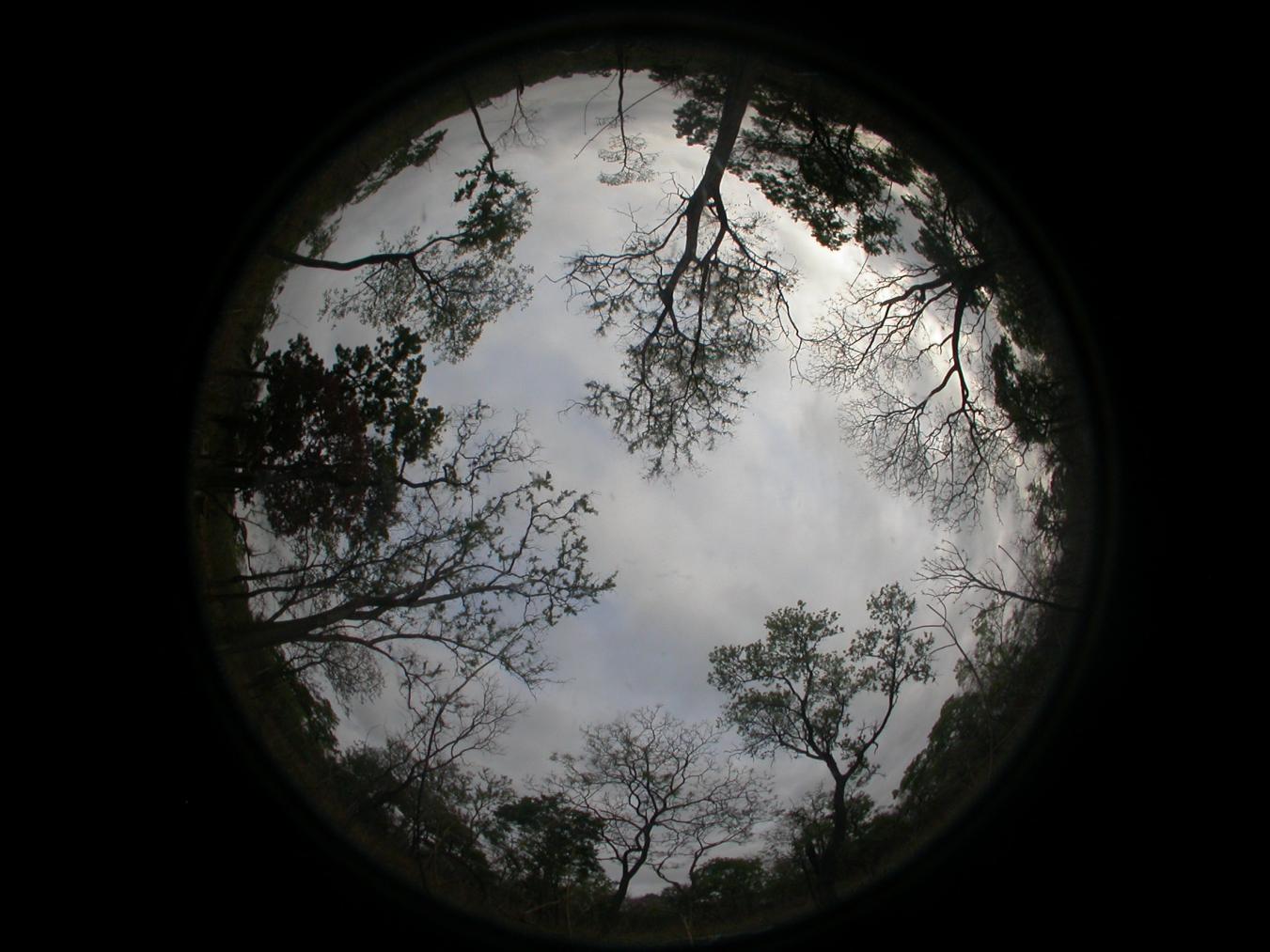

April 2017

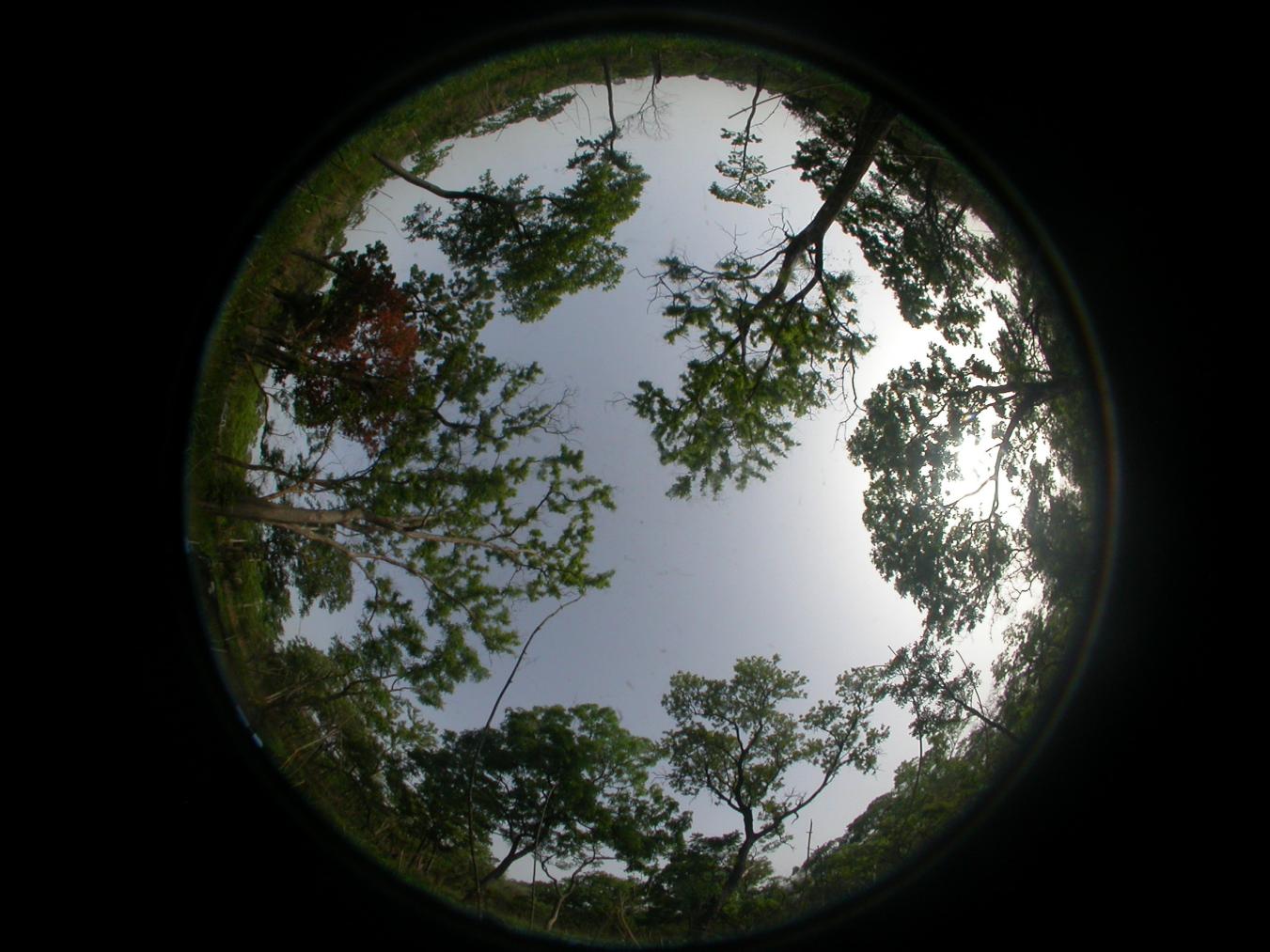

May 2017

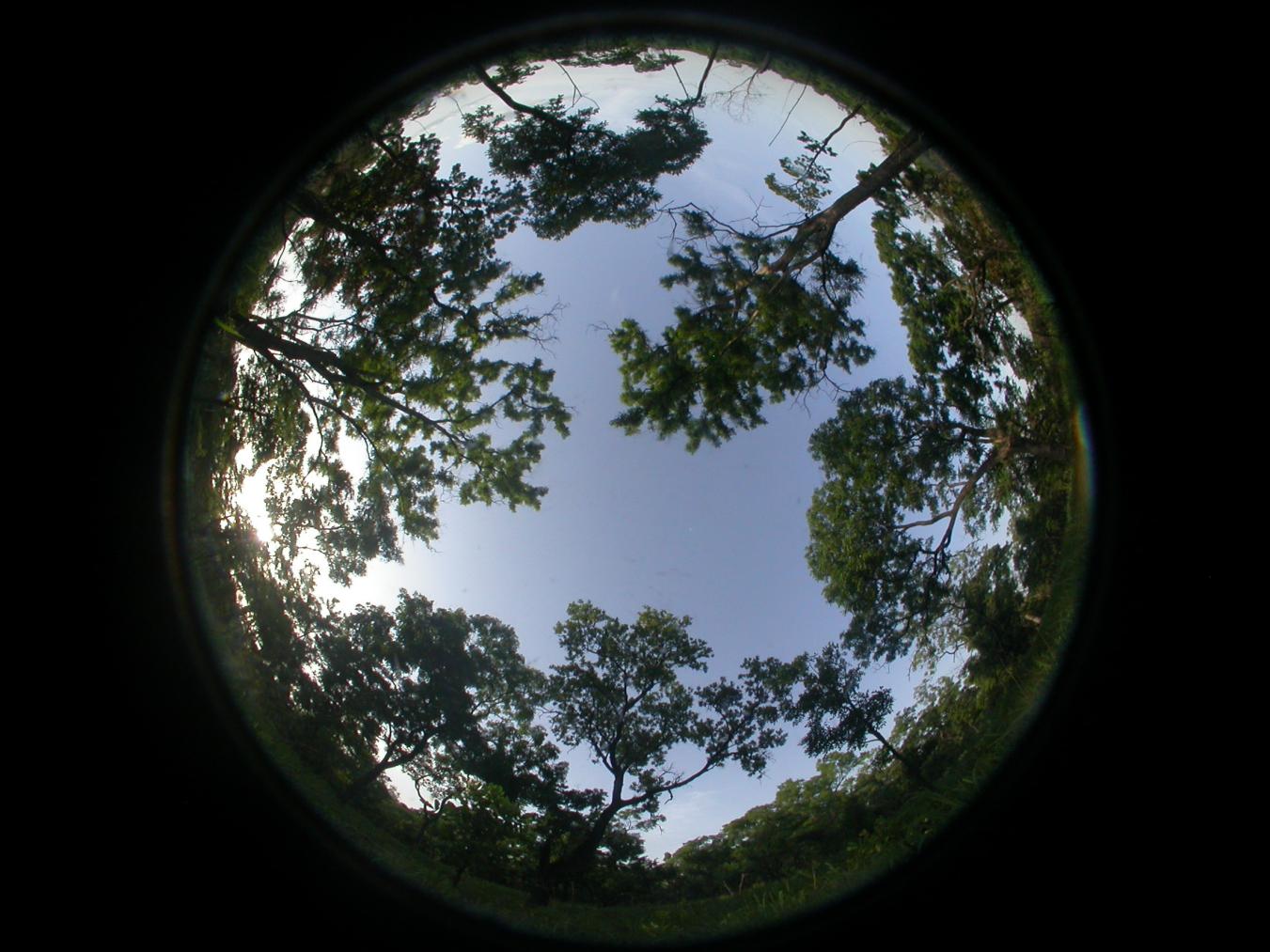

June 2017

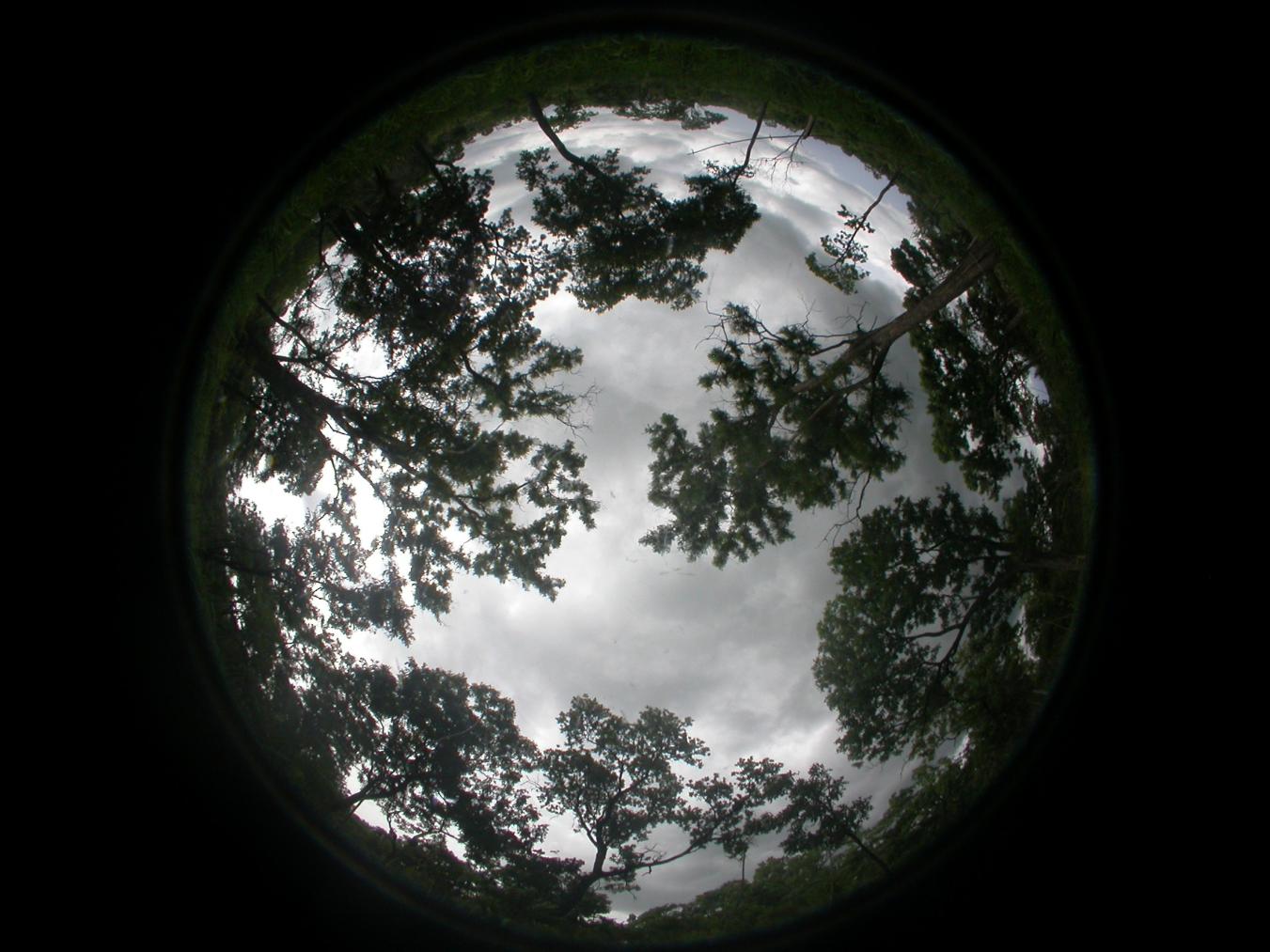

July 2017

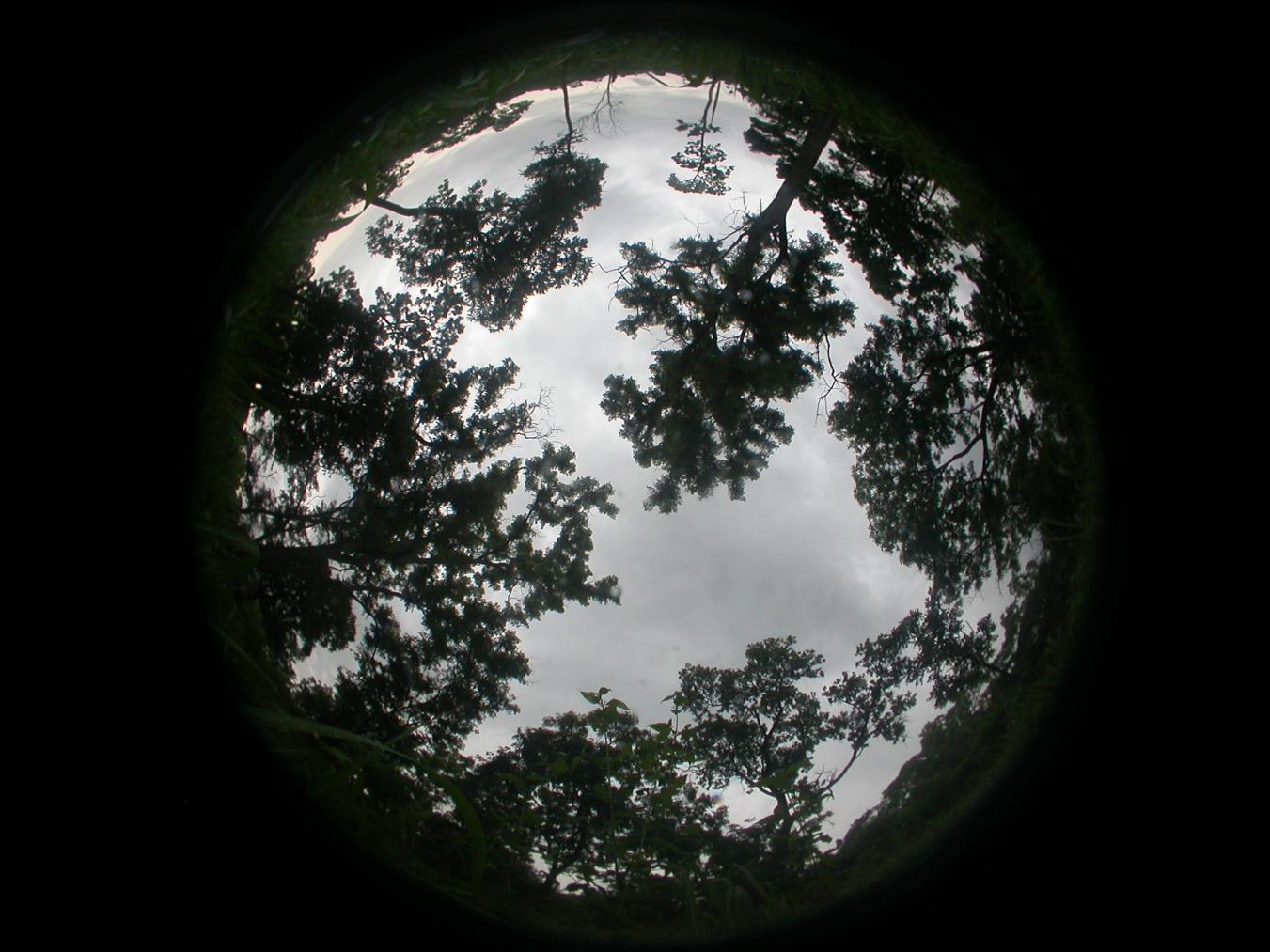

August 2017

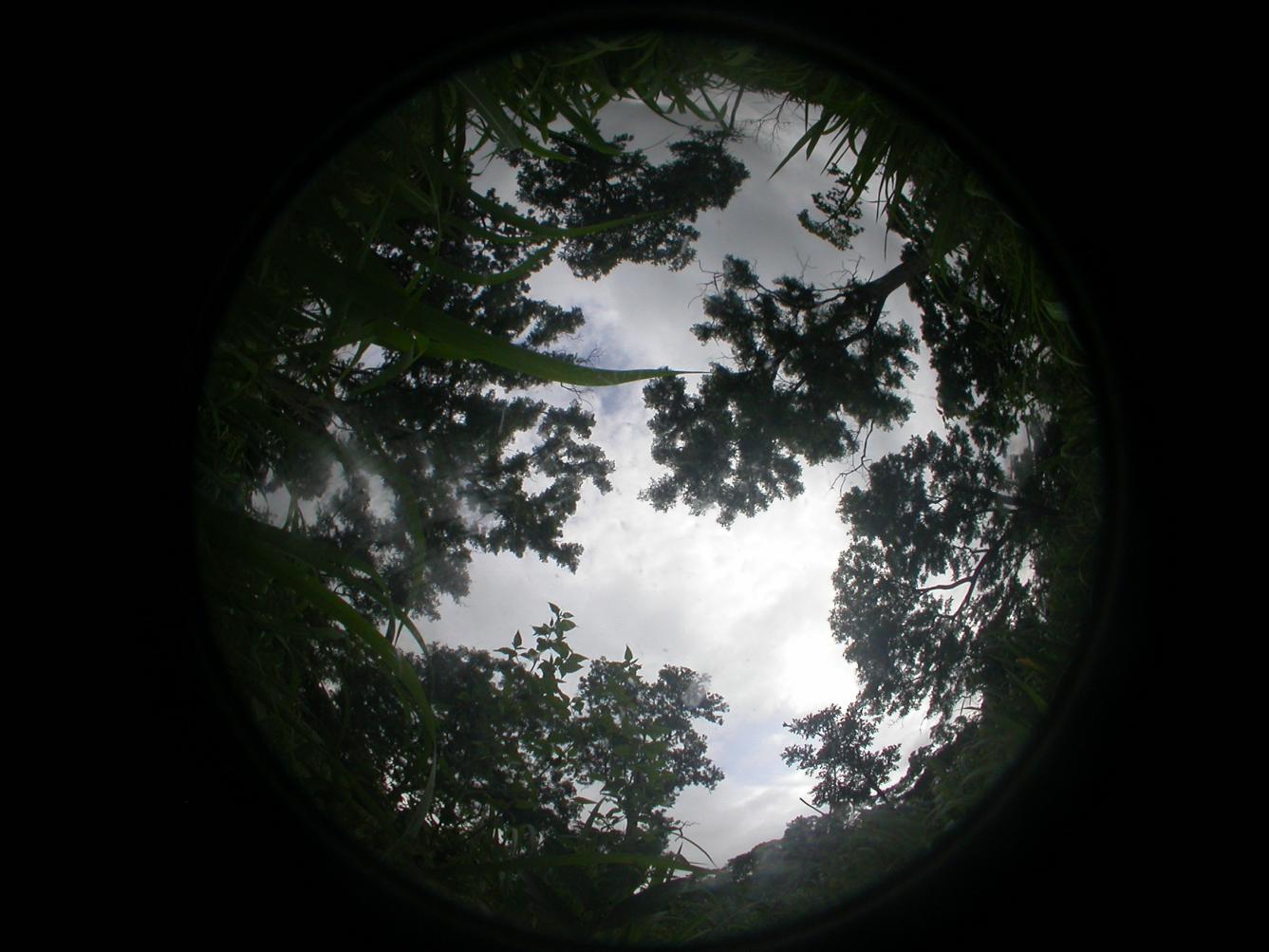

October 2017

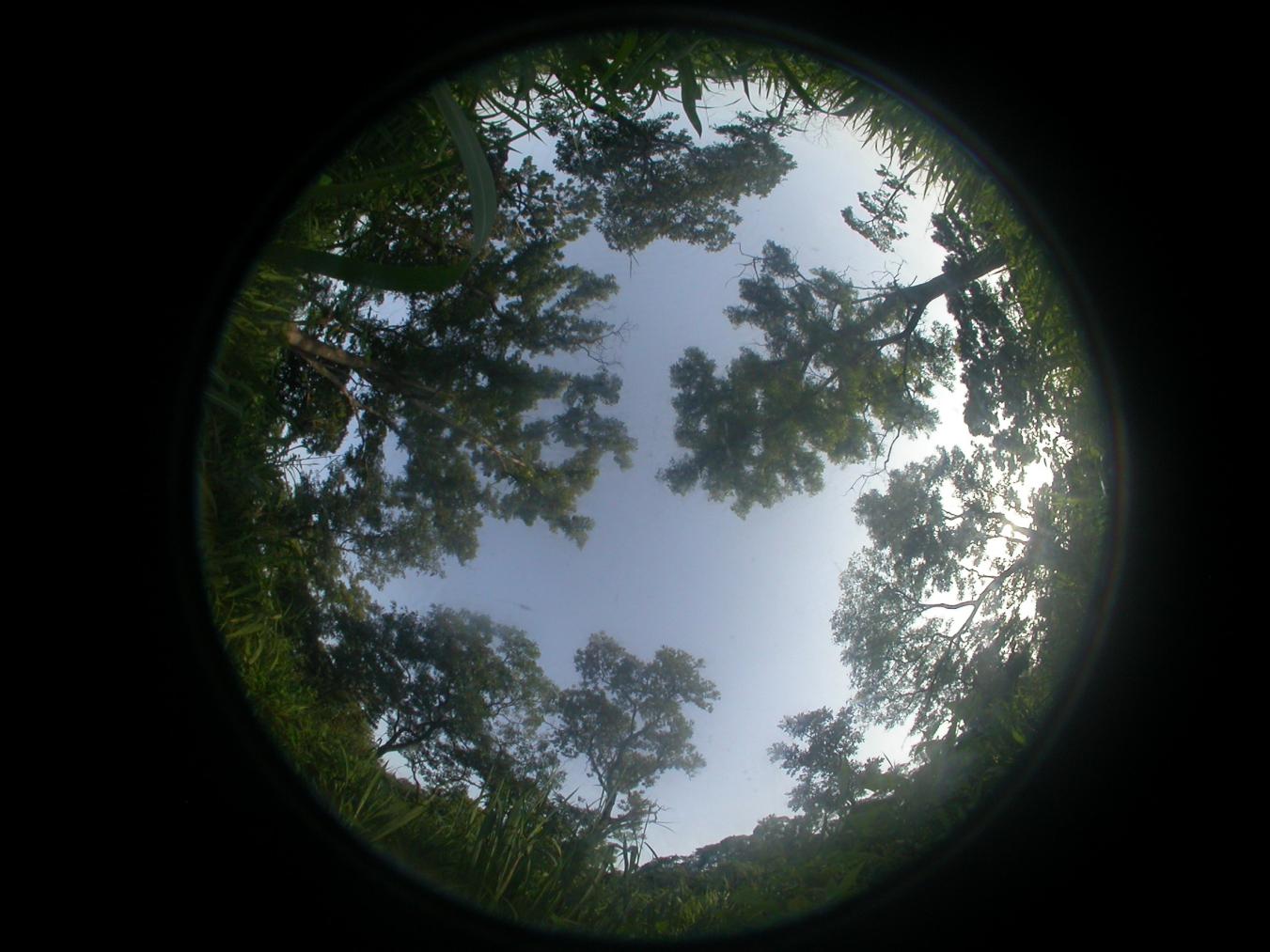

December 2017

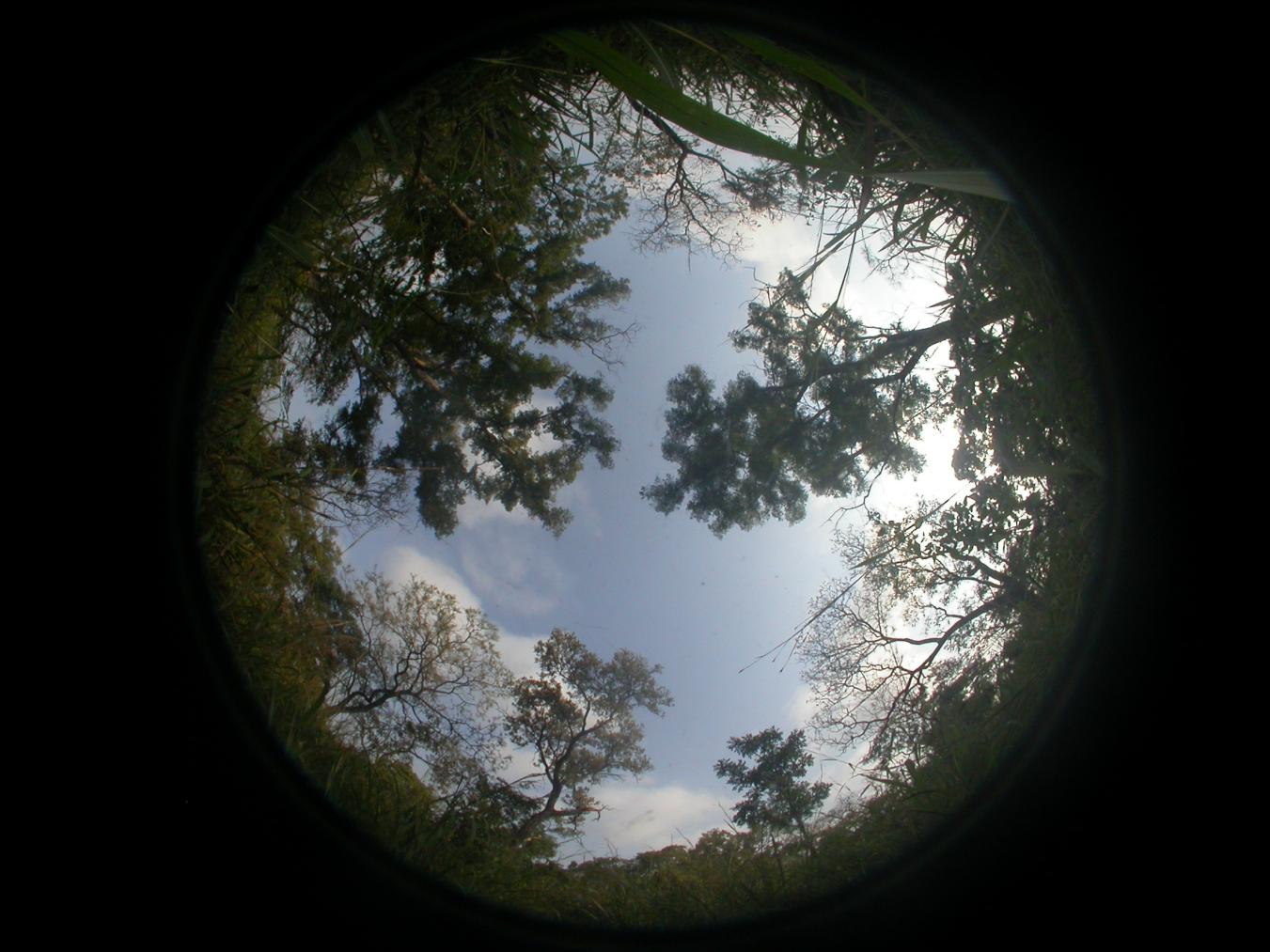

January 2018

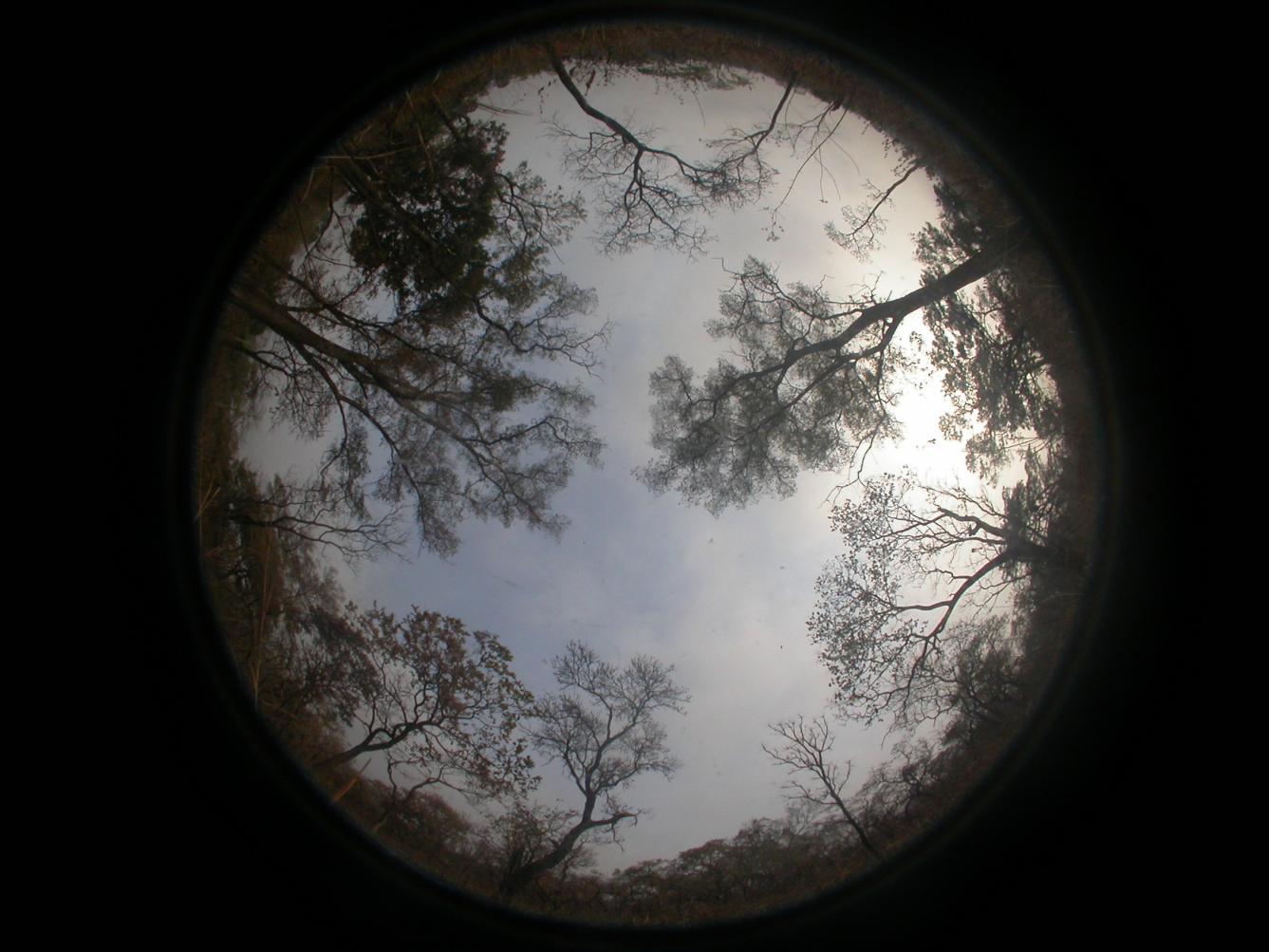

February 2018

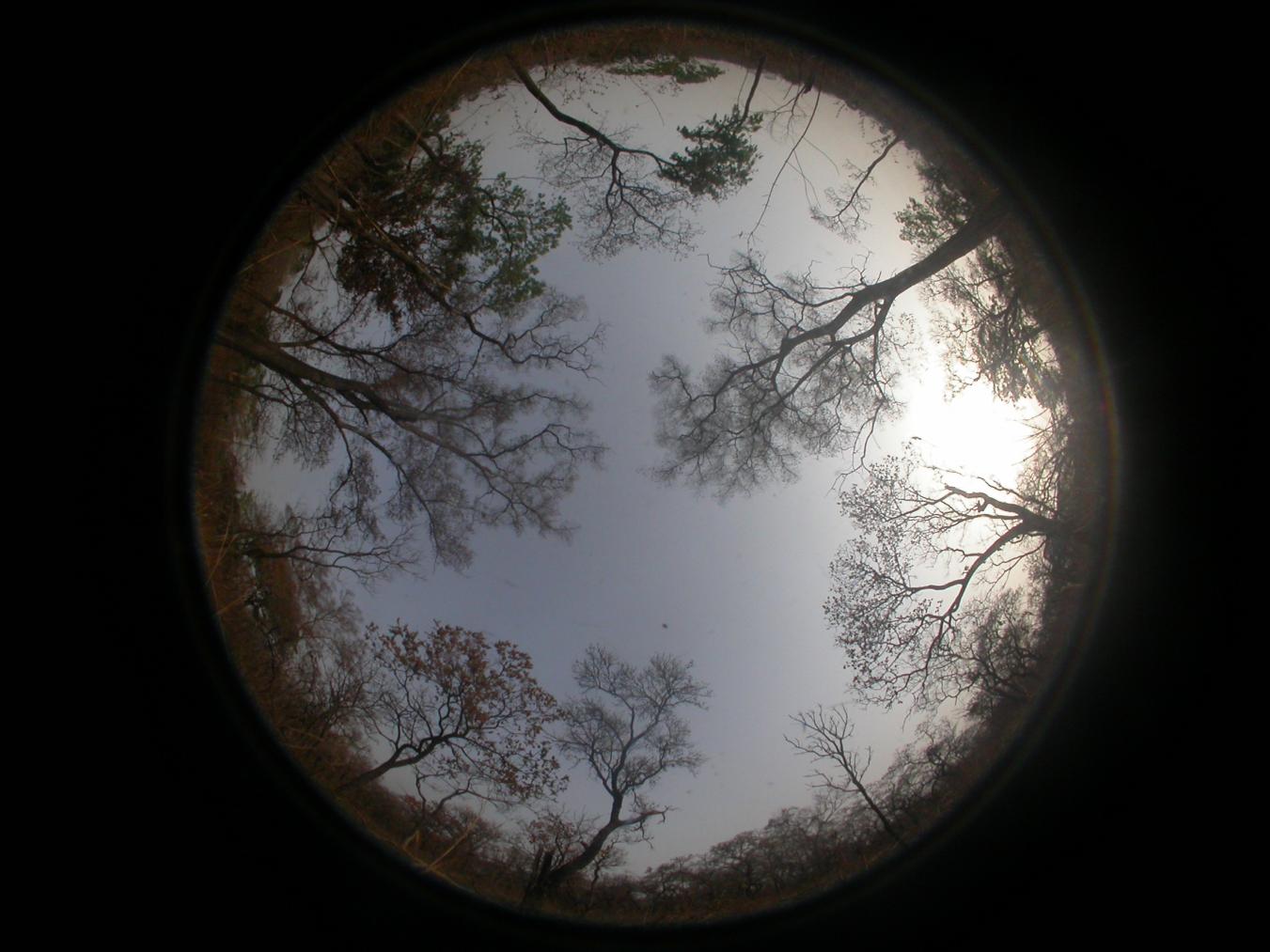

Supplementary Figure 13 Monthly hemispherical canopy photographs at KOG-05. Representative fisheye (hemispherical) images acquired at KOG-05 from March 2017 to February 2018 are shown to illustrate seasonal variation in canopy condition over the annual cycle.

a

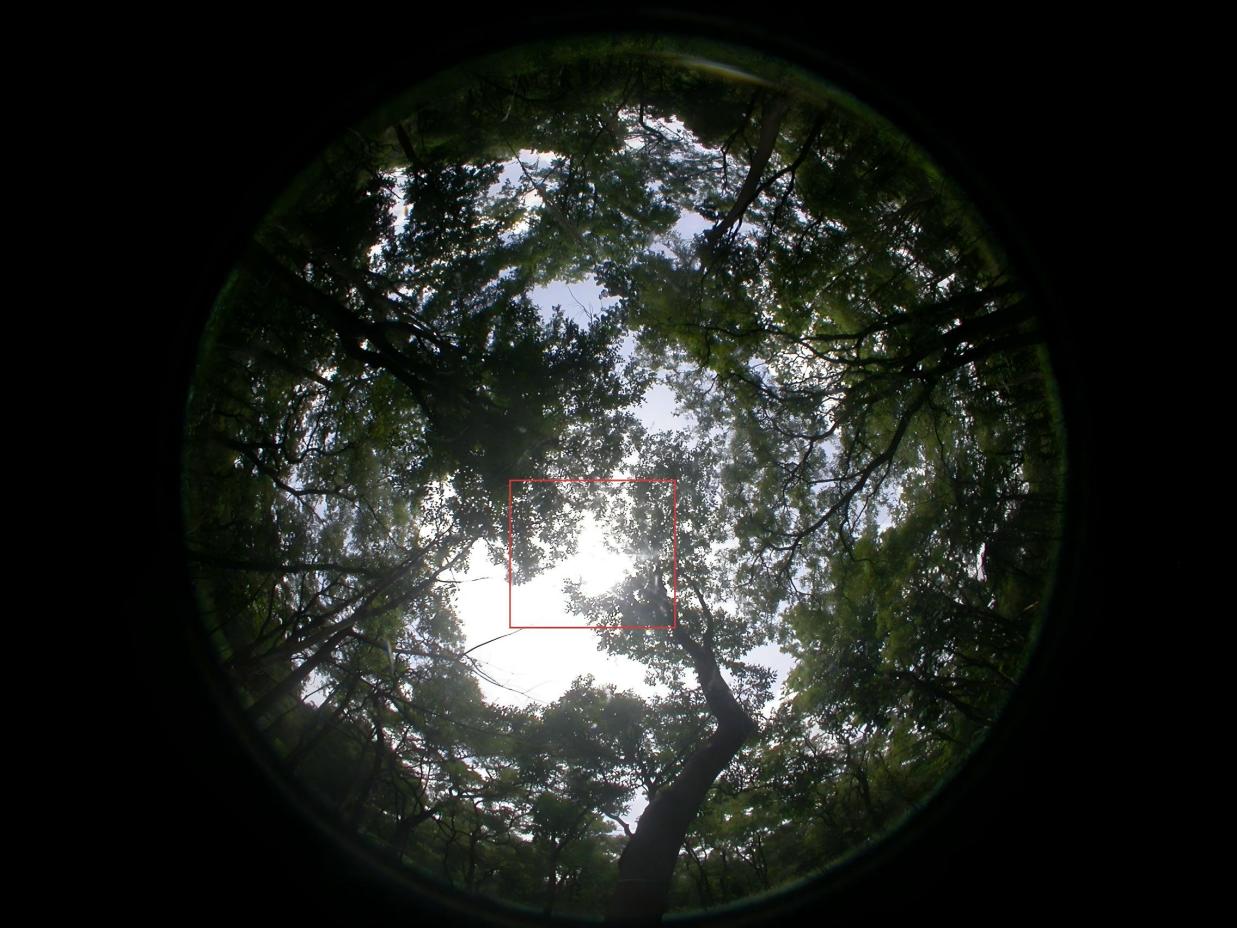

b

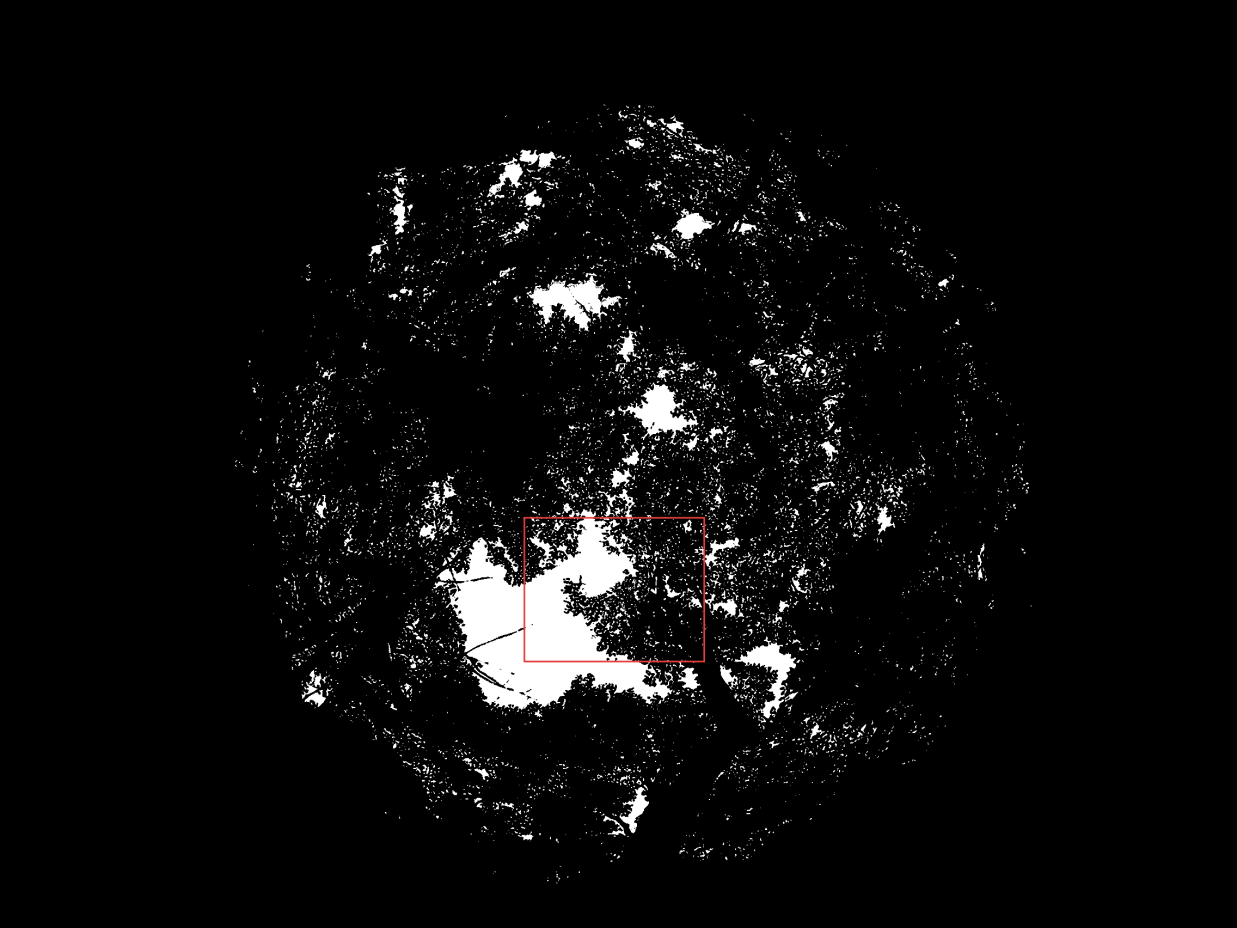

c

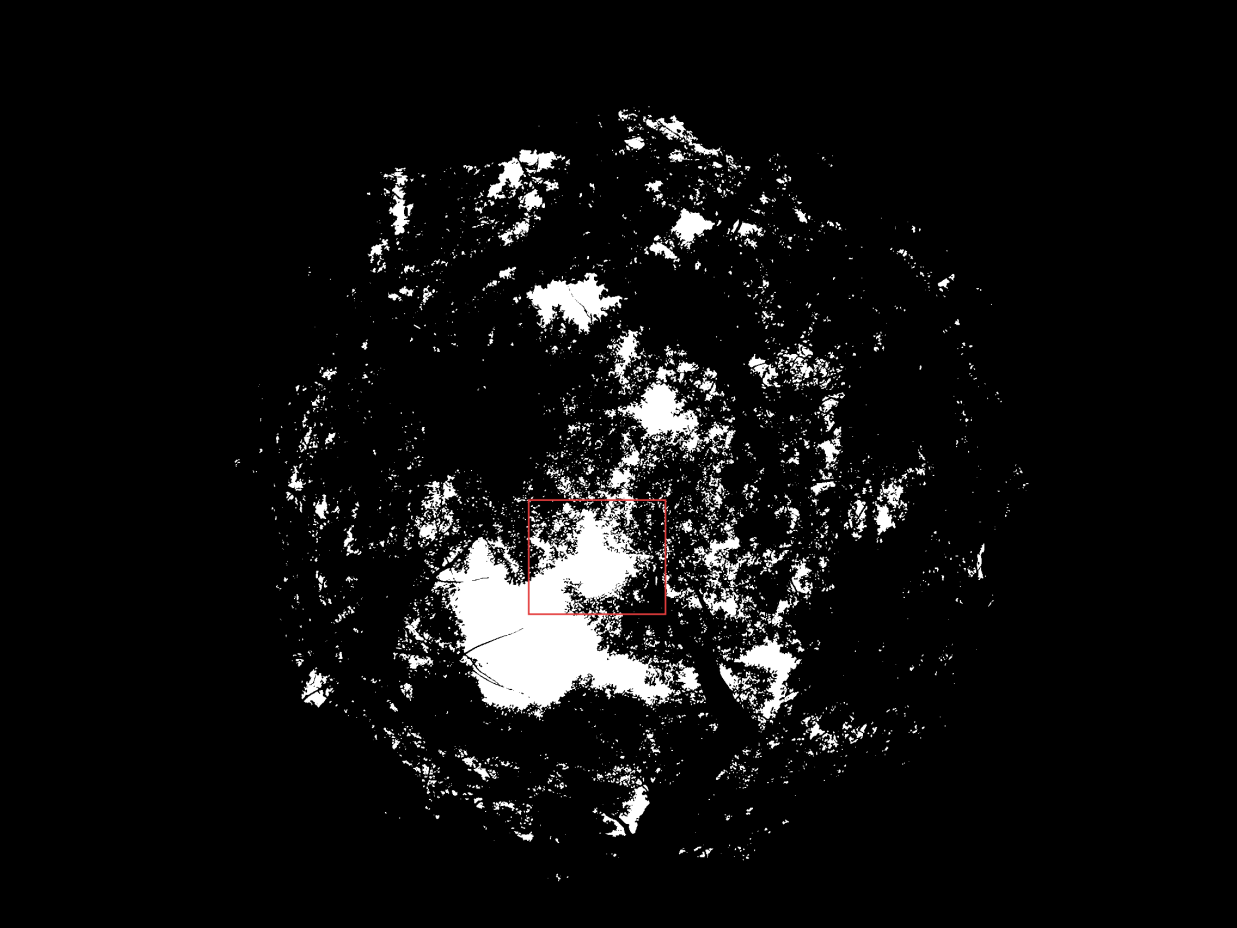

d

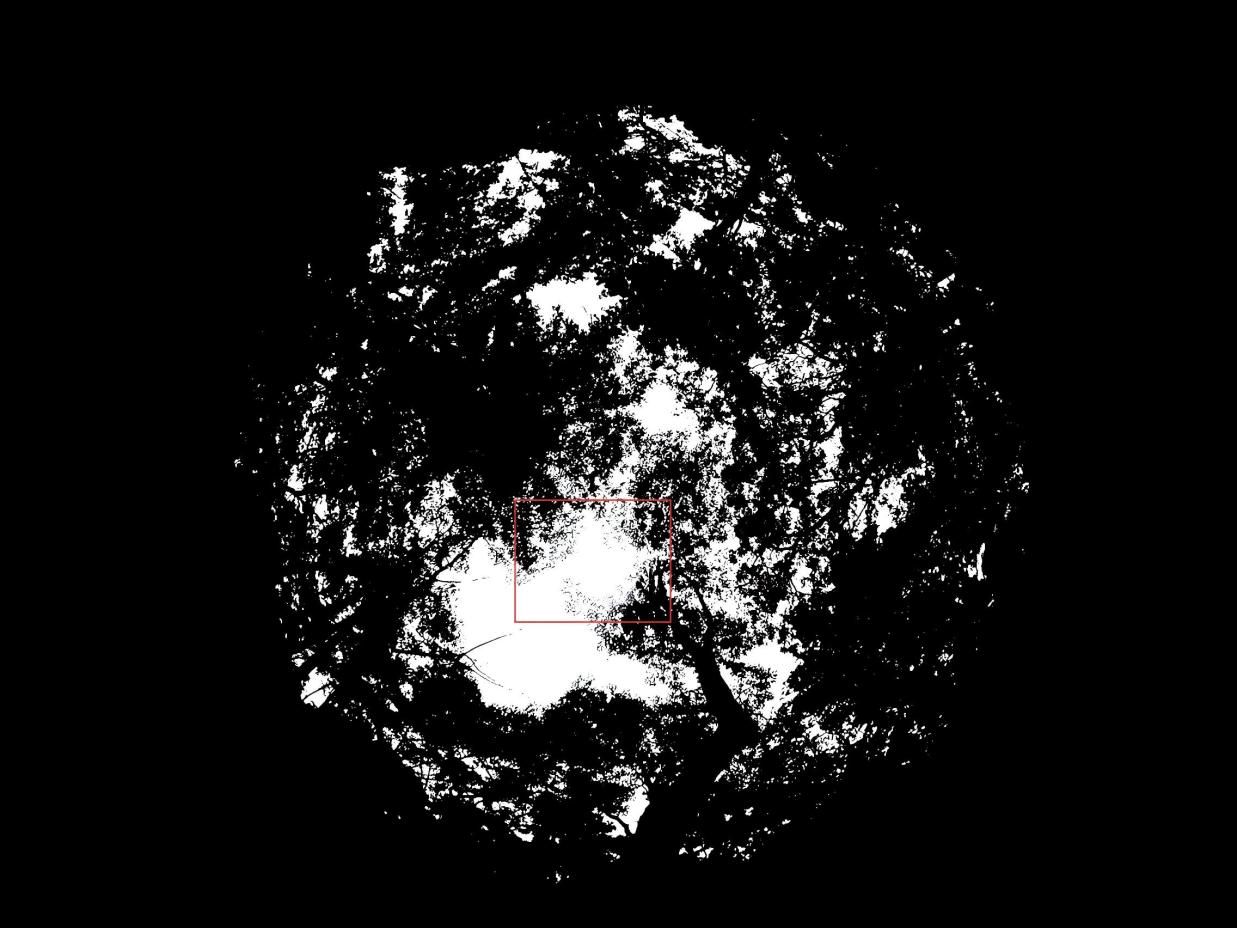

e

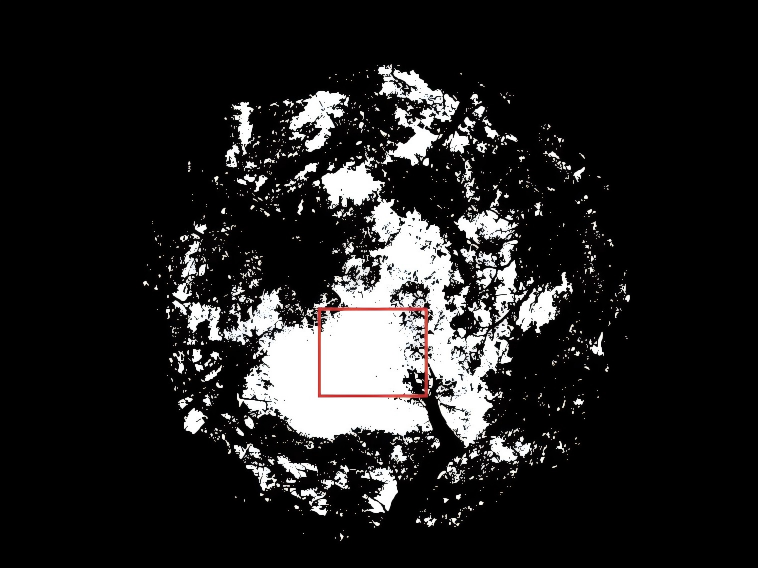

Supplementary Figure 14 Robustness of canopy/sky segmentation to “bright-gap bleeding” artefacts across software. (a) An upward-facing hemispherical canopy photograph from the testing set illustrating a hazy overexposure artefact. (b–e) Corresponding canopy/sky binary images generated by ilastik (b), hemispheR (c), hemisfer (d) and HemiPy (e). Black pixels represent canopy (leaves + wood) and white pixels represent sky.
