## Supplementary protocol for "An AI-based and coding-free integration for forest Leaf Area Index calculation"

### Taking Hemispherical photo in the field

The accuracy of exposure settings is crucial when using digital hemispherical photography (DHP) to measure canopy structure. Chen et al. (1991) was among the first to highlight that automatic exposure can introduce bias by reducing contrast between the sky and canopy. Later, Zhang et al. (2005) recommended adjusting manual settings—such as increasing exposure by 1–2 stops compared to open-sky conditions—to enhance measurement accuracy. Numerous subsequent studies have explored methods for camera exposure parameterization (highlighted in red). However, a recent study acknowledged that in practice—particularly in tropical forests—manual benchmarking against open-sky conditions is often impractical due to time and accessibility constraints (Jiao et al., 2023). The study proposed an innovative solution: capturing multiple images at varying exposures and merging them using image fusion techniques (e.g., Sony’s auto HDR function if you own Sony camera) to optimize canopy detail. While the researchers demonstrated the effectiveness of this approach, implementation remains challenging: no user-friendly software exists yet, and the requirement to capture multiple shots per location is time-intensive.

Here’s our practical advice: If you have a strong grasp of those papers and sufficient time, we recommend setting manual exposure or experimenting with multi-exposure fusion. However, if these methods are impractical in your field setting, don’t be overly concerned—just avoid extreme cases. For instance ensure the sky isn’t overly bright (Figure 1), as this can mask leaf details. For modern cameras (especially Sony models), enable Auto HDR and Auto Exposure, but familiarize yourself with brightness adjustments in your camera settings for necessary finetuning. Aim for good exposure, but avoid letting perfectionism hinder fieldwork progress. This is especially true if you’re using our AI-based workflow, as minor exposure variations have minimal impact on LAI results. Ideal lighting conditions include diffuse light, such as overcast skies or dawn/dusk periods (Chen et al., 1991). That said, in tropical forests—where lighting can be exceptionally dim—the environment may become too dark for clear shots. In such cases, prioritize shutter speed and focus—image clarity is far more critical than exposure. Discard any blurry images during post-processing (Figure 3), and never use a flash in low-light conditions.

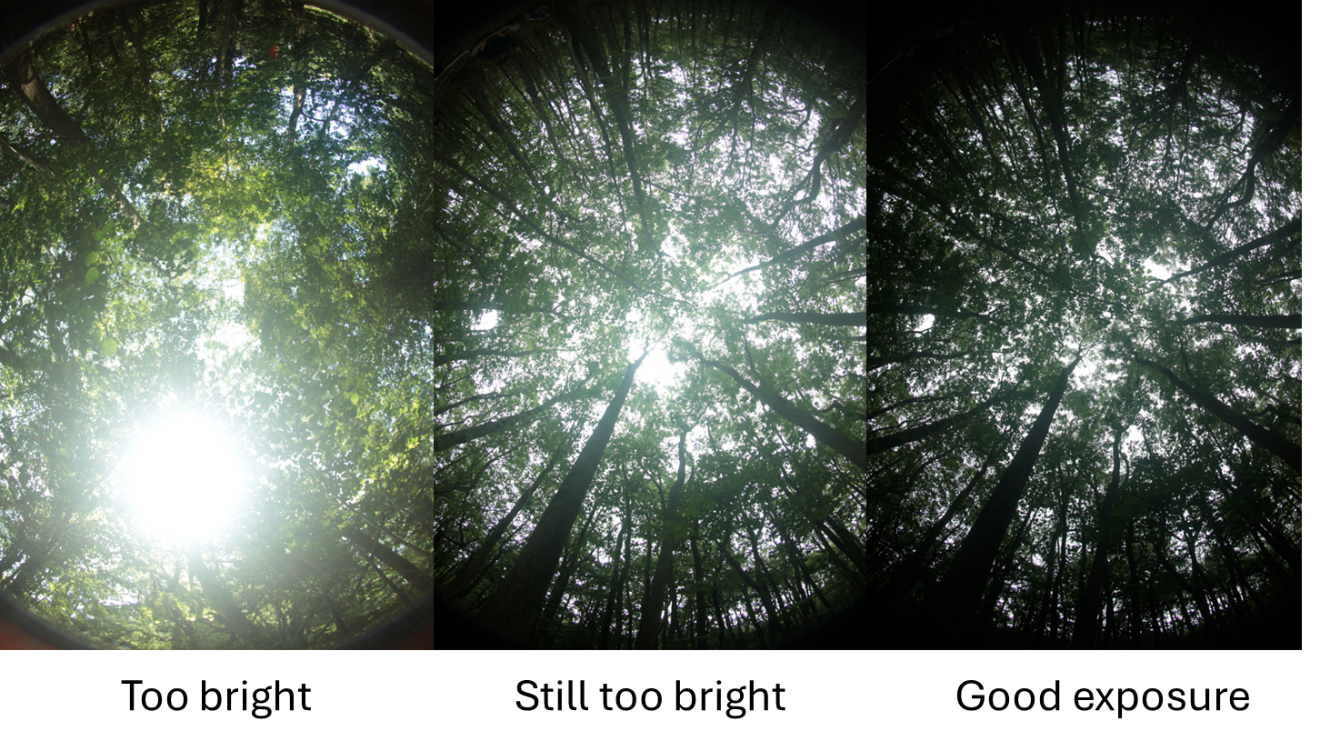

*Figure 1*

It is essential to use a camera equipped with a circular fisheye lens that provides a full 180-degree field of view (so when pointed upright, the photo’s edges capture the ground). Avoid diagonal fisheye lenses, as they typically offer only a 180-degree view along the diagonal, resulting in incomplete canopy coverage. If purchasing a dedicated camera is not feasible, a smartphone such as an iPhone can serve as an alternative. However, you must attach a 180-degree circular fisheye lens to your device and verify that it is truly circular (not diagonal).

For fieldwork in tropical forest plots measuring 1 hectare, the standard practice is to capture hemispherical photographs at 25 evenly distributed points each month (Xie et al., 2023). After collecting the initial dataset, you can experiment by analyzing subsets of these images to determine whether a reduced number of photographs still produces consistent LAI estimates, which may allow for fewer photos in future surveys. For camera orientation, mount the camera on a tripod and ensure it is perfectly level. The lens should point directly upward, and aligning the camera to magnetic north helps ensure consistent data analysis. Permanently mark each photo location point so the same points can be reliably relocated during subsequent surveys. In tropical forests, additional equipment such as a tree lopper and sickle may be required (Figure 2).

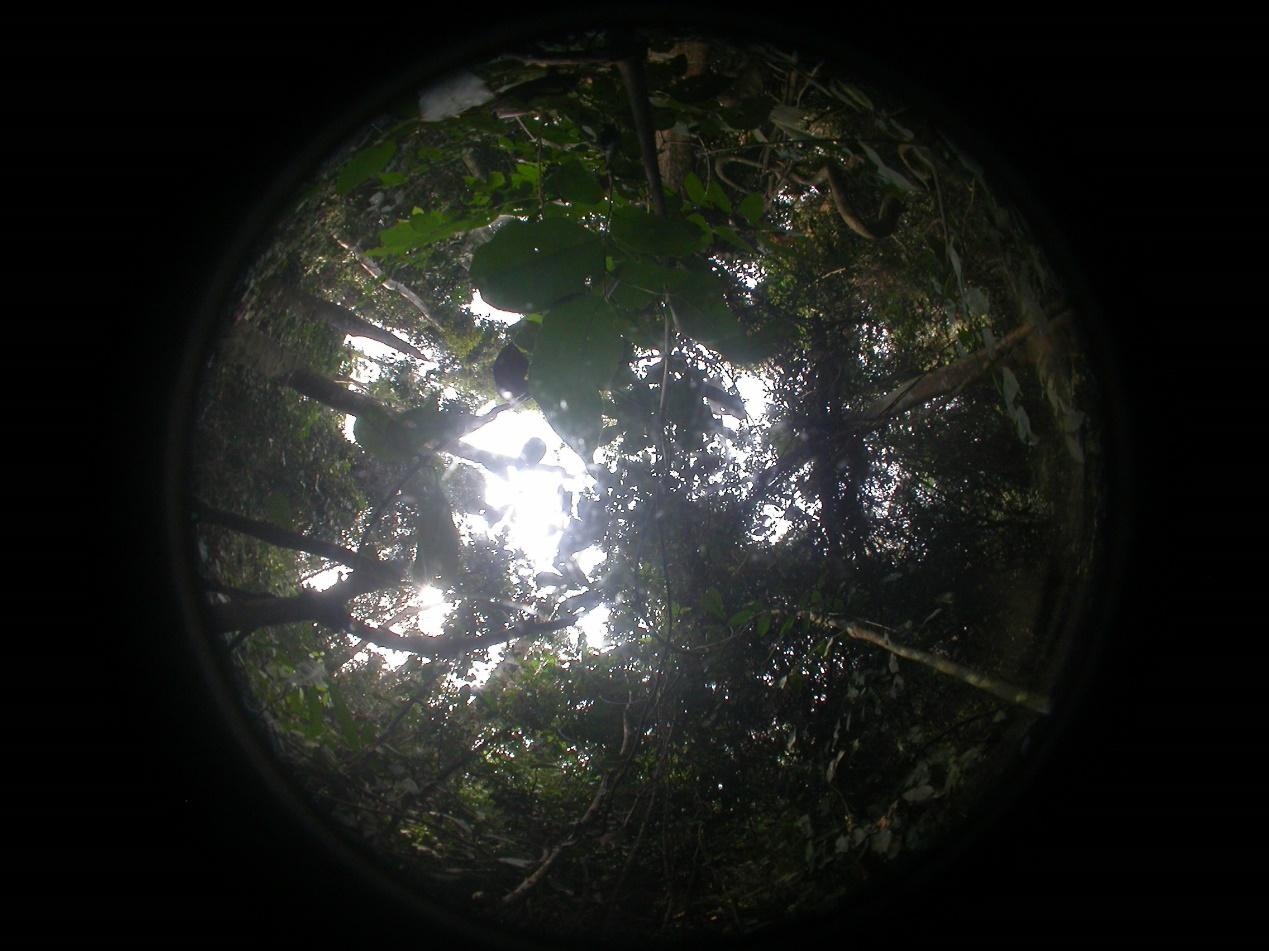

*Figure 2 Remember to bring a tree lopper, we don’t want big leaves that obscure most of the image.*

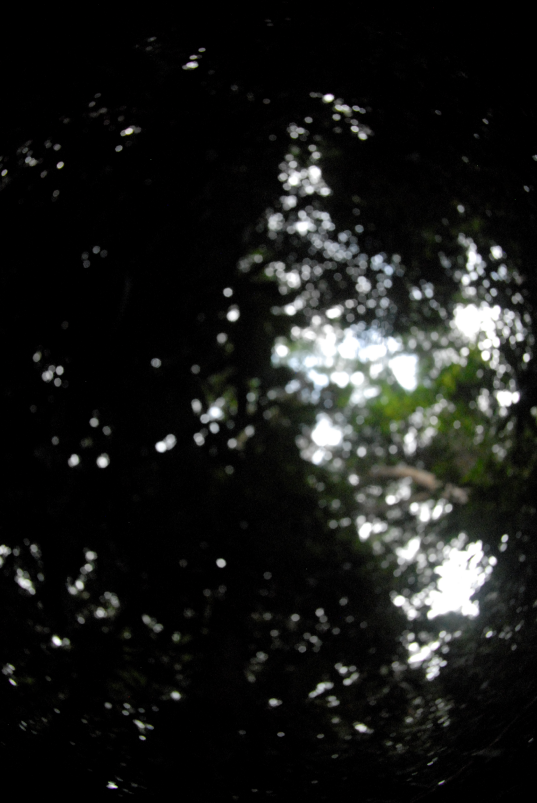

*Figure 3 Plot ANK-01, taken in July 2013. Using manual focus and a large aperture may help to get clear focus, but still in a dark tropical forest some photos may lose focus if not carefully checked in the field.*

Chen, J.M., Black, T.A., Adams, R.S., 1991. Evaluation of hemispherical photography for determining plant area index and geometry of a forest stand. Agric. For. Meteorol. 56, 129–143. https://doi.org/10.1016/0168-1923(91)90108-3

Jiao, S., Chen, Y., Cheng, Y., Duan, T., Gao, Z., Sun, Y., Huang, F., 2023. MEF-DHP: Digital Hemispheric Photography Method Based On Multi-Exposure Fusion, in: IGARSS 2023 - 2023 IEEE International Geoscience and Remote Sensing Symposium. Presented at the IGARSS 2023 - 2023 IEEE International Geoscience and Remote Sensing Symposium, pp. 2755–2758. https://doi.org/10.1109/IGARSS52108.2023.10282038

Xie, X., Yang, Y., Li, W., Liao, N., Pan, W., Su, H., 2023. Estimation of Leaf Area Index in a Typical Northern Tropical Secondary Monsoon Rainforest by Different Indirect Methods. Remote Sens. 15, 1621. https://doi.org/10.3390/rs15061621

Zhang, Y., Chen, J.M., Miller, J.R., 2005. Determining digital hemispherical photograph exposure for leaf area index estimation. Agric. For. Meteorol. 133, 166–181.

### Using ilastik to process raw fisheye images

Note that you may need to right-click ‘ilastik’ and select ‘Run as administrator’ on Windows computer (Otherwise there is an error at the Run feature selection stage). Ilastik may take up to 5 minutes to initialize. On Windows, you might need to press Enter when the first black screen.

First, create a new project using the Pixel Classification module (Figure 4).

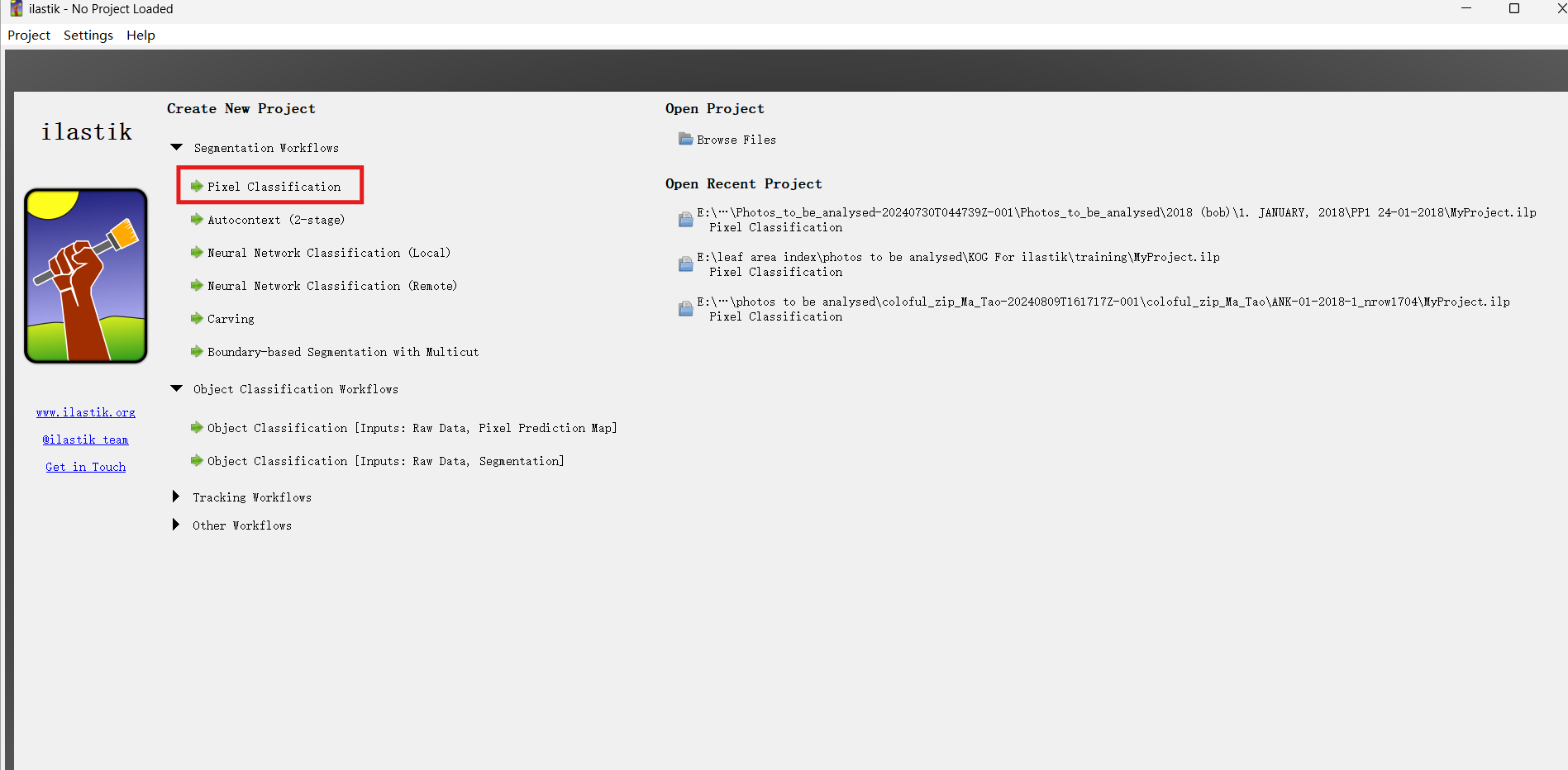

*Figure 4*

Next, choose a folder to store the project. Ensure the project is saved in the same folder as the training data before clicking save (Figure 5).

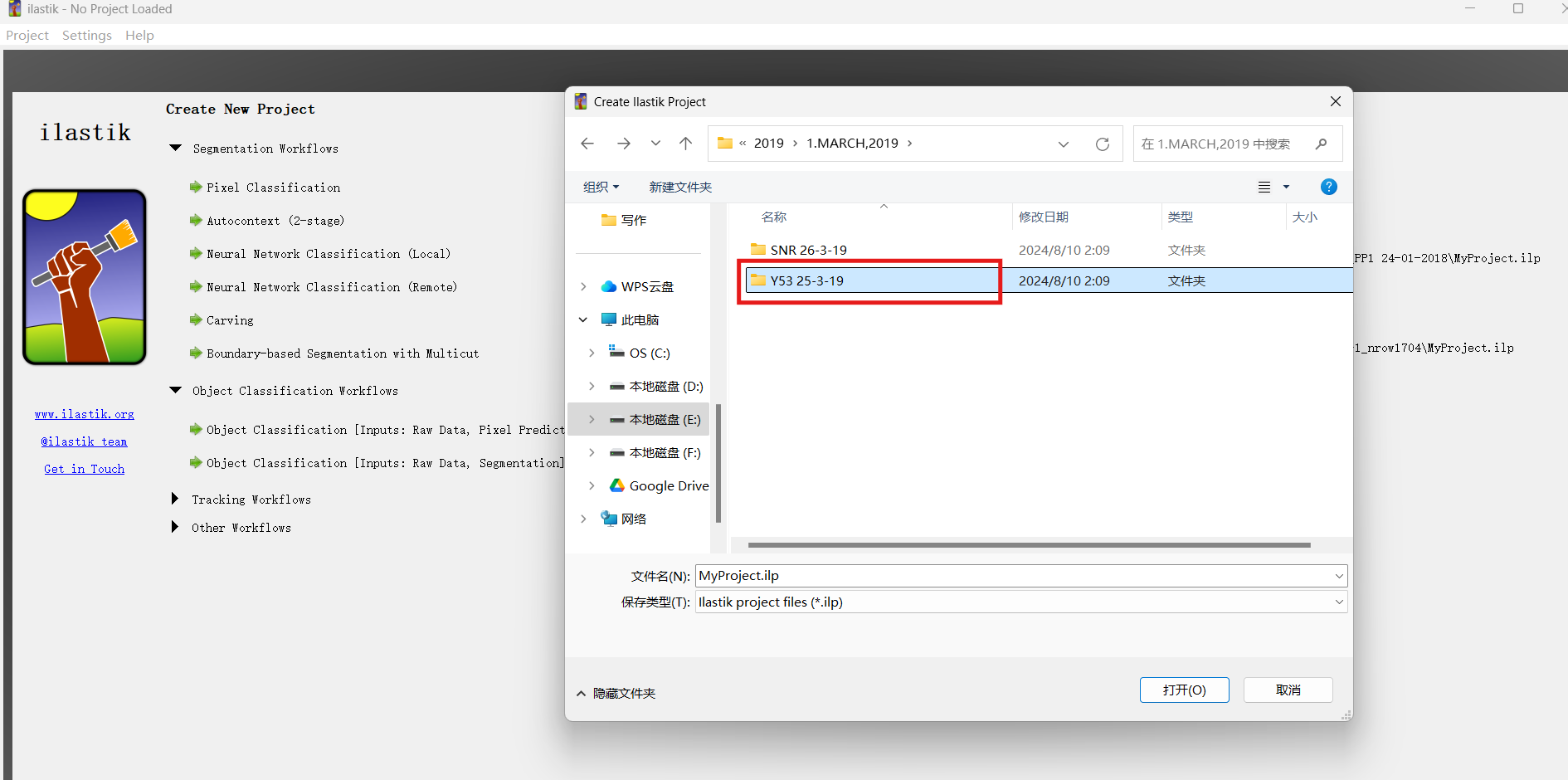

*Figure 5*

In the Input Data section, select images for analysis by clicking Add New, then click Add Separate Images (Figure 6). For example, the folder “Y53 25-3-19” would contain your training data (in this case, 25 images). I suggest naming it ‘training_data’ instead of Y53. It’s recommended to select contrasting photos from your photo pool and copy them into this folder. Typically, I include photos from various sites, seasons and weather conditions to ensure they represent the full range of data. A good starting point is 10 photos. Note: If you plan to share your ilastik model, you must send the entire “Y53 25-3-19” folder, not just the .ilp file. After saving, you’ll notice a new .lip file in this folder (this file stores your manual annotations, as explained later).

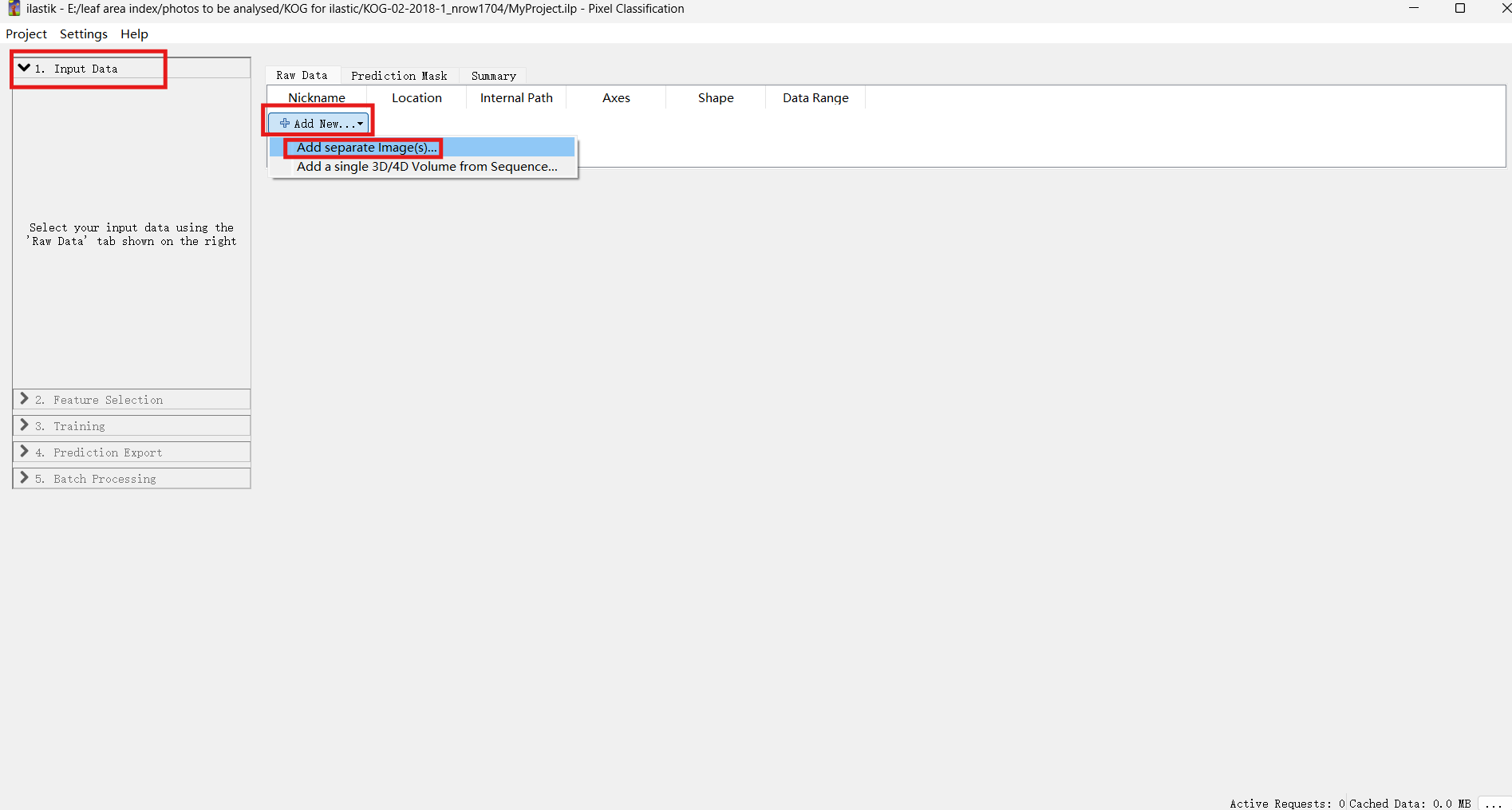

*Figure 6*

Next, go to the folder that contains all the training data. You can either drag the files or press Ctrl+A to select all photos at once (Figure 7).

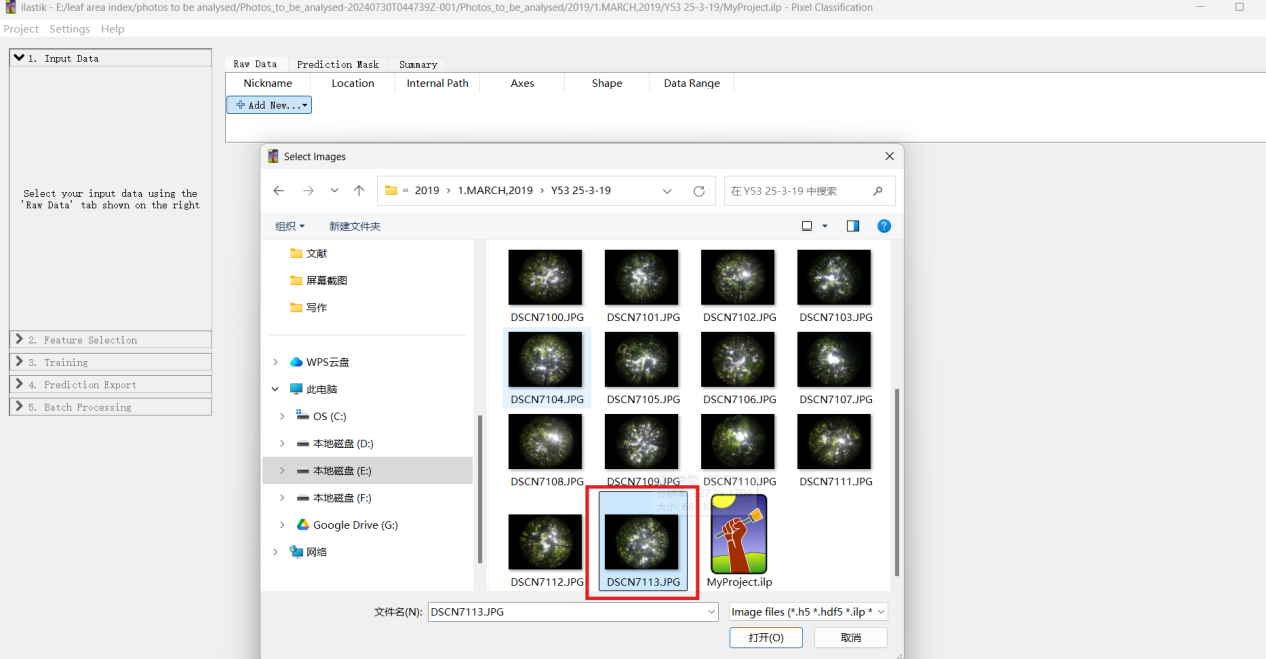

*Figure 7*

Alternatively, you may add only one photo now and come back later to add more if additional training photos are needed (Figure 8 and Figure 9). Naturally, you can delete those any photos that aren’t used for training (after completing the protocol).

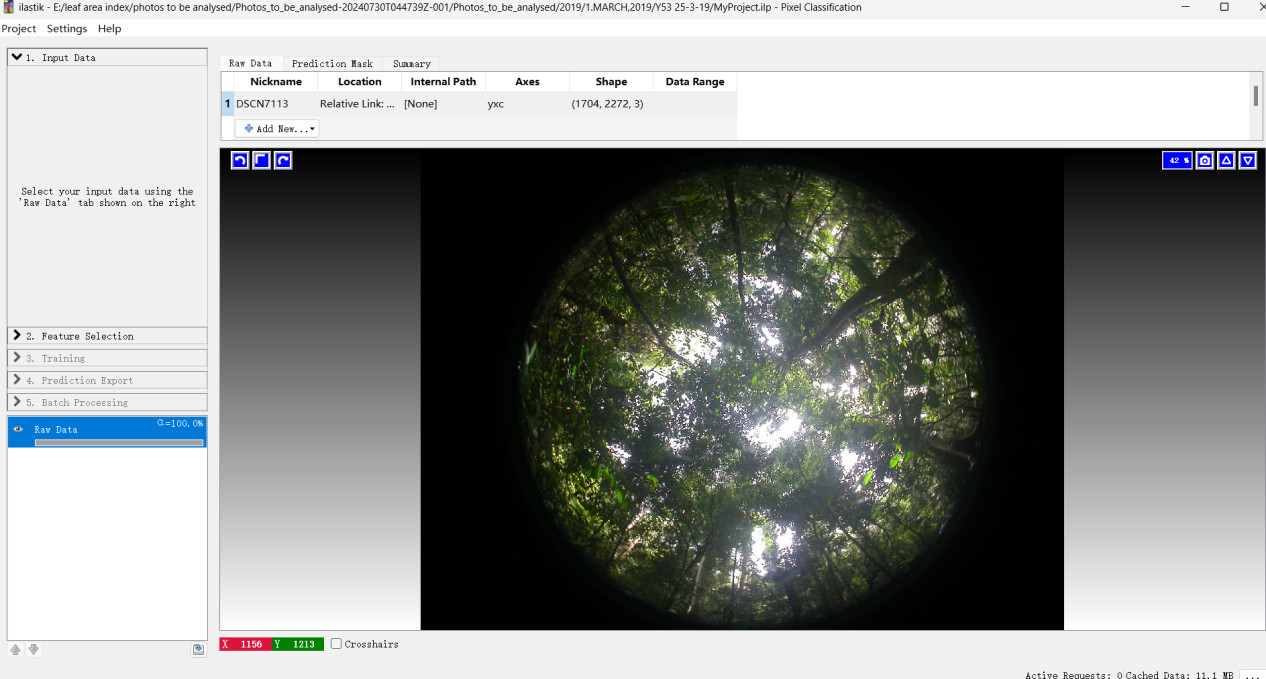

*Figure 8*

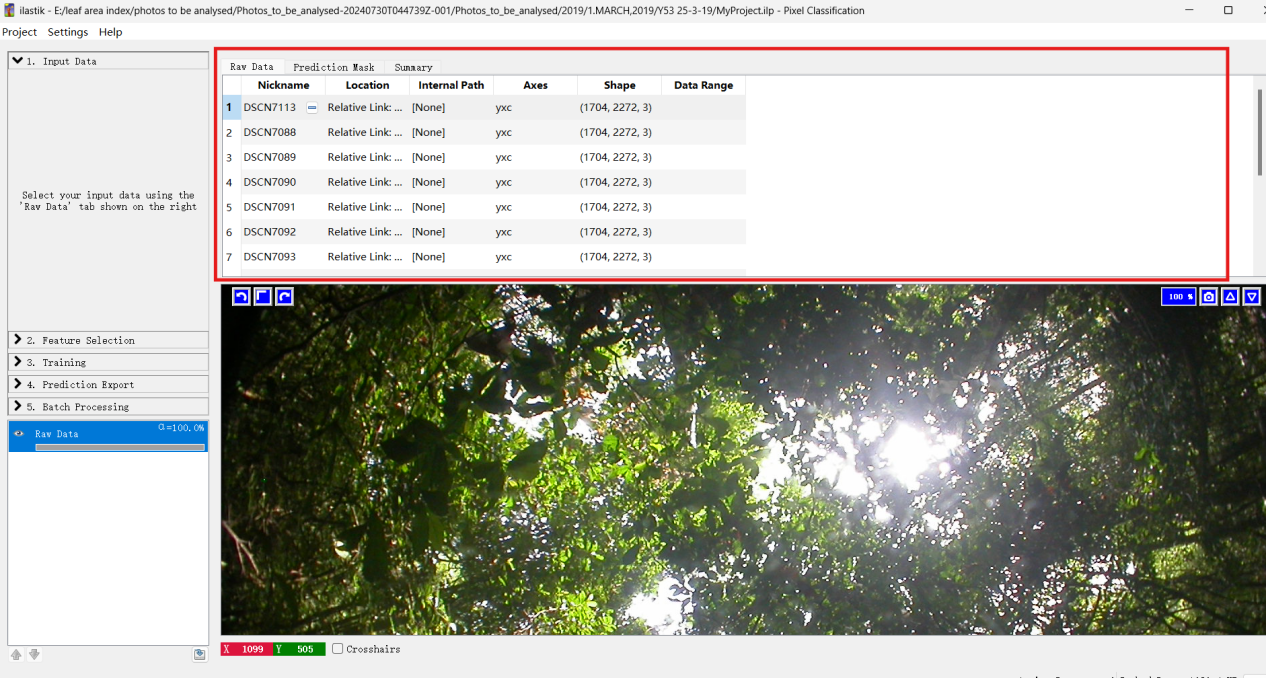

*Figure 9*

Next, we will go to feature selection. Click “Select Features” to proceed (Figure 10).

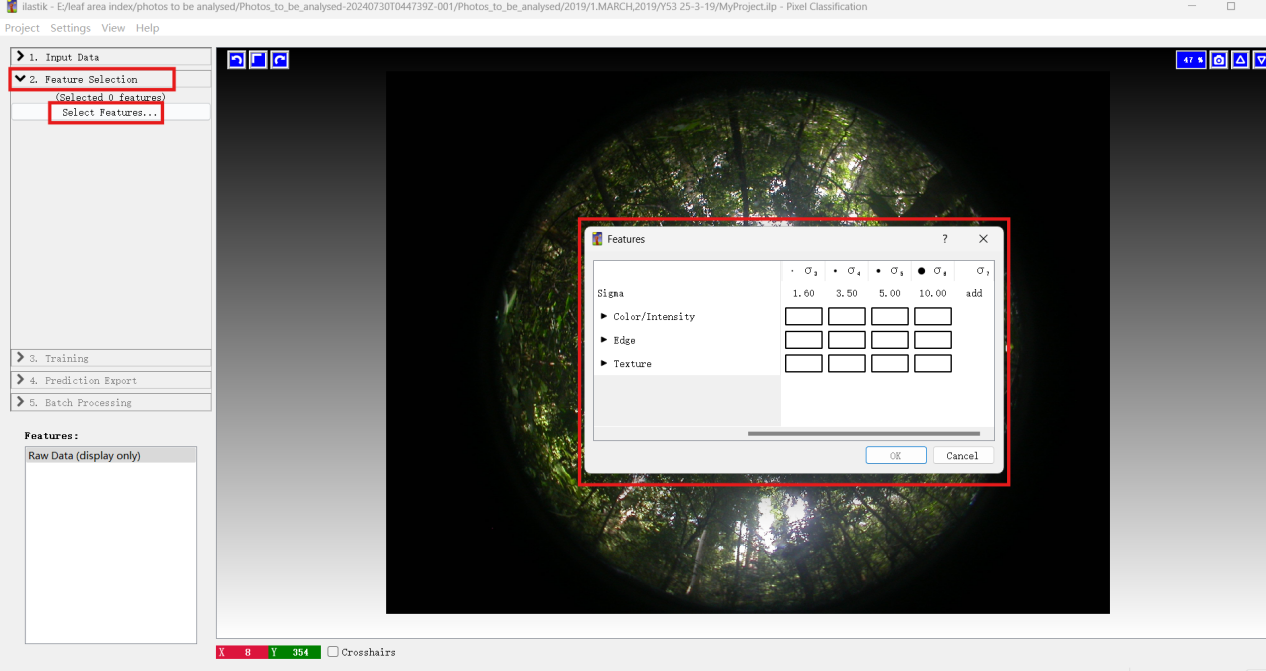

*Figure 10*

The ‘Features’ window in ilastik is dedicated to feature selection (Figure 11). These parameters allow users to configure various feature filters, enabling the classification algorithm to more effectively recognize and process image characteristics such as textures, edges, colors, and other relevant information.

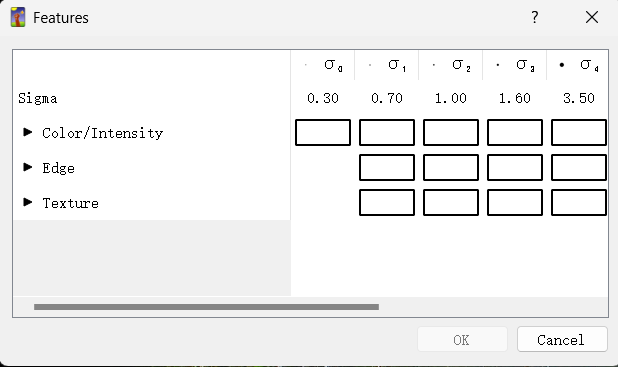

*Figure 11*

The following functions are available:

(1) Color/Intensity: Color is one of the most distinguishing features for differing between the canopy and the sky. Because of leaves, branches, and shadows, the canopy typically appears in shades of green, brown, or darker tones, while during sunrise or sunset, the sky often appears in lighter colors, such as shades of blue, white, or even orange and pink.

(2) Edge: Edge features detect boundaries characterized by significant changes in pixel intensity or color.

- Canopy-Sky Boundary: The irregular outlines of leaves and branches against the sky create distinct edges.
- Complexity: The natural shapes of the canopy introduce complex edges that enhance recognized.

(3) Texture: Texture captures the surface patterns and variations within image regions. Owing to the arrangement of leaves, branches, and varying shadows, the canopy exhibits a complex texture, while the sky typically has a smooth texture, though clouds can introduce some patterns.

Each feature can be applied at various scales, with the scale corresponding to the sigma of the Gaussian function used to blur the image before applying the filter. A higher sigma value allows the filter to capture information from a wider area while simultaneously reducing finer details. The following image provides an example of an edge filter computed using three different sigma values. For example, filters with lower sigma values detect small edges more effectively, while those with higher sigma values identify only more general or broader structures.

| 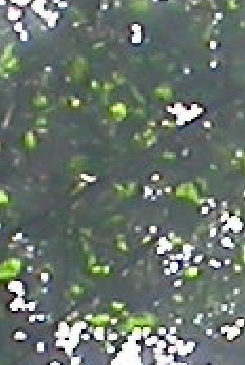 | 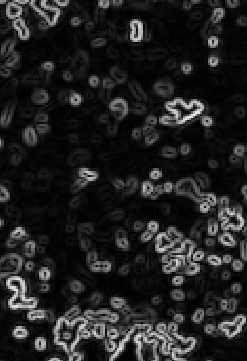 | 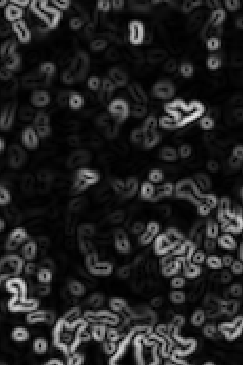 | 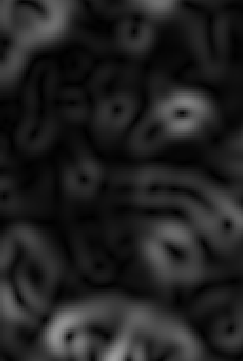 |
| --- | --- | --- | --- |
| Raw data | Sigma=0.7 | Sigma=1 | Sigma=1 |

To enable specific image features (e.g., color, edge, texture) and assign their corresponding Sigma values, activate the respective checkbox (green check mark). The selected features are incorporated into the machine learning model and utilized to enhance class discrimination during both the training and classification phases. For this analysis, I will enable the following features (Figure 12). While additional selections are possible, they may increase computational processing time.

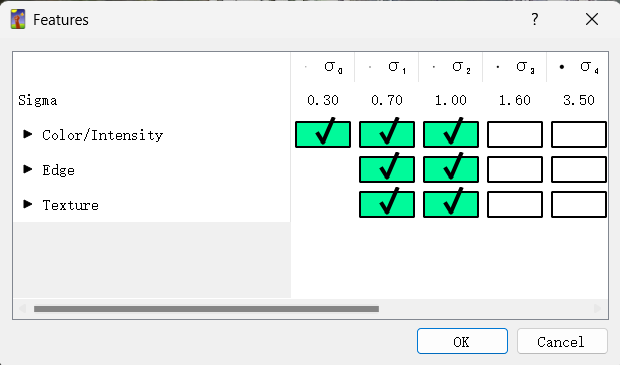

*Figure 12*

The complete set of selected features is displayed in this panel (Figure 13). Here, users can interact with individual features by clicking on them to assess their discriminatory power—specially, how effectively each feature differentiates the target objects (leaves) from background elements (sky). This visualization step enables qualitative validation of feature selection prior to model training.

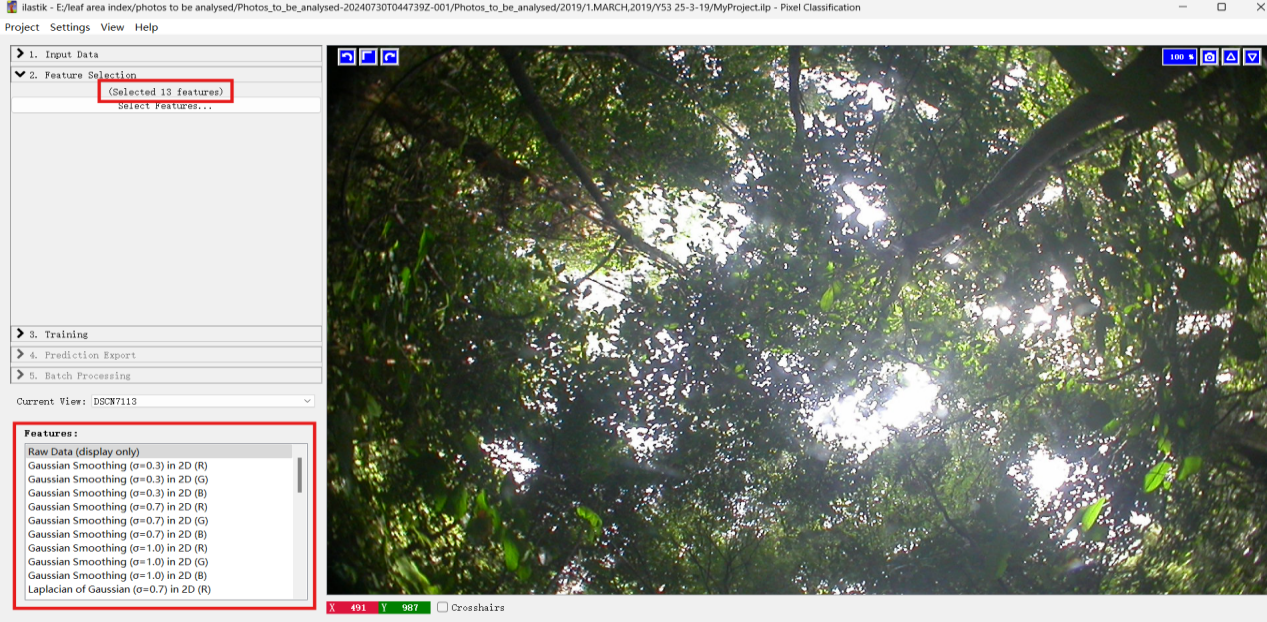

*Figure 13*

The subsequent phase of pixel classification entails training discriminative model to differentiate between distinct object categories. This iterative workflow requires the user to: (1) provide initial annotations for representative regions, (2) evaluate the model’s predictive output, and (3) iteratively refine the annotations to correct misclassified pixels and improve the model accuracy. Each iteration enhances thew model’s ability to generalize from the annotated examples.

We now proceed to the training phase within ilastik, which requires manual annotation of image regions to differentiate between sky and leaf categories (Figure 14). Label customization is achieved double-clicking the color swatch adjacent to each class designation where the default configuration assigns blue (first label) to sky regions and green (second label) to leaves. Critical implementation note: The ordinal position of labels directly determines the output values in the processed images, where the first category is encoded as 1^st^ (sky) and 2^nd^ (leaves). Additionally, enabling renormalization option [min, max] during subsequent processing stages may introduce alternate pixel values (0 or 100) in the exported results.

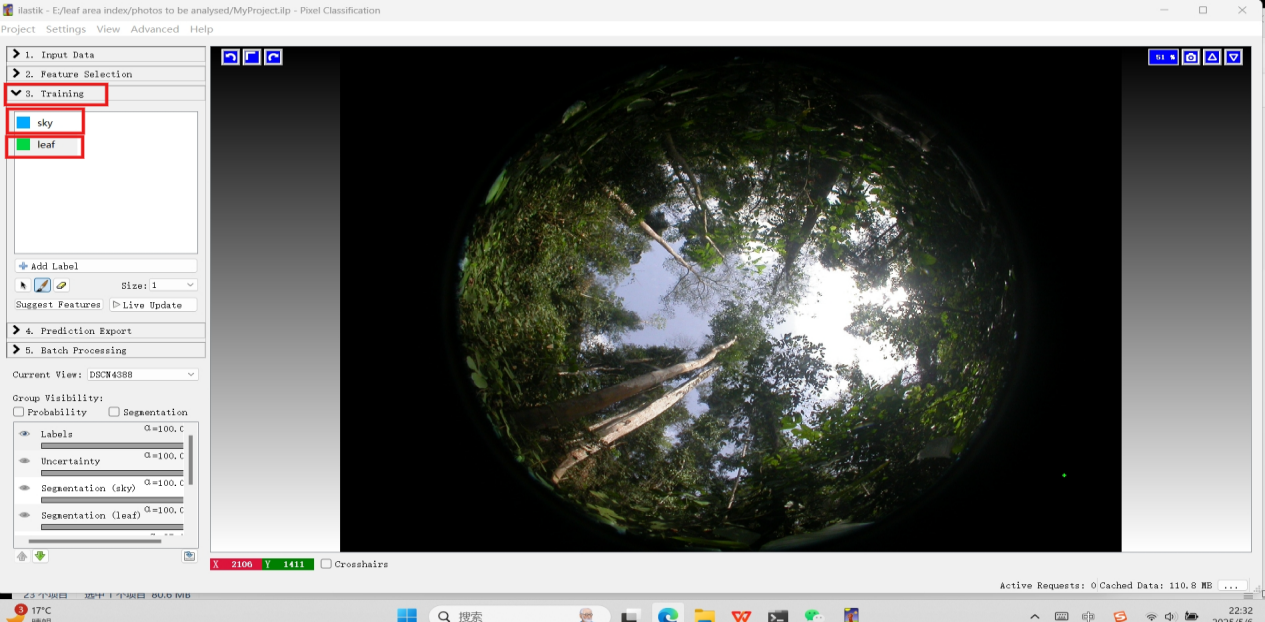

*Figure 14*

Here, we use two labels—blue for sky and green for leaves (Figure 15). Note that CAN-EYE prefers sky as the first category, so we recommend assigning as Label 1 to ensure consistency with standard processing.

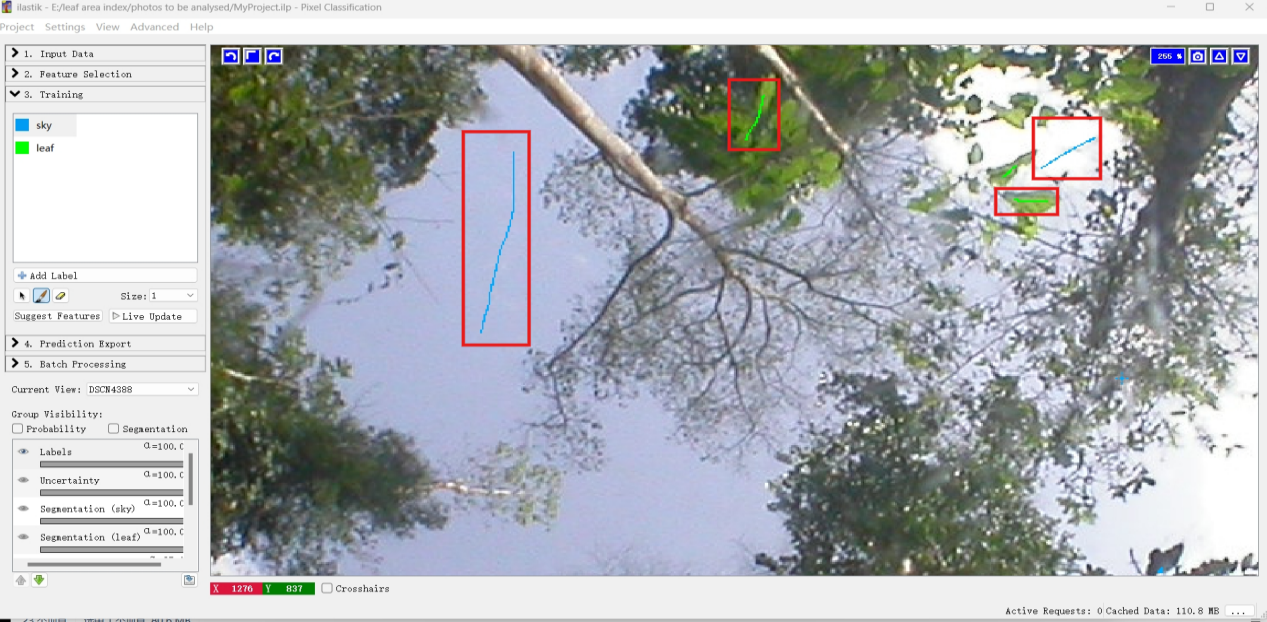

*Figure 15*

For particularly complex forest canopies, such as those with dense branching structures, a ternary classification approach may be applied. In this method, branches are labeled in red to explicitly exclude them from leaf classification (Figure 16).

Additionally, for "grey areas" where human judgement is uncertain (i.e., when distinguishing between branches and leaves is unclear), these areas should be left unmarked. The algorithm will automatically process these zones. Only confidently identifiable features should be annotated.

To adjust the view, hold the Ctrl key while scrolling to zoom, and use the middle mouse button to pan the image. Note that this function requires either a mouse or touchscreen device; touchpad controls are not supported.

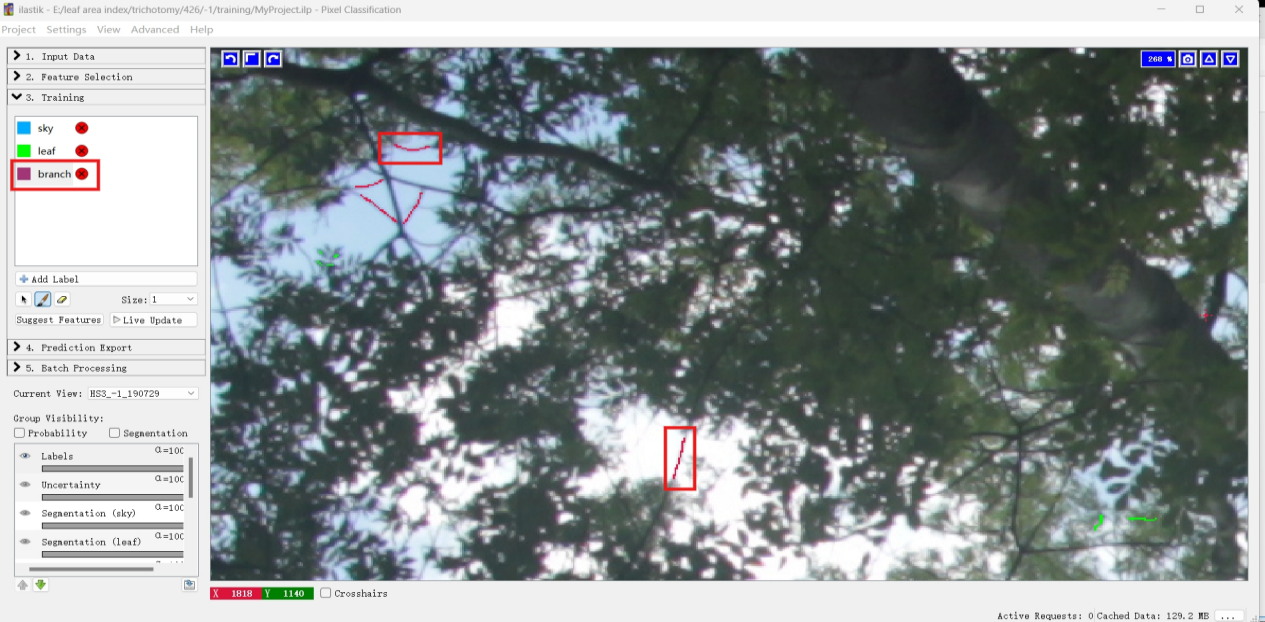

*Figure 16*

Click Live Update to observe ilastik’s classification results for your image (Figure 17). Toggle between probability view and segmentation view to examine the analysis output. If required, you may add manual annotations to improve results, though this should only be done when Live Update is disabled to prevent system instability. Important Note: Always ensure Live Update is turned off before annotating, as concurrent operations may cause software crashes, and save your project regularly to prevent data loss.

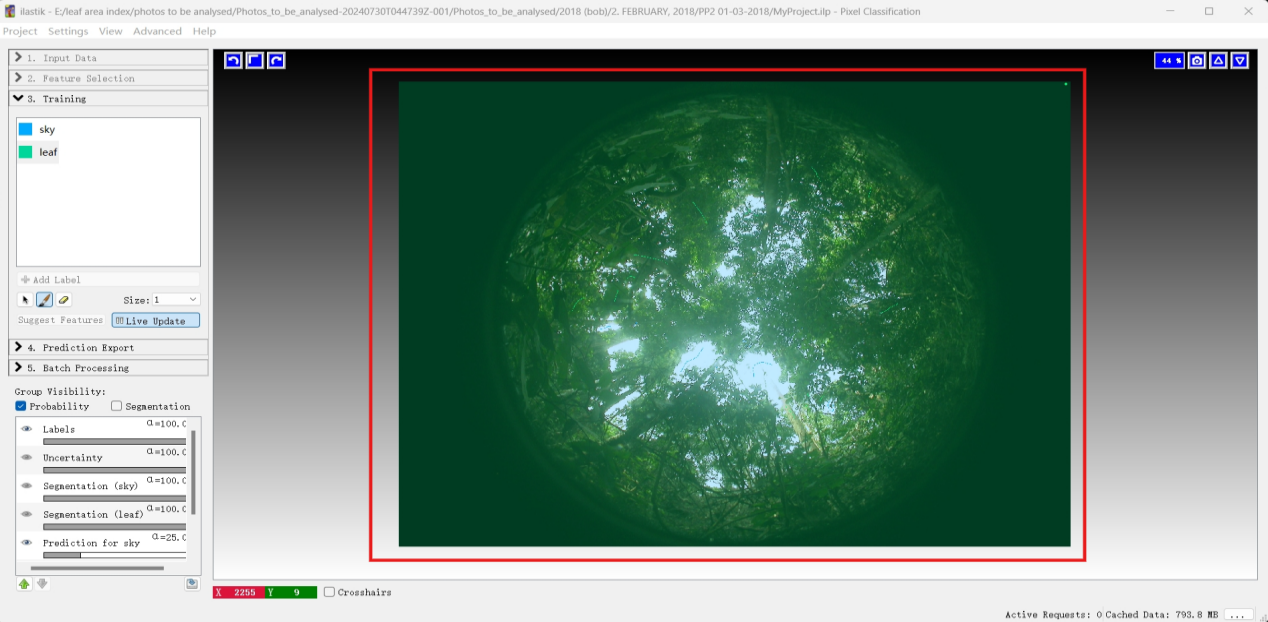

*Figure 17*

To view the final classification results, enable the Segmentation overlay by activating Segmentation checkbox in the interface (Figure 18).

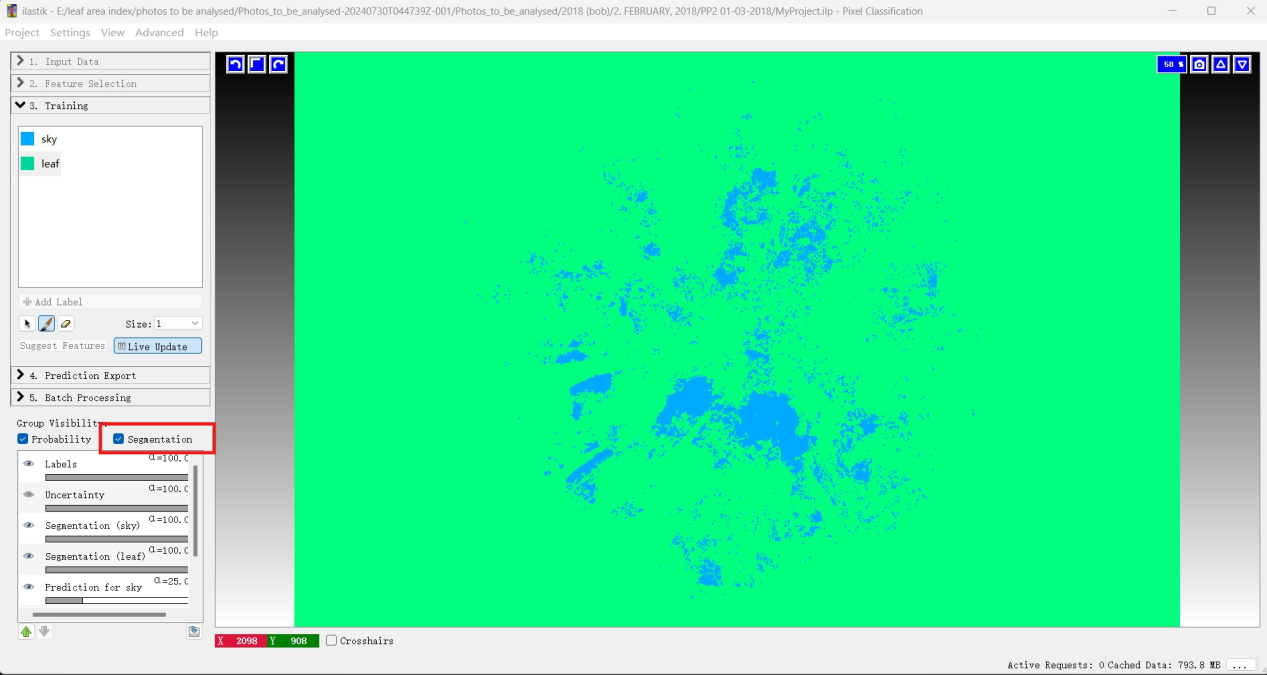

*Figure 18*

If the segmentation appears incomplete or inaccurate, pause Live Update and refine the annotations by adding more marks. Note that you can use Ctrl+Z (undo) or the eraser tool to correct any mistakes. Once you finish annotating the current image, click “Current View” to proceed to the next training image (additional images can be imported in the 1. Input Data step if needed). Re-enable Live Update—the software will now segment the new image using the knowledge gained from previous annotations. If the results are unsatisfactory, add more marks to improve segmentation. If the output meets expectations, move on to the next image. Repeat this iterative process until the model consistently produces satisfactory results without requiring additional manual corrections on new images.

If the current results meet your quality standards , disable Live Update by clicking it again, then select Suggest Features to open a new configuration dialog box (Figure 19).

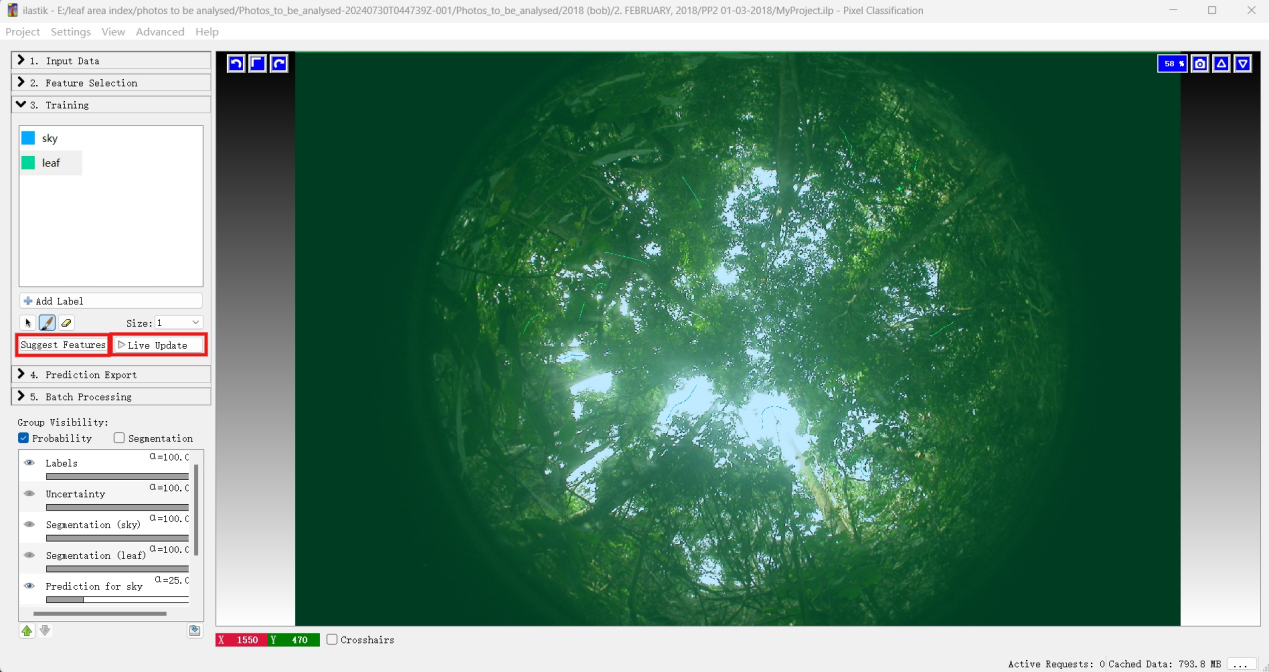

*Figure 19*

If you are uncertain about your pixel feature selections, the Suggest Features functionality can assist in optimizing them. This feature evaluates the classifier’s performance on annotated pixels by testing different feature combinations. The process employs machine learning technique, including random forest classification and "out-of-bag" error estimation, to assess prediction accuracy. Additionally, ilastik provides computation time metrics, allowing users to balance runtime efficiency with segmentation quality. For users without a computer science background, the system automatically refines selection to improve both prediction accuracy and processing speed, reducing the need for manual tuning.

Next, you will reach the “Feature Selection” page. For users with machine learning expertise, this section provides detailed explanations of key functionalities and adjustable parameters, allowing you customized the process based on your requirements. For those without a technical background, you may safely ignore these advanced options and proceed by selecting the “Filter Method (recommended)” option. “Run Feature Selection” to continue (Figure 20).

Note: You will need to run ilastik as Administrator on Windows if you get an error

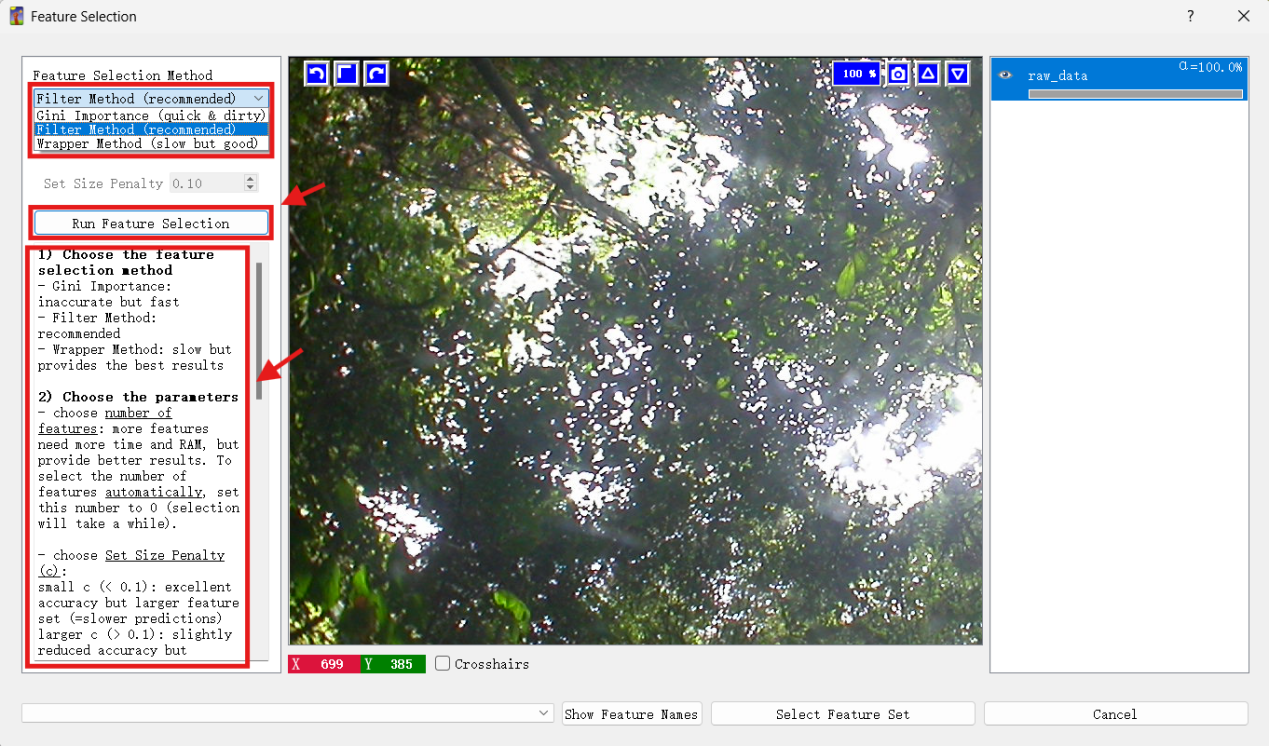

*Figure 20*

Select "Filter Method (recommended)" and then click “Run Feature Selection”. Choose then appropriate feature selection and click “Select Feature Set” (Figure 21).

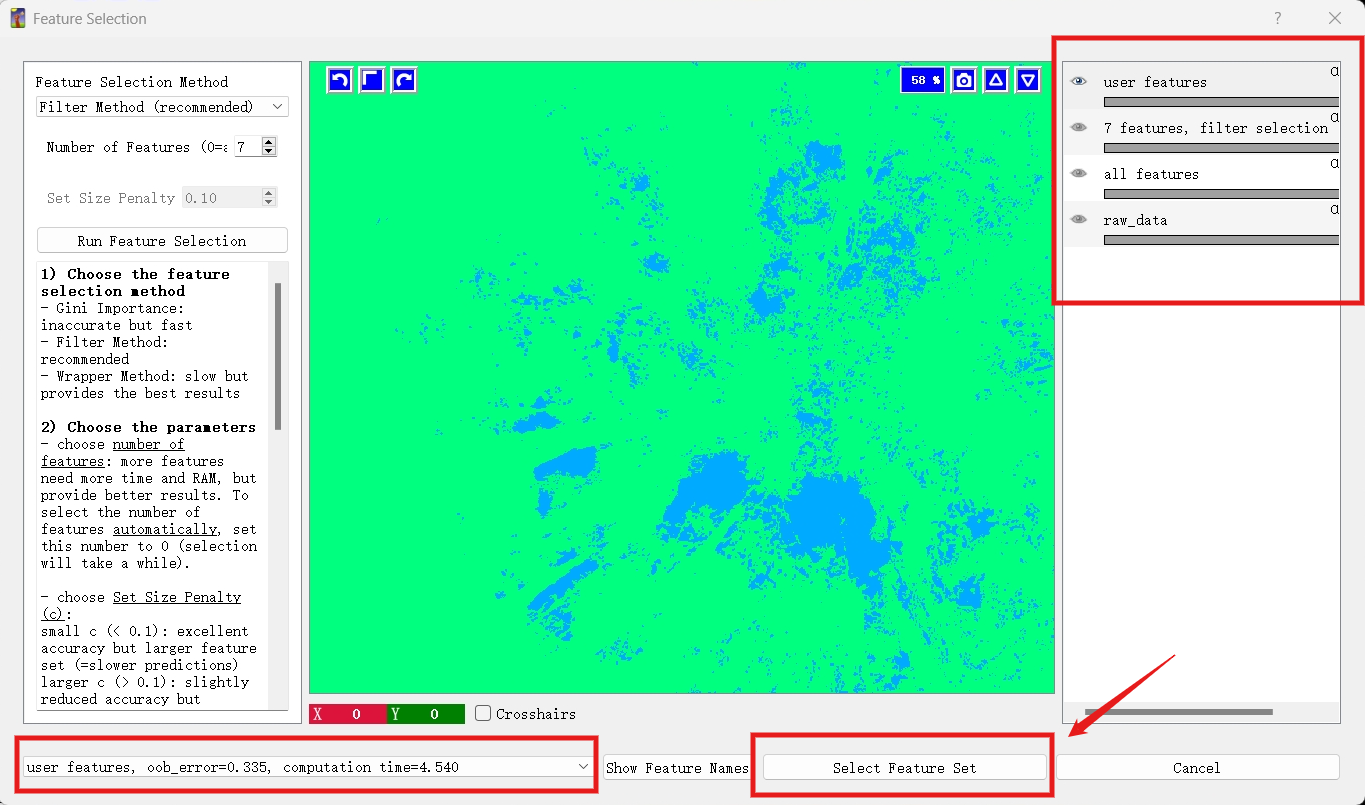

*Figure 21*

In the displayed ilastik feature selection interface, the three options highlighted in the red box at the bottom left are located in the drop-down menu below. Each option represents different feature selection methodologies and usage scenarios. Their key differences are as follows:

(1) User features: This denotes the feature set manually selected by the user during configuration.
(2) 7 features, Filter selection: This option indicates that 7 features were automatically selected using the Filter Method(recommended).
(3) all features: This option includes all available features without reduction (i.e., no filtering was applied), comprising the full set of initial features.

Notably, all three options provide two metrics:

- Oob Error (Out-of-bag Error): Quantifies the model's error rate when evaluated on unseen data, serving as a robust estimate of generalization performance.
- Computation Time: Represents the duration required for the model to process the corresponding feature set.

In an optimal scenario, both the oob_error and computation time should be minimize. However, practical implementation often requires trade-offs between these metrics. When computational resources are constrained or rapid results are needed, selecting fewer features may be optimal. Conversely, if model accuracy is the primary objective, increasing the feature count—despite longer computation times—may be justified.

For this demonstration, we select “User Features” as the most appropriate choice for our workflow. After selection, click “Select feature Set” to exit the Feature Selection interface, followed by “Live Update” to view the results. It is important to note that the “Suggest Feature” step can be optionally bypassed as it is particularly computationally demanding. This step may cause performance issues or system instability when processing extensive training datasets, numerous pre-selected features, or when running on hardware with limited resources such as standard laptops. In such cases, we recommend first reducing the number of pre-selected features, limiting the quantity of training images and annotations, then running the “Suggest Feature” process once to identify the optimal feature set for future use. If these adjustments still prove problematic, users can confidently proceed with the predefined recommended features and skip this step entirely, as it is not strictly necessary for successful implementation.

When we click “Select feature Set”, the Feature Selection screen closes. Then, click “Live Update” to view the results (Figure 22).

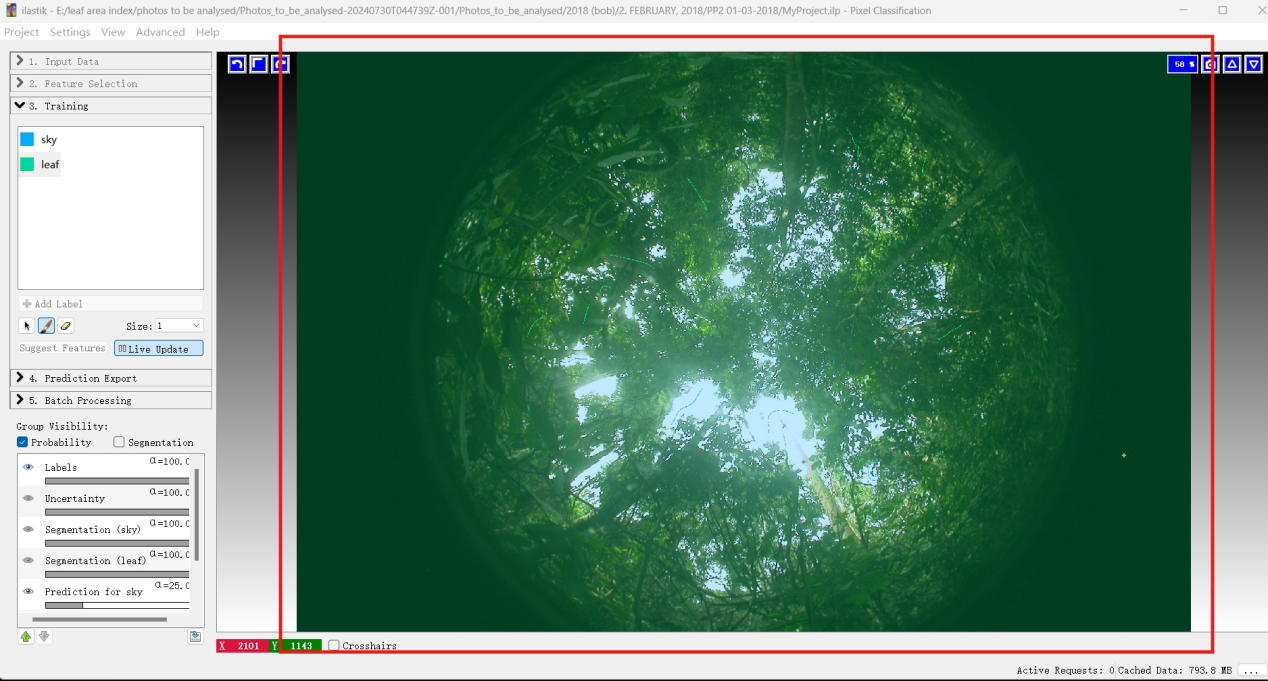

*Figure 22*

(NoteL: If you are using CAN-EYE, set “sky” as the 1^st^ category.)

Stop the “Live Update” and navigate to 4, the “Prediction Export” section (Figure 23). Now that you're in “Prediction Exports”, locate the “Export Settings” panel and under “Source”, change the selection “Probabilities” to “Simple Segmentation”.

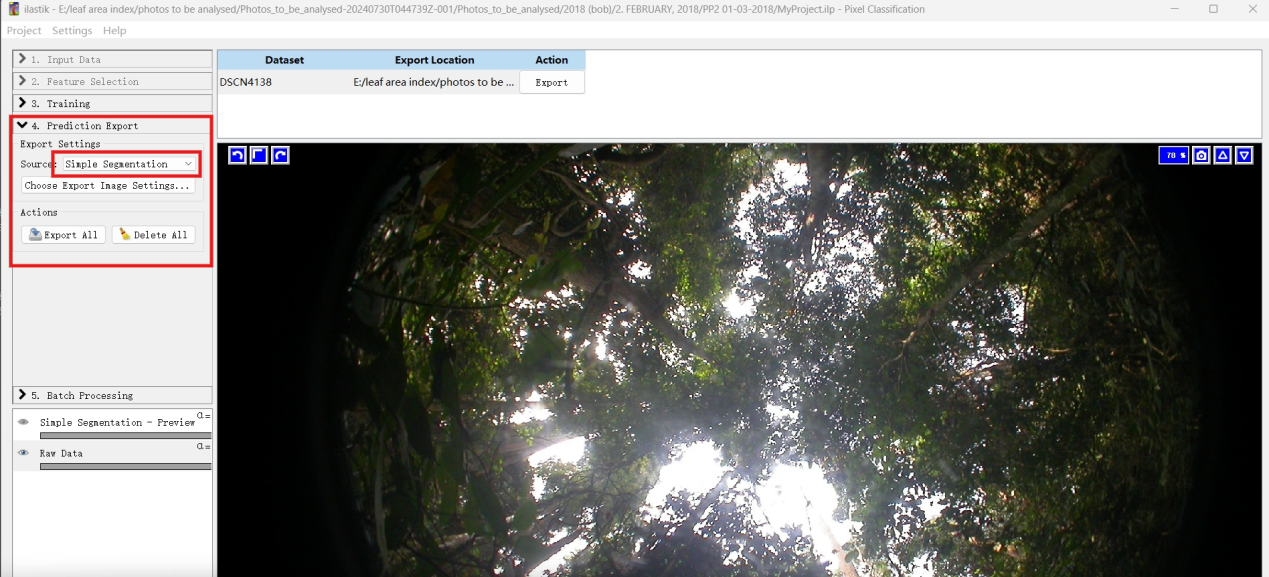

*Figure 23*

(Change Probabilities to Simple Segmentation)

First, Click ‘Choose Explore Image Settings' and under the 'Transformation', select 8-bit integer. Then enable ‘Renormalize’ and change the value from '1 to 2' to '0 to 100' (Figure 24). Note that in the training phase, label 1 represents the sky while label 2 represents the leaf. If you don’t enable Renormalize, ilastik will save output images containing values 1 and 2. When you enable Renormalize, all pixels in your 1^st^ category (label 1, which should be sky for CAN-EYE) will be converted to 0, while the 2^nd^ category (label 2) will become 100. For ternary classification (when you have categories 1, 2, and 3), enabling Renormalize with '1 to 2' to '0 to 100' setting will convert category 1 to 0, category 2 to 100, and category 3 will become 100. In such cases, it’s recommended to leave ‘Renormalize’ disabled, and instead use Python, R or Matlab to manually convert the output values (1, 2, 3) to your preferred numbers. To summarize the protocol for CAN-EYE: make sure your category 1 is assigned to sky, then enable Renormalize and set the conversion from '1 to 2' to '0 to 100'.

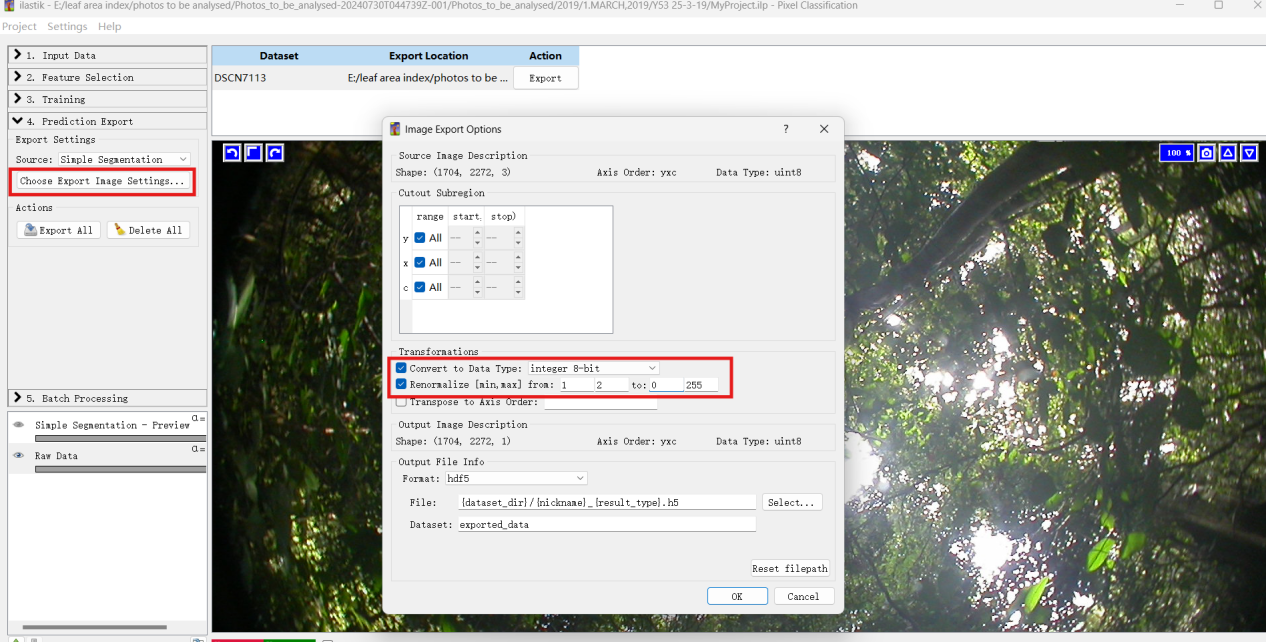

*Figure 24*

Under ‘Output File Info’, change the data type from hdf5 to jpeg, as this format easier to view. Then click OK (Figure 25).

Under files, the interface shows where your outputs will be saved. You can change this to another folder if needed, but remembering that the naming convention {nickname}_{result_type} is critical. For example, my output path is: C:/Outputs/{nickname}_{result_type}.jpg

Important: Always use forward slashes (////) in the path, not backslahes (\\\\).

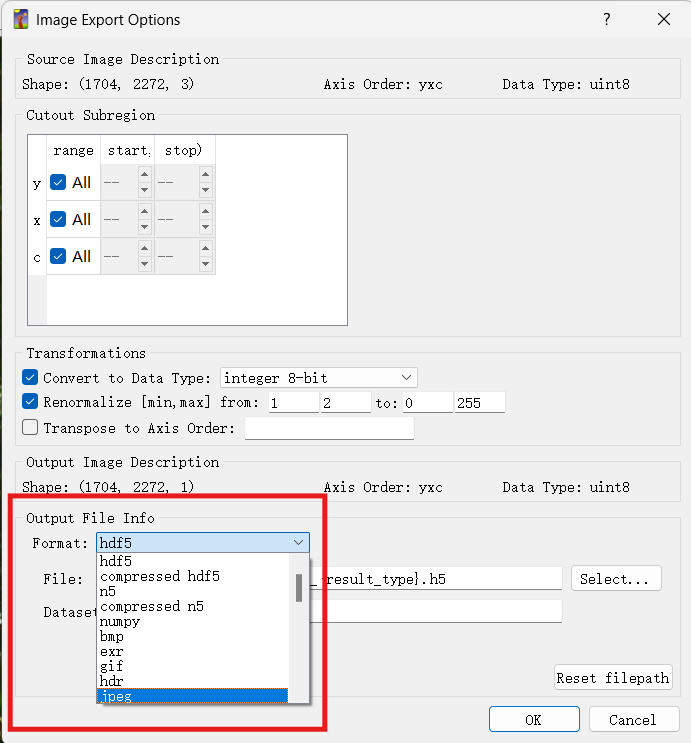

*Figure 25*

Then use ‘Export’ or ‘Export All’ to convert all your training photos to black and white (Figure 26).

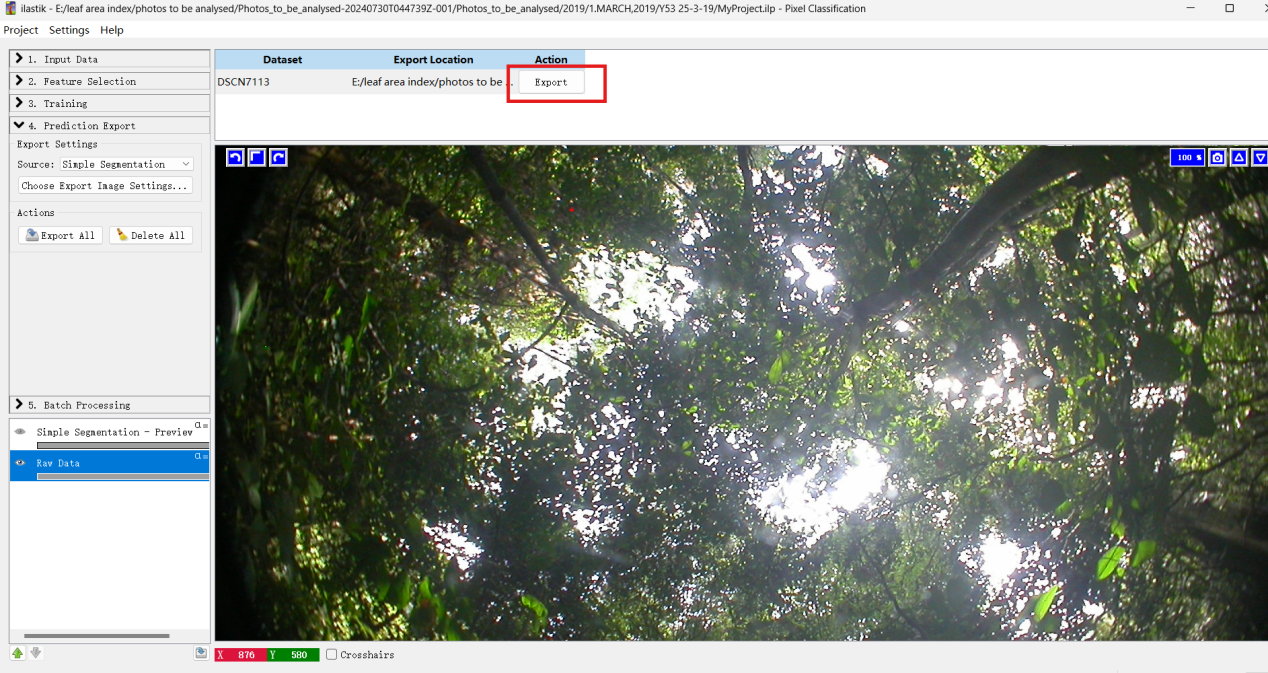

*Figure 26*

Don’t forget to save your project regularly! Otherwise, you may encounter errors when you next open the ilastik file (Figure 27).

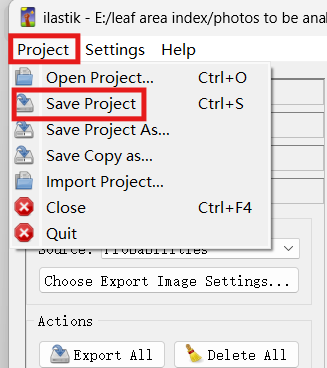

*Figure 27*

Now, you will find the black and grey image in your output folder (Figure 28).

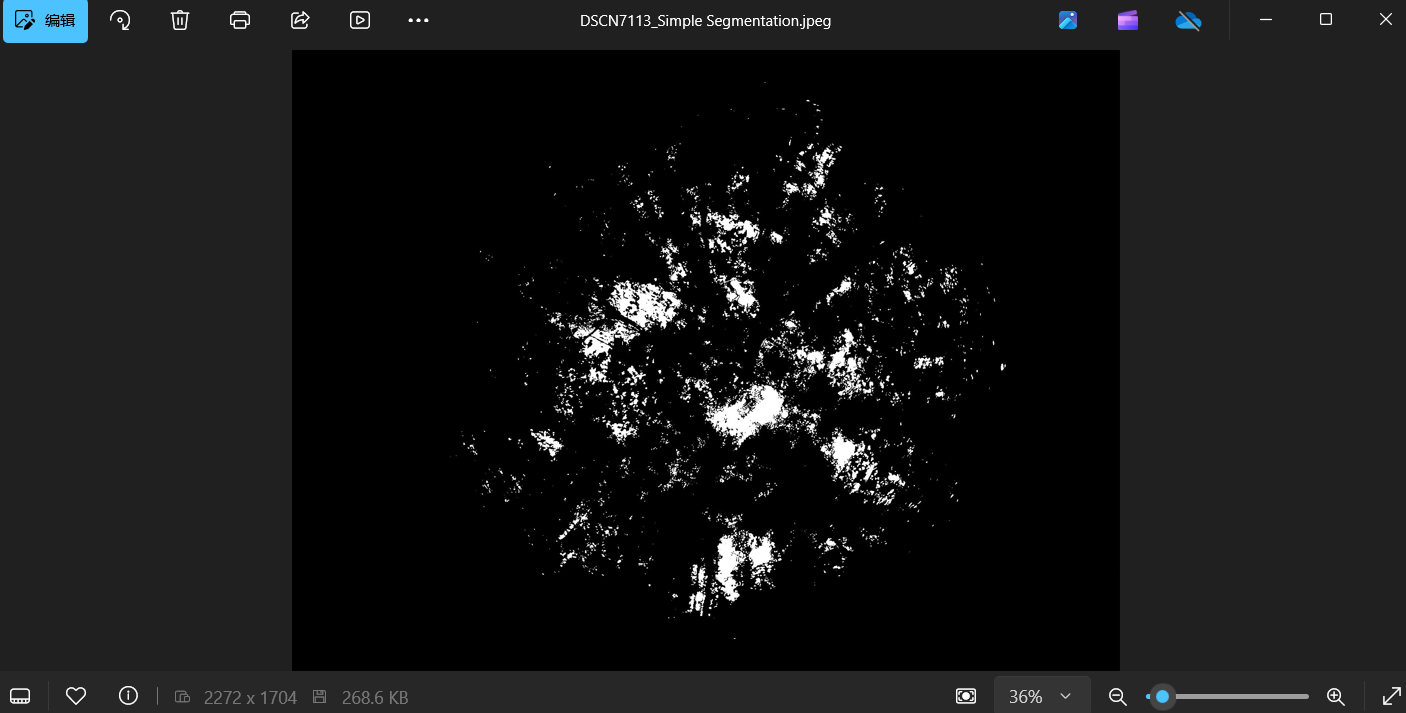

*Figure 28*

Because this is a JPEG, the values of 0 and 100 would be understood as greyscale – where 0 is black and 100 is grey. Thus, if you uncheck ‘Renormalize’, meaning you have values of 1 and 2, the image **will appear totally black**! To view such an image, I normally use QGIS software.

To get better results, you should include more images in your input data. Here I have 20 images, which is approximately the minimum number needed to get relatively good results when processing many images.

Now, you can use Batch Processing to process all your images (other than those used for training)(Figure 29).

Before batch processing, an **EXTREMELY** important step is to name your images consistently so that you can identify them later.

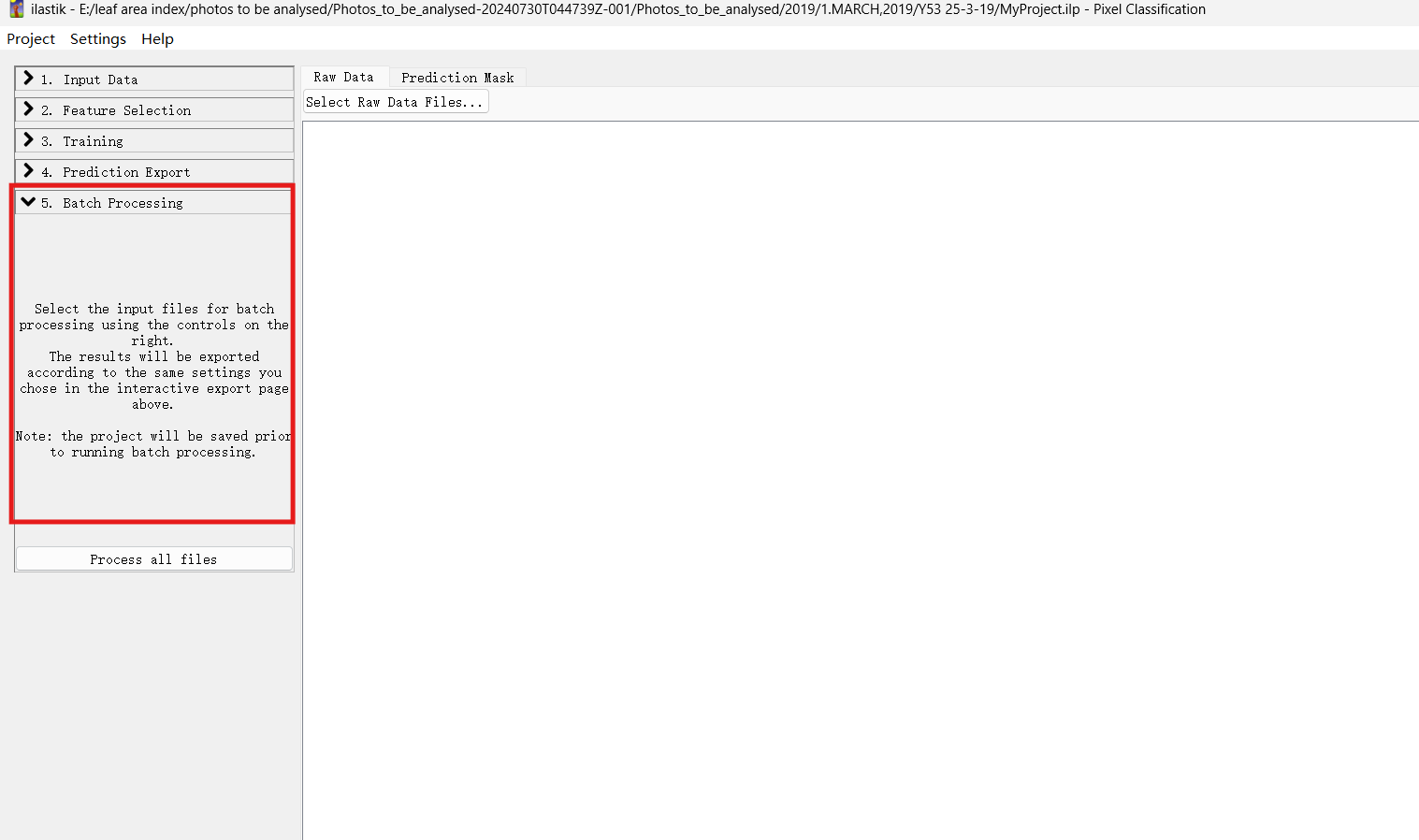

*Figure 29*

Click ‘Select Raw Data Files’ and find the folder containing all images you want to analyze (Figure 30). Select all of them and click ‘Process All Files’. Use CTRL+A to select all files quickly (Figure 31).

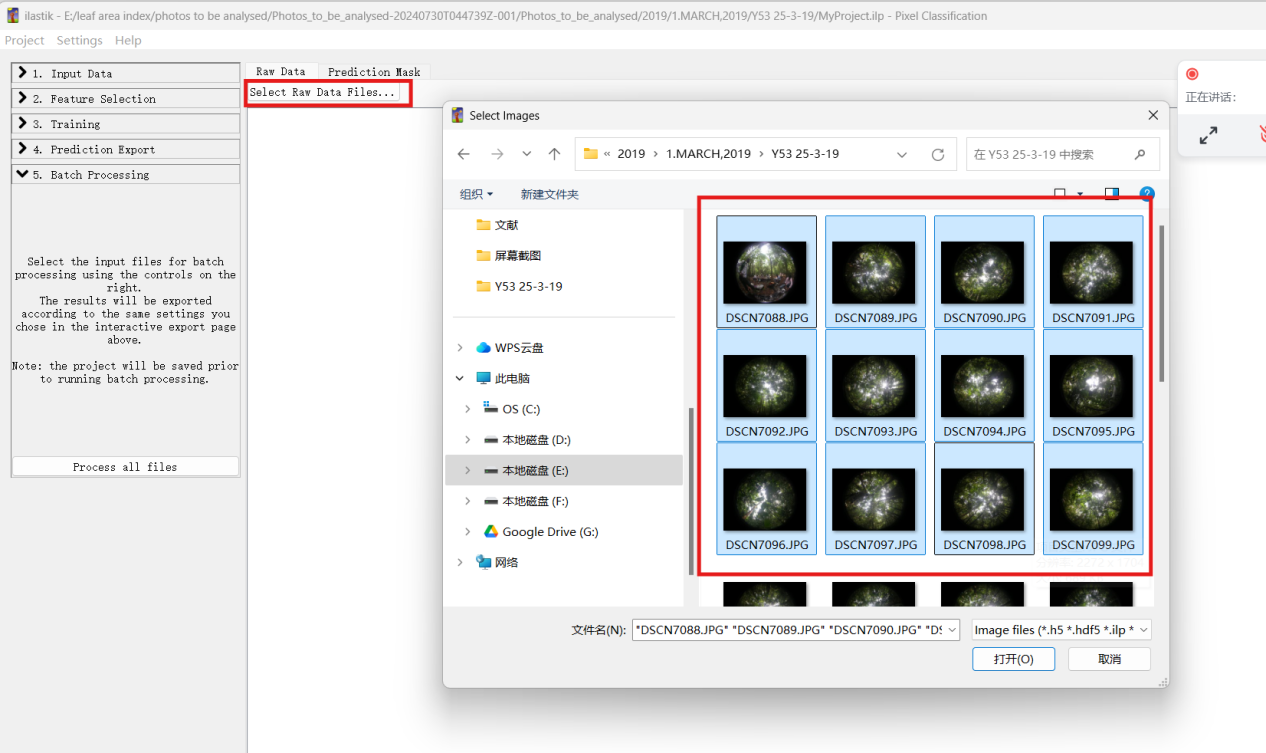

*Figure 30*

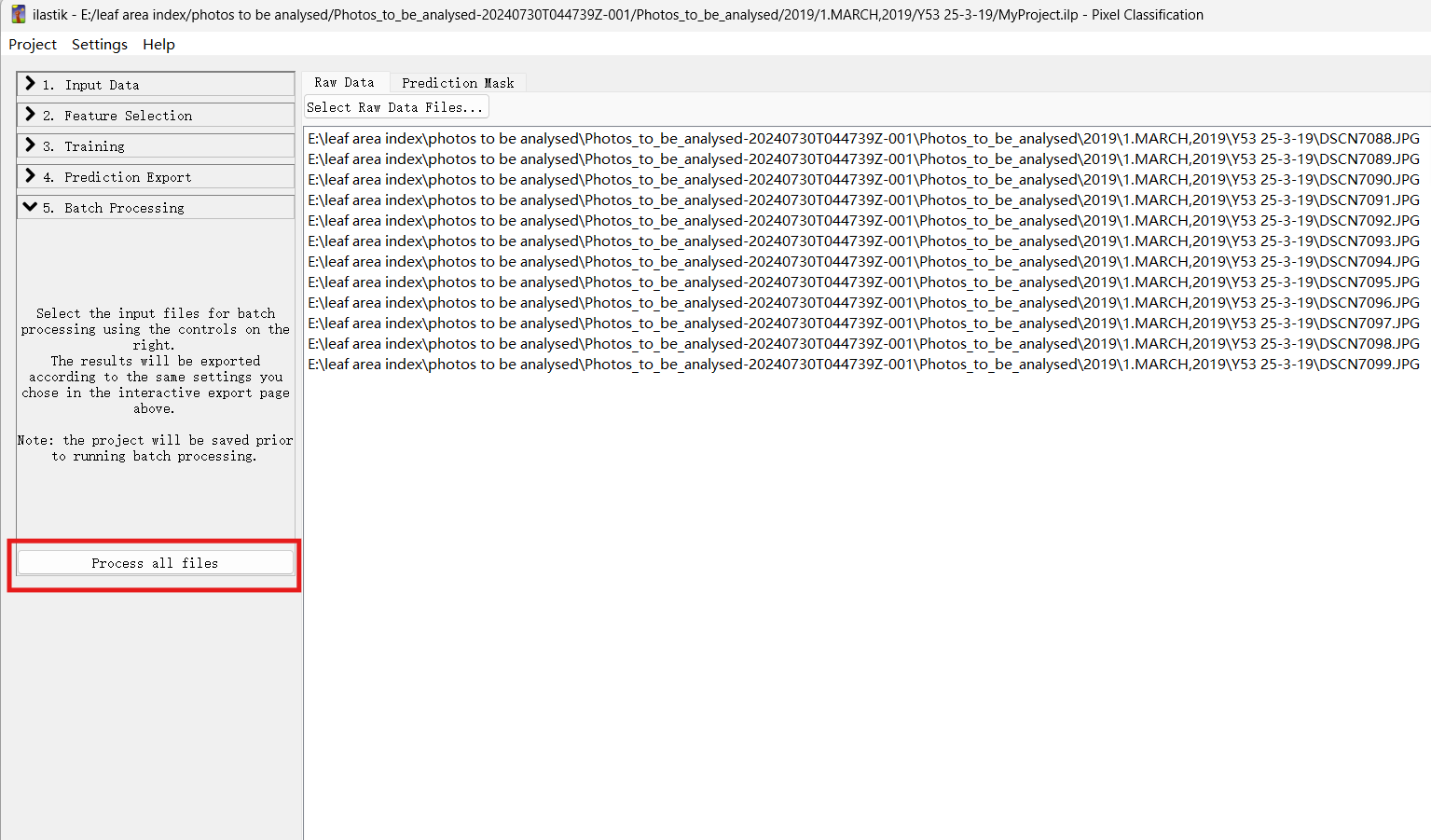

*Figure 31*

Your output files will be in the output folder you set earlier. For me, it is:

C:/Outputs/{nickname}_{result_type}.jpg

Please note that if you save the ilastik project and come back later to batch process more images, **you will have to redo Step 4 (Prediction Export) before Batch Processing, because Step 4 settings are not saved in the .ilp file.**

### Using CAN-EYE to process Binary DHP images that have already been classified using ilastik

First, convert all images into black-and-white, or black-and-grey using the machine learning-based software ilastik (as described in the steps above). Then, organize all the monthly and site-specific processed photos in a dedicated folder for each forest (Figure 32).

*Figure SEQ Figure \* ARABIC 32*

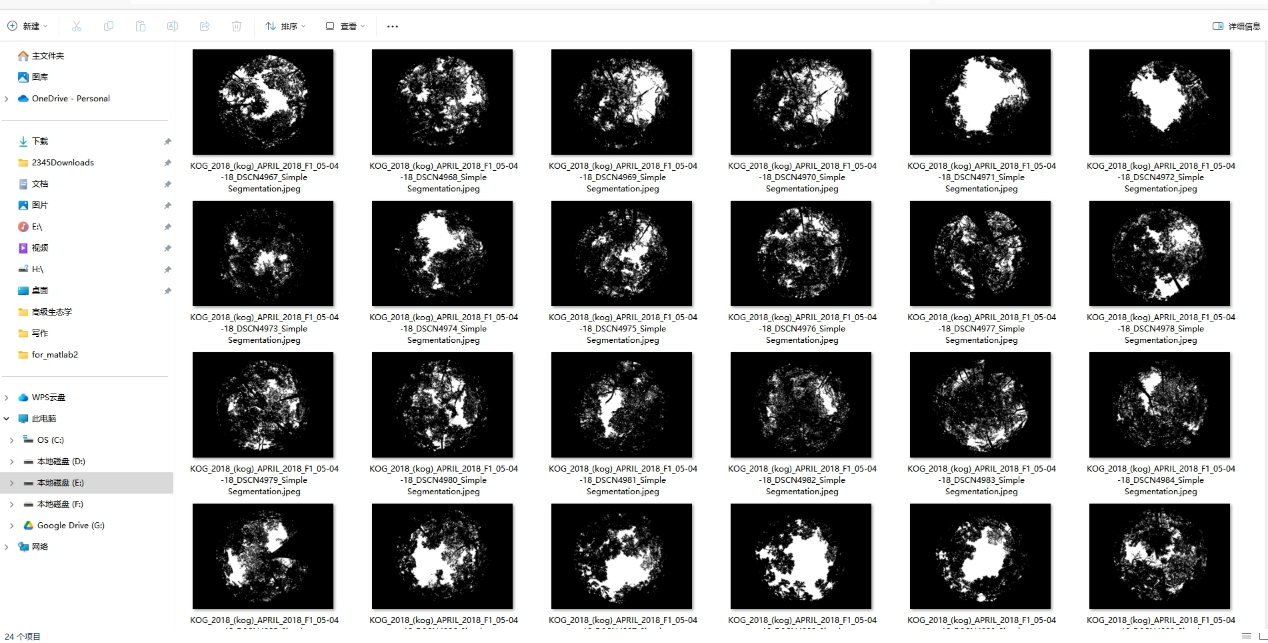

*Figure 32*

There appears to be a bug in CANEYE V6.49. While the protocol states that only three values are possible—with 0 representing vegetation, 100 indicating a gap, 255 denoting a masked pixel—we discovered that CANEYE actually functions differently: grey (100) represents leaves, while black (0) represents gaps (sky). If you use ilastik with category 1 set as sky and category 2 as leaves, and enable 'Renormalize’ to 0-100 option, the output will automatically produce a grey-black image without requiring additional coding. However, if your output is a black-and-white image or entirely black (due to not enabling Renormalize), you will need to convert it to a black-grey format using code. To assist with this, I have uploaded two scripts: **Image_conversion_20220407.m** and **lets_change_values.R**. Additionally, we mask the area outside the fisheye circle with white (255) for visual clarity, though this step is optional since CAN-EYE handles masking later. Below is a comparison of the images before and after conversion.

| 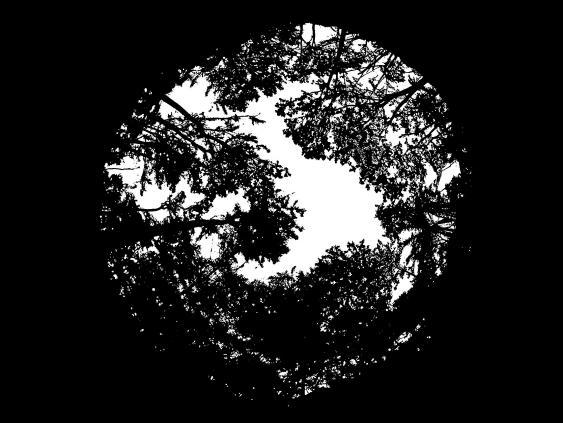 | 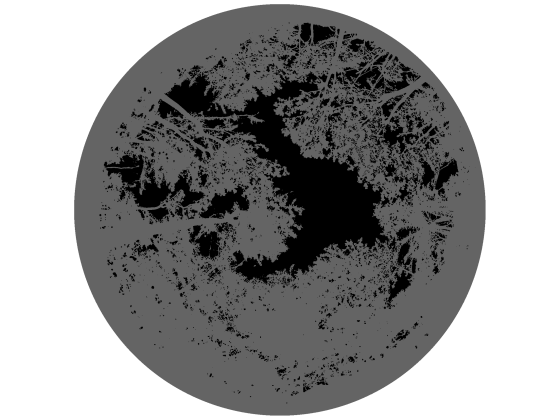 |
| --- | --- |
| Before conversion | After conversion |

Important Note: Only a black (sky) and grey (leaves) image is the correct format for CAN-EYE, which differs from the CAN-EYE manual. I believe there is a bug in the program…

Compress the binary images into a zip folder as required by CAN-EYE (Figure 33). For my field setting, we have 14 one-hectare forest plots. Each plot has 25 image-taking pods. Photos are taken every month. Each zip folder contains 25 images, representing one forest plot for a given month. Each zip folder will yield one LAI value.

According to my experience, I can delete up to 15 images in each zip folder. So, if you find any photos that are problematic (i.e. extreme weather, wrong exposure, obstruction by ferns, camera malfunctions, etc), I would suggest deleting them. I don’t want to include this in the main text without proper testing, but my impression is that, for dense tropical forests, 15 images per plot should be enough. Using only 10 images per plot is definitely insufficient because some of the photos will inevitably fail due to various unavoidable issues, as always.

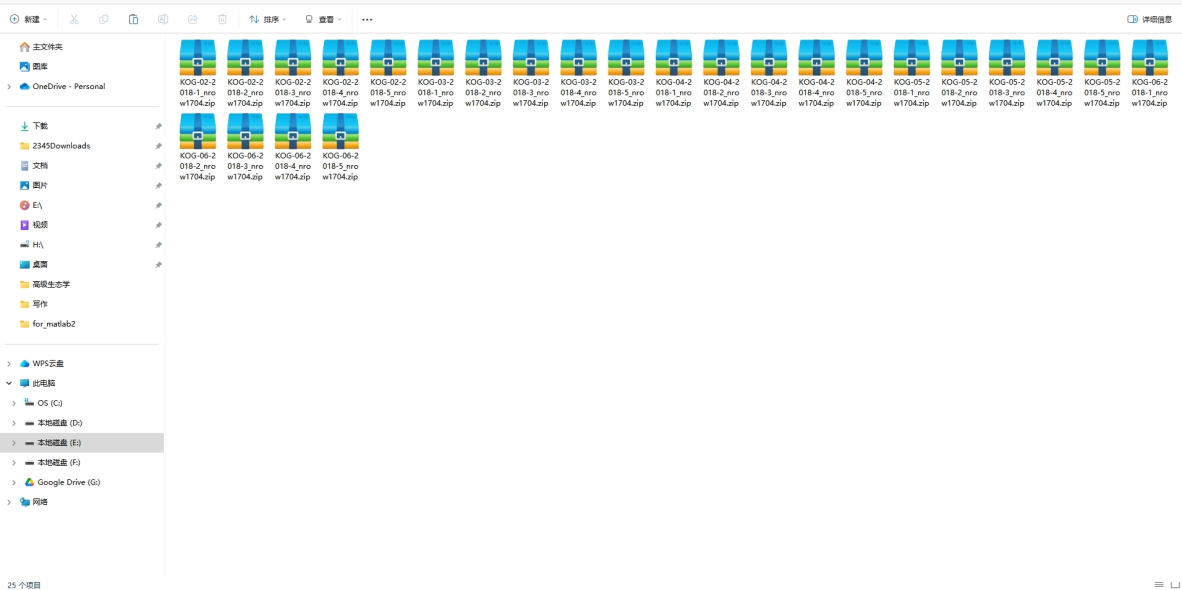

*Figure 33*

If your CAN-EYE application could not be launched from the desktop, try launching it from this location as an administrator: C:\Program Files\UMT_CAPTE\CAN_EYE\application. Note that you will need to install Acrobat (a PDF reader software) before opening CANEYE. The free version of Acrobat works fine.

In CAN-EYE, we select from the menu: DHP > Binary DHP (Figure 34).

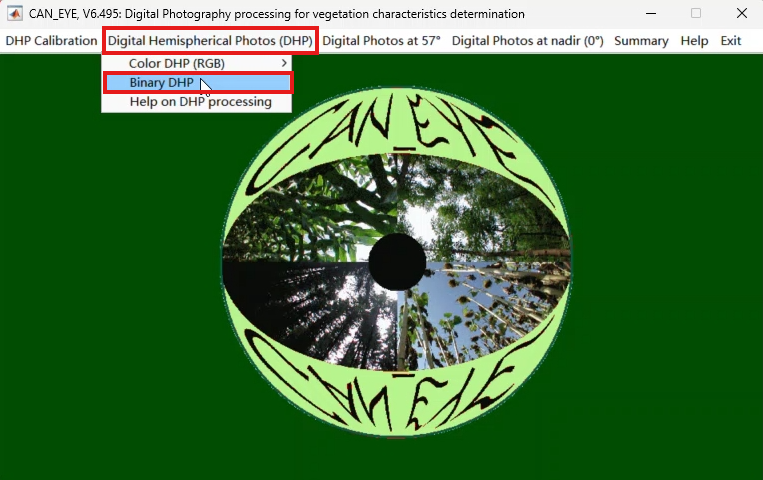

*Figure 34*

Now, in the Excel file that pops up, we fill in the Day of Year (DOY) and latitude, save the Excel File (Figure 35), and then click ‘OK’ (Figure 36). The DOY and latitude values will be used to calculate the total daylight hours for fapar estimates, but they are irrelevant to LAI calculations. However, since Kogyae Strict Nature Reserve (my study site) is very close to the equator, we set the latitude to 5, making DOY unimportant.

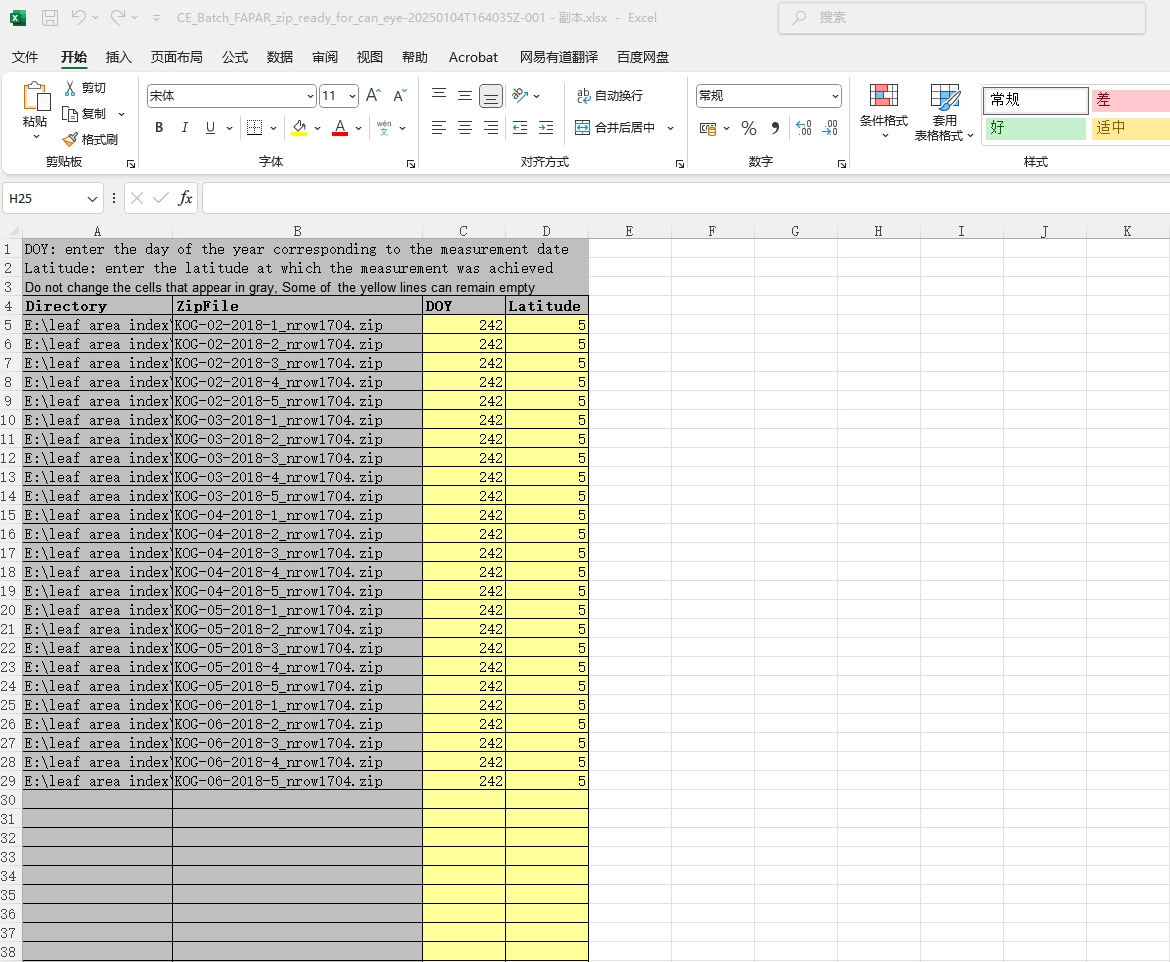

*Figure 35*

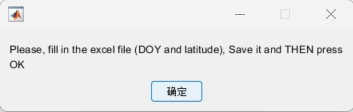

*Figure 36*

In the pop-up 'Processing binary images' window, click 'Yes' (Figure 37). We need to set the relevant parameters.

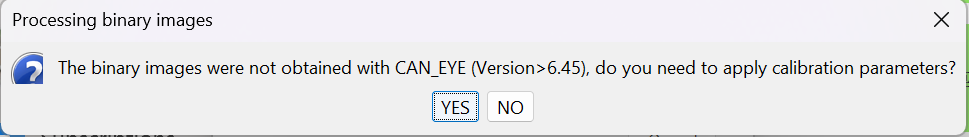

*Figure 37*

Set the following parameters (Figure 38).

(1) Image Size: Row: 1704, Column: 2272, This could be found by right clicking the image and choose properties (Figure 40)
(2) COI (°) = 80, (field of view is 80 degrees; we exclude the edges as they appear slightly fuzzy)
(3) Sub Sample Factor = 1
(4) Fcover (°) = 20 degrees (calculates the percentage of leaf pixels within the central 20-degree ring, indicating canopy openness. Note: This does not include the leaf pixels from the entire image).
(5) PAISat = 10 (When a pixel is completely black, the leaf area index (LAI) becomes mathematically infinite. Since CAN-EYE uses 25 subplot images for LAI estimation, this value addresses cases where all subplot shows black at a pixel. We use 10 for dense tropical forests—based on the assumption that the maximum LAI in such forests in 10—and 6 for savannas).
(6) Latitude and Day of Year (Unclear why the software requests DOY again)

Then, click 'Create' to set the parameters for projection and optical center (Figure 39).

(7) Optical center: Enter line 852 and Column 1136 (the center of the fisheye circle), this is half of the ‘image size’
(8) P1 = 90/809 = 0.11119, where:
    - 90 is the max zenith angle of the fish eye (Note: this example uses a Sigma circular fisheye lens (4.5 mm) with 180 degrees field of view). If you have a 210 degrees field of view fisheye, you will replace 90 with 210/2=105.
    - 809 pixels is the distance from the image center to the edge of the fisheye circle (Figure 41). So at 90 degree zenith angle, the image appear at 809 pixels. You can estimate this by using a ruler on your screen to determine how many % of the fish eye edge is completely black, in our case 5%. So we have distance from the center to the edge as 1702/2 * (100 - 5%) = 809. If you want a precise number, you will need software like QGIS or Photoshop.
    - Press Enter display the line.

Important note: Fisheye lenses are not linear. You could follow CAN-EYE’s more complicated method (via the menu) to derive a curve, but if you prefer simplicity, you can ignore P2 and P3— they have minimal impact on LAI results. Finally, click ‘OK’.

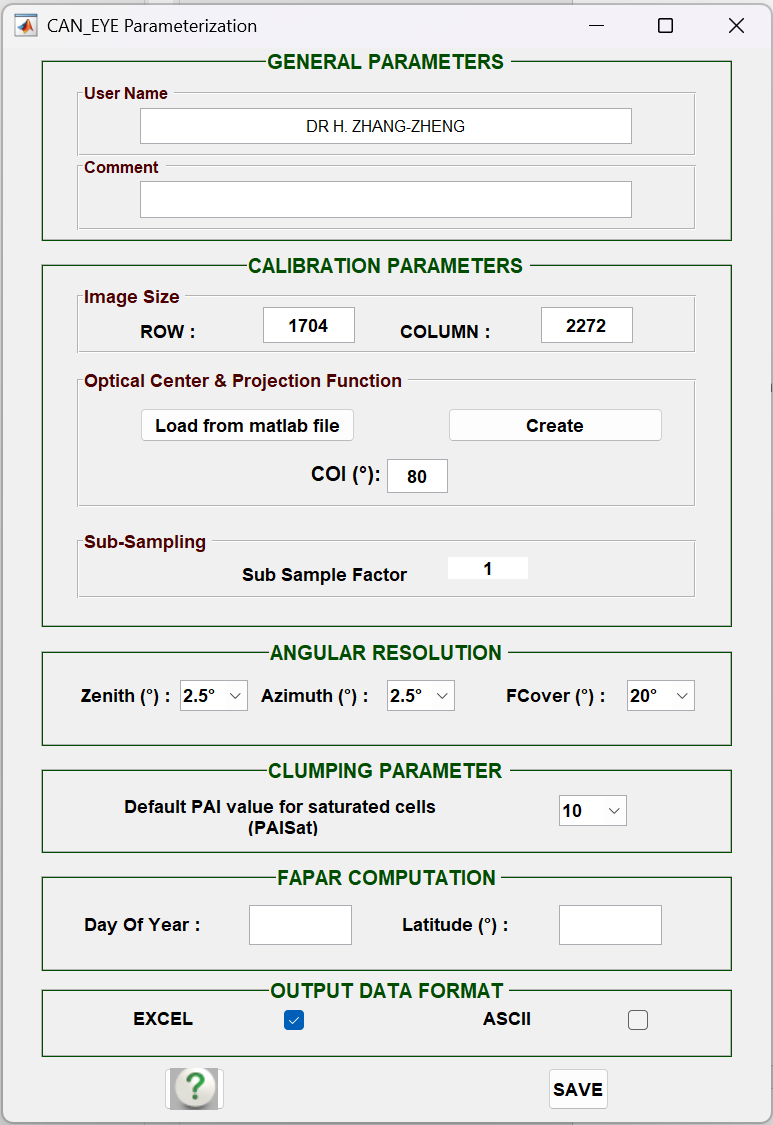

*Figure 38*

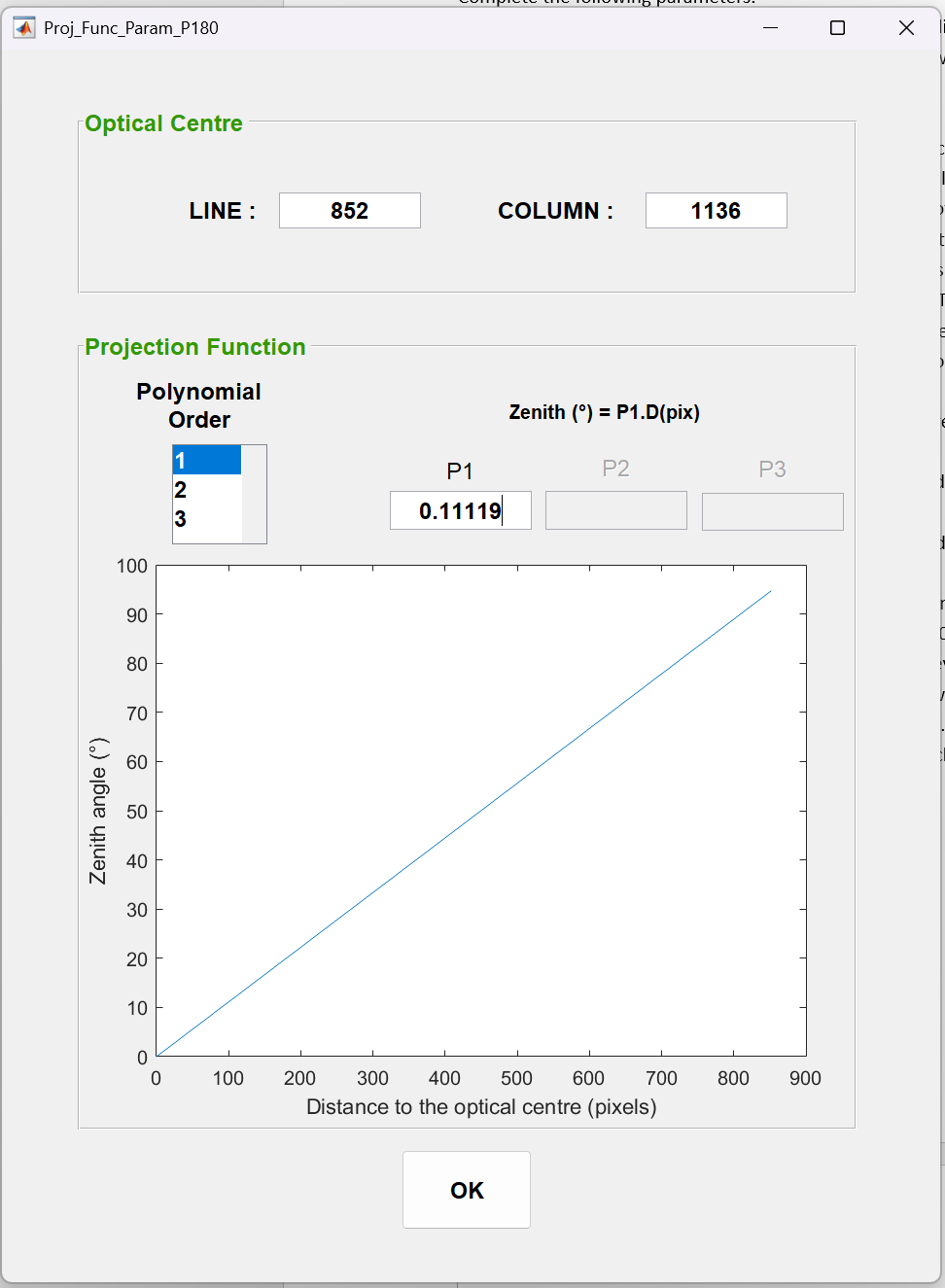

*Figure 39*

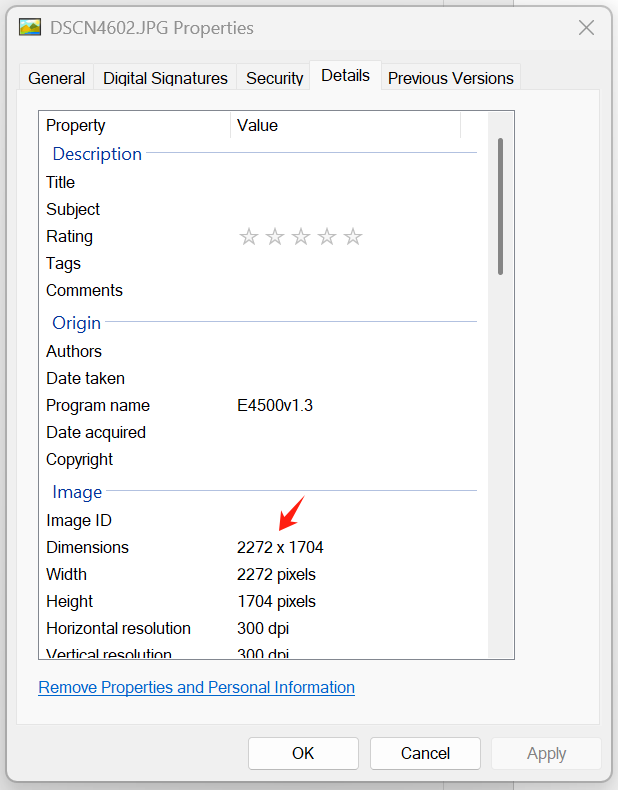

*Figure 40*

**Note: Please do not use the parameter values shown in the screenshots above. You will need to determine your own values for all measurements and calculations (Figure 40 and Figure 41).**

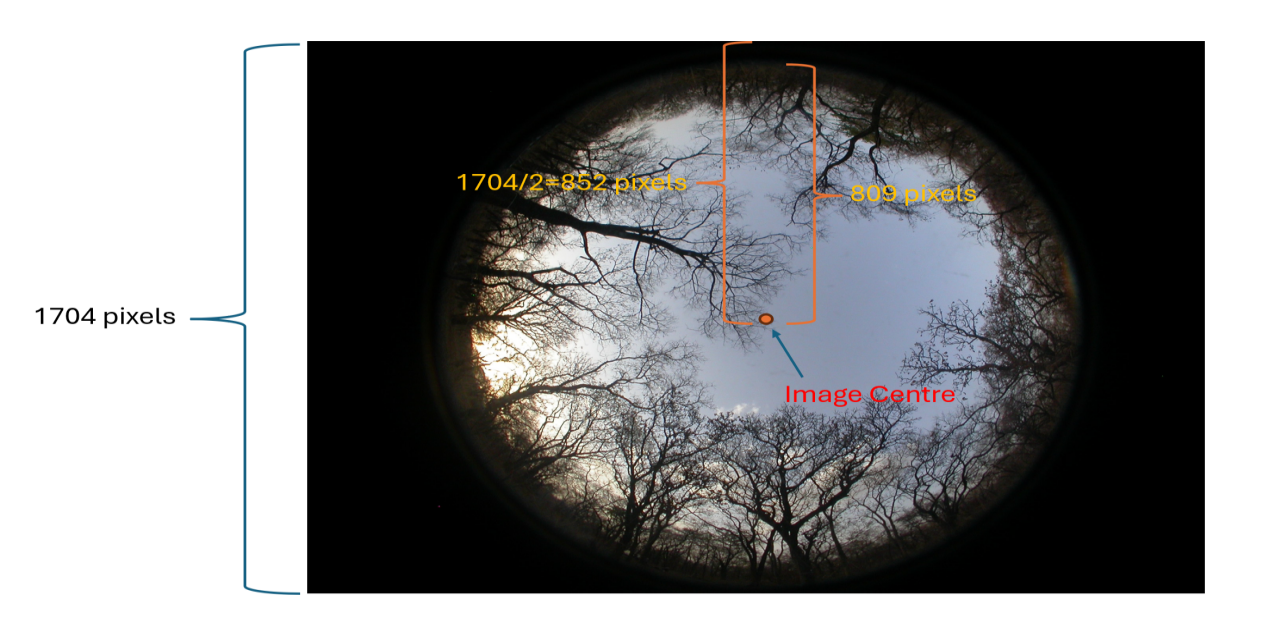

*Figure 41 Number on this screenshot is explained below. 1704 is the height of the image (Figure 40). 852 is the ‘line’ (Figure 39). 809 is the portion of the 852 without the black edge. In our case, ,we estimated it by 852*0.95. It may not be 0.95 for you. This 809 is used to calculate P1 (Figure 39). 809 represents where the horizon is (because this is 90 degree fish eye lense). You will need to address number 90 based on your fisheye lens. You should adjust the 809 depending on your image height and the thickness of the black edge.*

After clicking ‘OK’ and wait for the progress bar to finish loading (Figure 42). Next, save the parameter file, which can be directly applied to the remaining zip packages (Figure 43). By default, the parameter files are saved in the following location: C:\Users\Dr H. Zhang-Zheng\AppData\Local\CAN_EYE\Param_V6

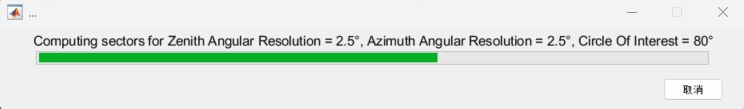

*Figure 42*

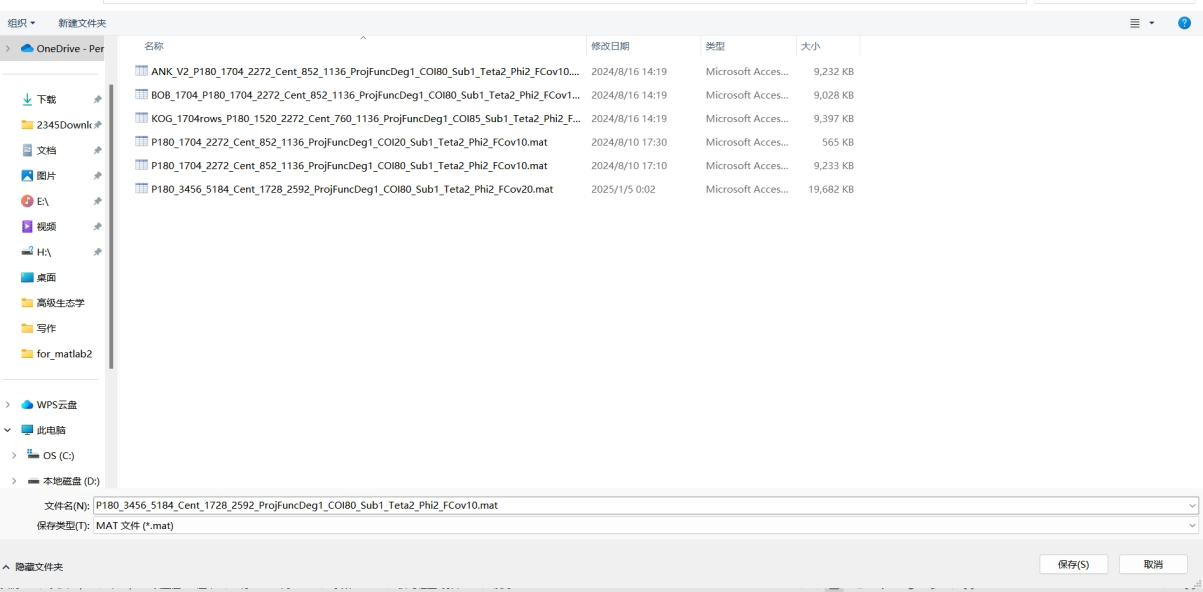

*Figure 43*

Click 'OK' and wait for CAN-EYE to process the first parcel (Figure 44 and Figure 45).

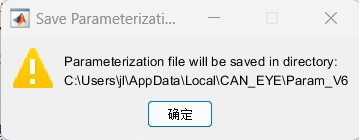

*Figure 44*

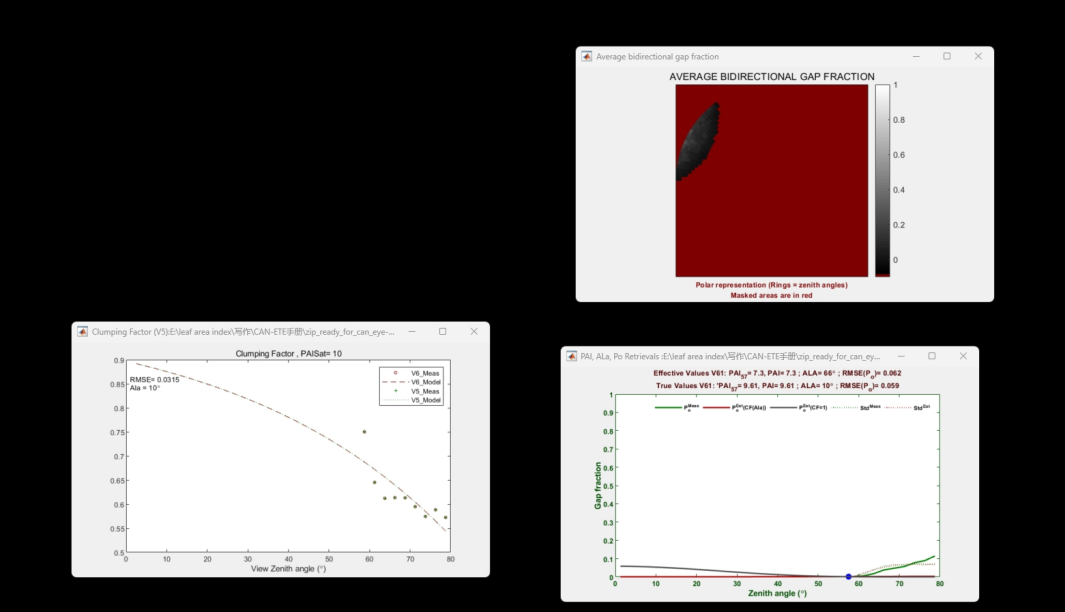

*Figure 45*

Then, in the pop-up 'Processing binary images' window, click 'YES' to process the second zip folder (Figure 46). Select the previously saved parameter file, click 'Load,' and wait for CAN-EYE to process the second parcel (Figure 47).

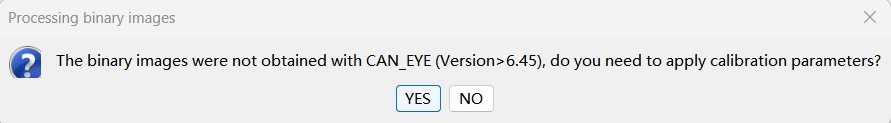

*Figure 46*

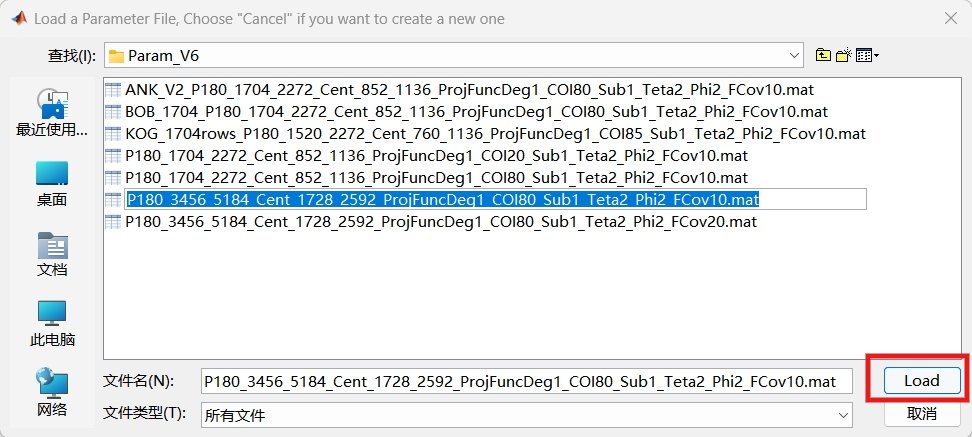

*Figure 47*

Follow the above process to handle all the remaining parcels.

The output from CANEYE (fAPAR and LAI) consists of an Excel file and an HTML file. You can find the LAI values in these files (Figure 49). We have also prepared R code to read multiple Excel files automatically and join these LAI values into one table. Please find **Gather_LAI_fapar_from_caneye.R**

At this step, if you have lots of image packages, you may have to click ‘Load’ and ‘OK’ repeatedly for hours. If you wish to leave your computer unattended to automatically deal with hundreds of packages, you will need a ‘mouse clicker app’ to help you automatically click this button. We used ‘Auto Mouse Click v95.1’, but there are many alternative free software.

### Leaf area index uncertainty

This is the output from CAN-EYE (Figure 48). Note that for each group of images (25 images in our case, representing 25 subplots in each one-hectare forest plot), the results are generated.

58
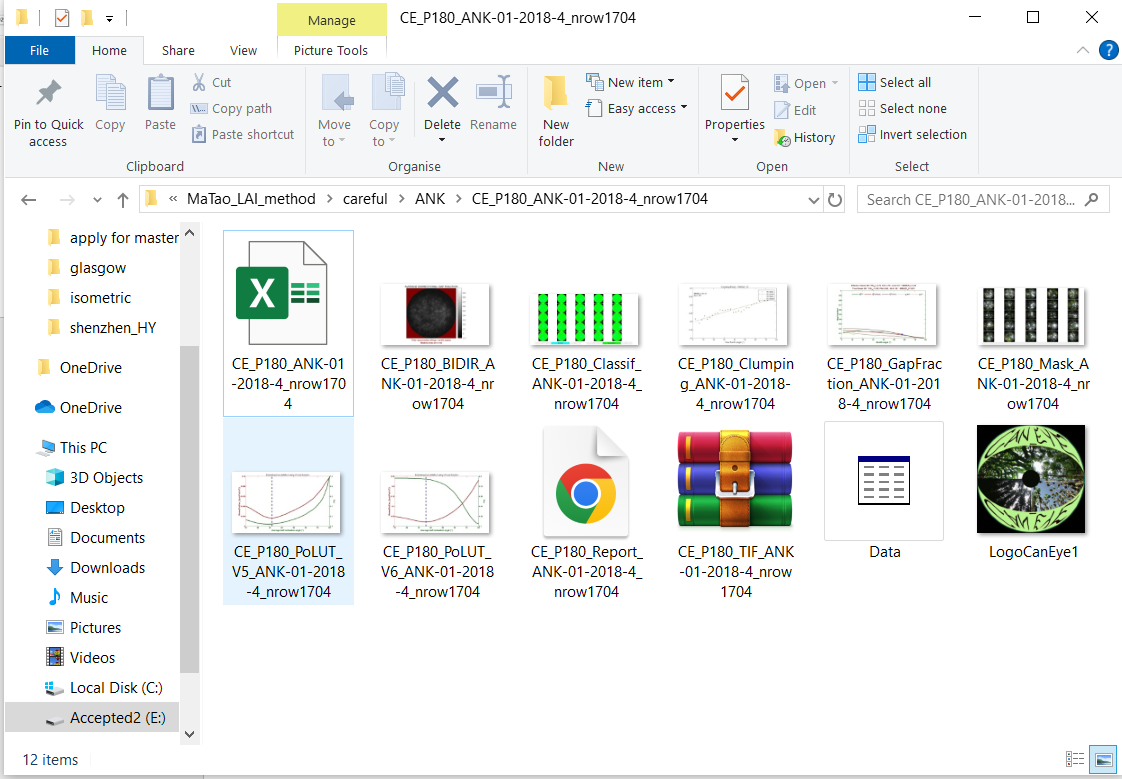

*Figure 48*

The HTML file shows LAI and fAPAR values in a clear way, but we will extract LAI values from the excel file.

5
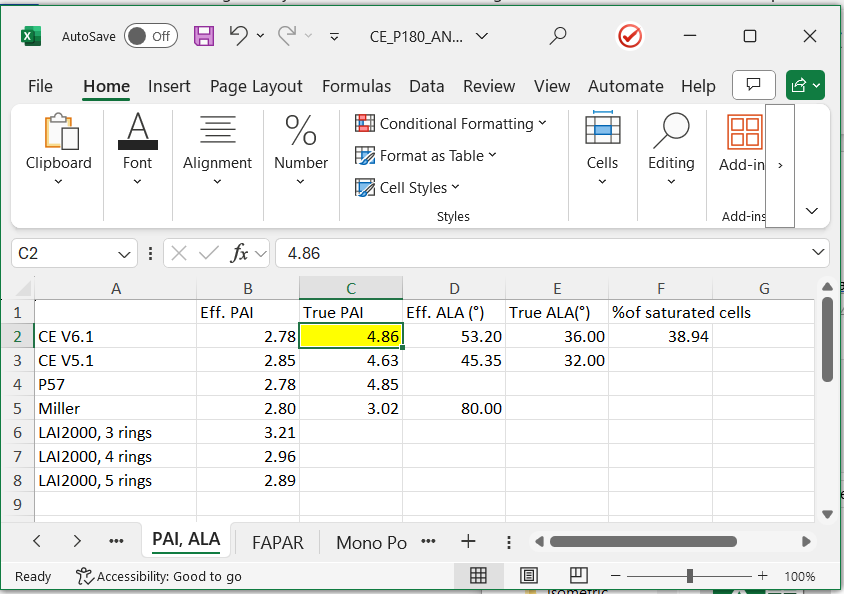

*Figure 49*

We recommend using TRUE PAI under CE V6.1 in tab ‘PAI, ALA’, which is 4.86 (Figure 49). PAI refers to plant area index. This software does not report leaf area index because it believes that all the hemispherical photos could contain stem and branches.

‘True’ means the value has been corrected for clumping. PAI (Plant Area Index) is used because your hemispherical images include branches in addition to leaves. For other values, please check CAN-EYE manual.

You may also want the ‘standard error’ for LAI. Unfortunately, CAN-EYE does not provide an LAI value for each image. Such information is likely impossible for CE V6.1, which combines multiple images together. If you use Hemisfer instead of CANEYE, you can get an LAI for each image.

Nonetheless, we can estimate LAI for each image in CANEYE via an indirect approach. Under the ‘P57 results’ tab, you find P57 values for each image, but not LAI (Figure 50). Note that, as documented in CAN-EYE manual, LAI can be calculated using:

LAI = - LN(P57)/0.93

Then you can obtain LAI for each image following the P57 method. As shown in cell D3, the standard error for P57-based LAI could be calculated (Figure 51). Since we are using CE V6.1, we should scale the standard error from P57 to CE V6.1 (Figure 52), as shown in cell E3. Thus, error_CE_V61 = error_P57 / LAI_P57 * LAI_CE_V61.

Note that ‘25’ refers to 25 images as listed in Column A. ‘4.86’ refers to the CE V6.1 True PAI. You should use your own value instead of these.

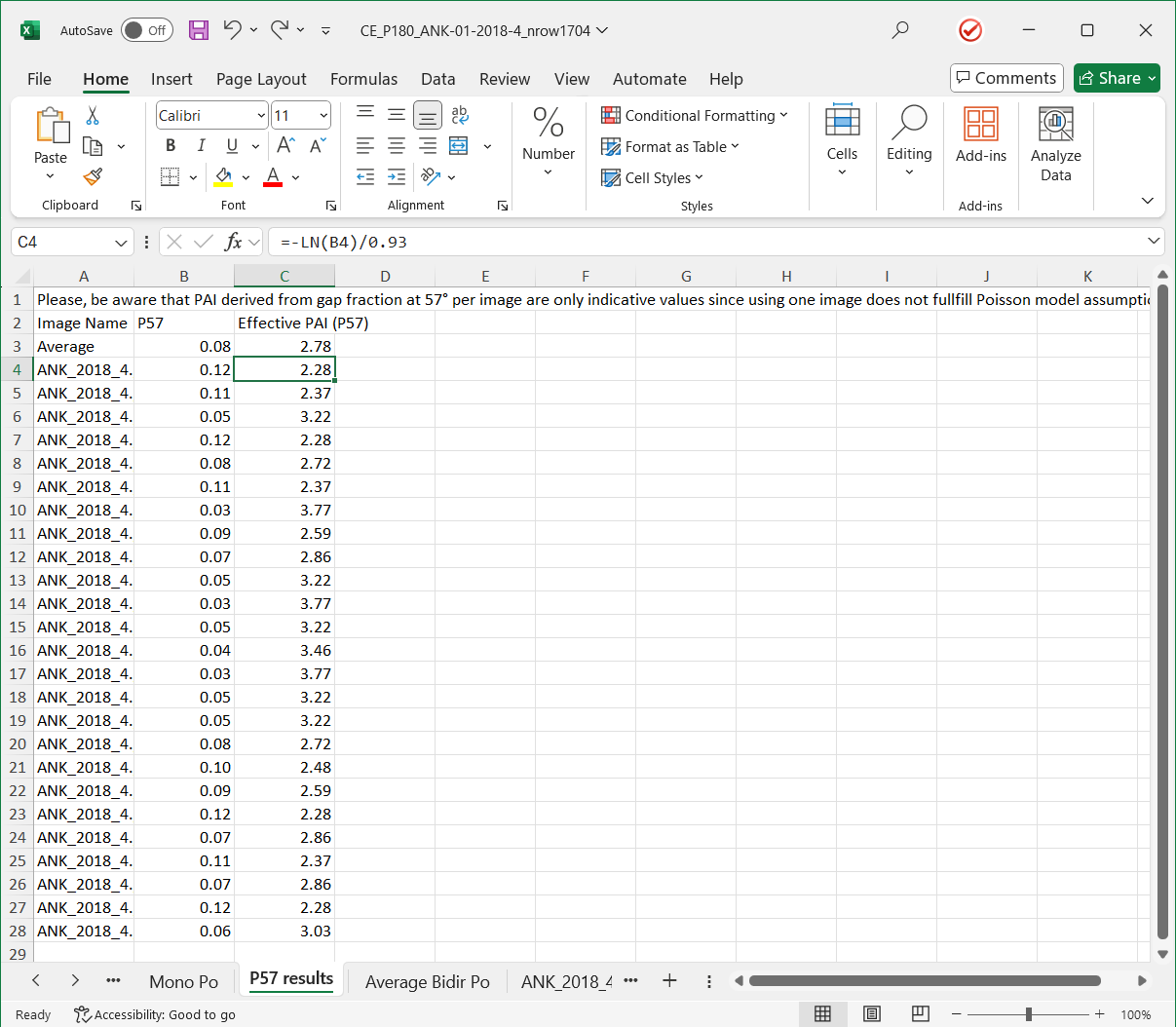

*Figure 50*

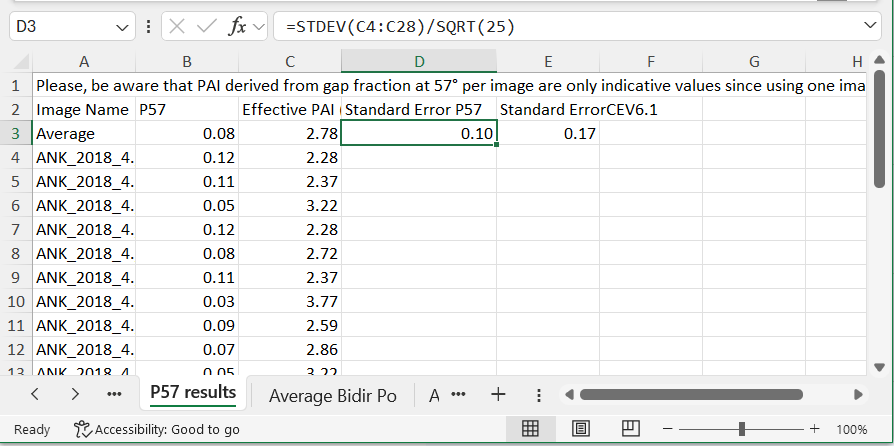

*Figure 51*

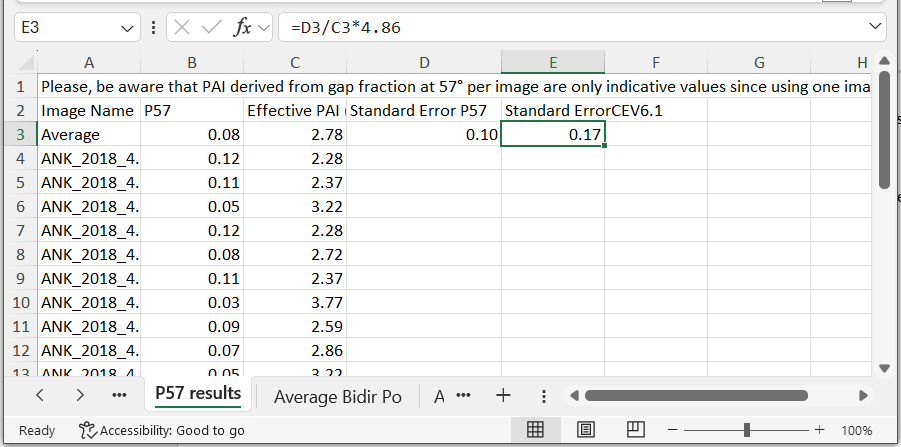

*Figure 52*

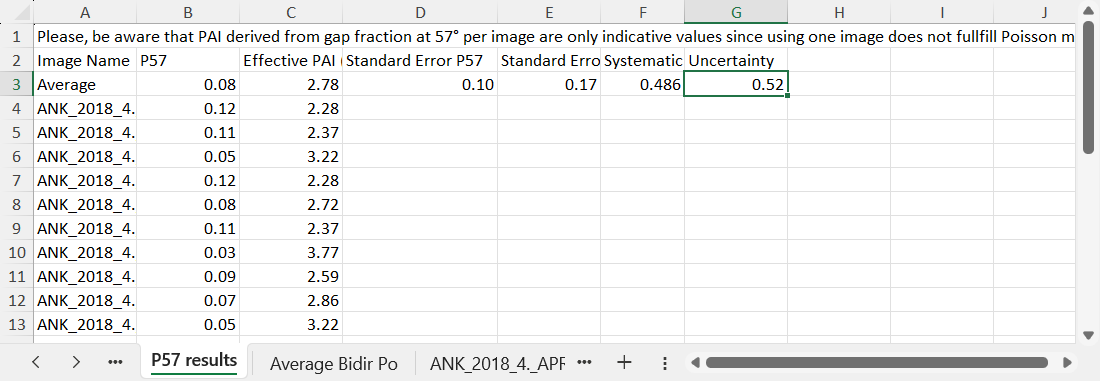

*Figure 53*

The standard error accounts for the random errors of associated with hemispherical photo sampling. We have not accounted for systematic errors, which originate from the algorithm used to convert gap fraction to LAI. Here, we set the systematic error at 10% (Figure 53). Thus, systematic_errors = 0.1 * LAI_CE_V61 = 0.486.

To combine both systematic and random errors (Figure 53), the uncertainty is calculated as = sqrt (systematic^2 + random^2) = sqrt(0.17^2 + 0.486^2) = 0.52.

Note that 0.52 is about 10.7% of the LAI value (4.86). This analysis thus shows that systematic error dominates in the leaf area index calculation, in agreement with the literature (see [this](https://www.sciencedirect.com/science/article/pii/S0168192311001535?casa_token=rjP_yyml13sAAAAA:UzkytpWtOy9KIO_buUdbm-ZqrcimaRDBCpm5AtrI1LT3TXfr1rpCQIl0hVHcDrkw8a4cghU)). Unfortunately, there is no better method for calculating systematic error than an arbitrary estimate.

In short, if you find the above confusing, for simplicity we recommend using an arbitrary 15% uncertainty. For LAI = 4.86, this gives 4.86 ± 4.86*0.15 = 4.86 ± 0.729

### Using hemispheR to compute LAI from ilastik-derived binary hemispherical images

As an alternative to CAN-EYE, ilastik-segmented hemispherical images can be analysed in R with the package hemispheR. Ilastik was configured to produce binary images where sky pixels take one value (255) and all canopy pixels (leaves + wood) take another value (0). These black–white images are imported as single-channel rasters, mapped to hemispheR’s internal convention (canopy = 0, gap = 1),and then converted to LAI from angular gap fractions. The example below shows how to obtain LAI for a single ilastik image using both a LAI-2000–like and a hinge-angle configuration:

\# Example: Computing LAI from an ilastik binary hemispherical image

\# ilastik coding: 0 = canopy (leaves + wood), 255 = sky

**library(hemispheR)**

\# Path to a binary image exported from ilastik

**img_file <- "path/to/ilastik_binary_image.tif"**

\# Import as single-channel fisheye image with a fixed circular mask

**img <- import_fisheye(**

**filename = img_file,**

**channel	= 1,**

**circular = TRUE,**

**circ.mask = list(xc = 1136, yc = 760, rc = 760), # camera-specific**

**gamma	= 1, # no gamma correction (already binary)**

**display	= FALSE,**

**message	= FALSE**

**)**

\# Map ilastik coding to hemispheR convention: canopy = 0, gap = 1

\# sky = 255 becomes gap = 1; canopy = 0 stays 0

**img.bw <- binarize_fisheye(**

**img,**

**method = "Manual",**

**manual = 1,**

**zonal	= FALSE,**

**display = FALSE**

**)**

\## (1) LAI from a LAI-2000–like configuration (0–85°, 2.5° rings and sectors)

**gf_can <- gapfrac_fisheye(**

**img.bw,**

**startVZA = 0,**

**endVZA	= 85,**

**nrings	= 34, # 85 / 2.5**

**nseg	= 144	# 360 / 2.5**

**)**

**lai_can <- canopy_fisheye(gf_can)$L**

\## (2) LAI from a hinge-angle configuration (55–60°, 1 ring, 8 sectors)

**gf_hinge <- gapfrac_fisheye(**

**img.bw,**

**startVZA = 55,**

**endVZA	= 60,**

**nrings	= 1,**

**nseg	= 8**

**)**

**lai_hinge <- canopy_fisheye(gf_hinge)$L**

This procedure can be applied in a loop over all ilastik images within a plot or sampling period.

**Using HemiPy to compute LAI from**

**ilastik-derived binary hemispherical images**

We also analysed ilastik-segmented hemispherical images in Python using the open-source module HemiPy. After ilastik classification, upward-facing digital hemispherical photographs were exported as 8-bit binary images in which canopy pixels (leaves + wood) were coded as 0 and sky pixels as 255. For compatibility with HemiPy, images from each measurement plot were stored in a directory named after the plot and contained in a sub-directory called overstory (e.g. root_dir/plot_a/overstory), following the structure described in the HemiPy documentation.

Binary ilastik images were then passed to hemipy.process, together with pre-computed zenith and azimuth angle arrays based on the camera–lens geometry, the acquisition date, site latitude, and the image direction ("up" for overstory photographs). HemiPy internally applies the Ridler–Calvard clustering algorithm to the input greyscale images; for our 0/255 binary photographs this step simply preserves the ilastik canopy/sky separation. The minimal workflow for one upward-facing plot is:

\# Example: computing LAI from ilastik binary hemispherical images with HemiPy

\# ilastik coding: 0 = canopy (leaves + wood), 255 = sky

**import os**

**import glob**

**import numpy as np**

**import hemipy**

\# Root directory: contains one sub-folder per plot

**root_dir = r"path/to/root_dir"**

\# Example plot: images stored in 'plot_id/overstory'

**plot_id	= "KOG-02-2015-03"**

**layer_dir = os.path.join(root_dir, plot_id, "overstory")**

\# Site information

**lat = 7.0	# site latitude (degrees)**

**date = "2015-03-15"	# acquisition date (YYYY-MM-DD)**

\# Image geometry and radial calibration (camera-specific example)

**img_size = np.array([1704, 2272])	# [height, width]**

**opt_cen = np.array([852, 1136])	# [row, column] optical centre**

**cal_fun = np.array([0.0, 0.0, 0.11119]) # calibration coefficients (^3, ^2, ^1)**

\# Compute zenith and azimuth angle for each pixel

**zenith = hemipy.zenith(img_size, opt_cen, cal_fun)**

**azimuth = hemipy.azimuth(img_size, opt_cen)**

\# Run HemiPy on all ilastik images in the 'overstory' directory

**results = hemipy.process(**

**layer_dir,**

**zenith,**

**azimuth,**

**date	= date,**

**lat	= lat,**

**direction = "up",**

**save_bin_img = False	# ilastik already provides binary images**

**)**

\# Example: extract key canopy variables

**paie_hinge = results["paie_hinge"]**

**pai_hinge = results["pai_hinge"]**

**pai_miller = results["pai_miller"]**

**clumping	= results["clumping_hinge"]**

**fipar	= results["fipar"]**

**fcover	= results["fcover"]**

This procedure can be repeated for each plot by looping over all plot_id/overstory directories within the root folder.
